# Estimating sampling and laboratory capacity for a simulated African swine fever outbreak in the United States

**DOI:** 10.1101/2024.09.26.615206

**Authors:** Jason A. Galvis, Muhammed Y. Satici, Abagael L. Sykes, Kathleen C. O’Hara, Lisa Rochette, David Roberts, Gustavo Machado

## Abstract

The introduction of African swine fever virus (ASFV) into uninfected countries can impact economic and animal welfare. Rapid detection and control of the outbreak contribute to successful eradication and promote business continuity. We developed a model to determine the number of samples, sample collectors, laboratory capacity, and processing times following an ASFV introduction into the U.S. We simulated the spread of ASFV in one densely populated swine region, generating a median of 27 (range = 1-68) outbreaks in 150 days, resulting 616 (range = 1-15,011) sampling events with a total of 3,068 barns (range = 7-69,118) sampled. We calculated the total sample collectors needed, considering daily working hours, sampling and driving time, and laboratory capabilities with and without blood sample pooling. Samples included 31 blood samples and five oral fluid samples per barn, which equal 84,830 (range = 52-2,066,831) and 14,195 (range = 10-345,590) blood and oral fluid samples, respectively. The median number of sample collectors needed to prevent sampling delay varied from 136 to 367 and, in the worst epidemic scenarios, from 833 to 3,115. Notably, excluding downtime–which prevented the sampler from visiting additional farms for 24 or 72 hours–reduced the number of sample collectors needed between 28% and 75%, while switching from blood to oral fluid samples reduced this number between 47% and 75%. At a laboratory processing daily capacity of 1,000 samples, the median days for sample processing without pooling were 92 days, with a maximum of 5.7 years. We demonstrated a need to redistribute 10,062 (range = 2-67,940) unprocessed samples daily to other laboratories to prevent processing delays. Our study addresses the challenge of efficiently organizing resources for managing a potential ASFV outbreak, providing information about the number of sample collectors and laboratory capacity needed for one densely populated swine region in the U.S.

## Introduction

African swine fever virus (ASFV) is a highly contagious hemorrhagic viral disease affecting domestic swine and feral and wild pigs (Blome et al., 2020). It has disseminated worldwide, spreading widely through Europe (Cwynar et al., 2019), Asia (Mighell and Ward, 2021) and Africa (Njau et al., 2021), and has more recently been identified in the island of Hispaniola in the Americas (Jean-Pierre et al., 2022; Schambow et al., 2023). The potential introduction of ASFV into the U.S. swine industry presents a significant challenge that threatens not only animal health but also the national economic stability and food security (Sánchez-Cordón et al., 2018; Dixon et al., 2019). Consequently, robust surveillance and early detection mechanisms have become paramount to prevent catastrophic losses and ensure business continuity of the swine industry (Sykes et al., 2023; Checkoff, 2023; Carriquiry et al., 2020). A critical component of a national response plan includes efficient collection, processing, and analysis of samples to promptly detect infected premises, and demonstrate ASFV effective control measures and disease elimination (Gallardo et al., 2019; Schambow et al., 2022).

In the U.S., swine veterinarians oversee or directly handle the collection of samples to detect and control endemic pathogens. However, in the event of an ASFV epidemic, the number of swine industry veterinarians and animal health officials may be insufficient given the large number of samples needed to surveil for disease (Secure Pork Supply Plan, 2024). To address this challenge, a collaborative initiative involving the swine industry, government officials, and academic experts has developed a certified swine sample collector training program (Secure Pork Supply Plan, 2024). This program aims to train on-farm personnel, such as animal caretakers, to collect, package, and submit samples. By increasing the number of qualified personnel who can submit high-quality samples, this program is expected to enhance the efficiency and effectiveness of the response plan to control ASFV spread. Despite this unique sample collector program, the process of sample collection is intricate, requiring a well-coordinated effort that hinges on the availability of sufficient manpower to collect a large volume of samples (USDA, 2023; Alvarez et al., 2023). Insufficient staffing can lead to delays in sample collection and, consequently, delays in the detection and response (Sykes et al., 2023; Hayes et al., 2021).

The volume and type of samples collected are critical factors when estimating the number of sample collectors and time involved in ASFV response. The U.S. ASFV response plan includes blood swab samples from suspected farms for initial investigation (USDA, 2023). Furthermore, these plans also have established that farms within infected, buffer, and surveillance zones, and those directly linked with ASFV-positive premises via recent animal shipments or vehicle movements need to be tested (USDA, 2023; Sykes et al., 2023). Although this national response plan established an ASFV sampling scheme, the number of samples, sample collectors, time, and resources necessary to collect and process these samples during an ASFV incursion could vary and have yet to be estimated. This gap in our understanding highlights opportunities to evaluate additional information to support planning to effectively control an ASFV outbreak.

The ongoing outbreaks of ASFV in Europe after its reintroduction in 2007 have underscored the critical importance of timely detection to prevent a foreign animal disease from becoming endemic within new territories (Cwynar et al., 2019). To prepare the U.S. swine industry, a comprehensive understanding of the logistics involved in sample collection and processing is needed to optimize the strategic planning and implementation of ASFV response strategies (Cochran et al., 2023). This work aimed to evaluate the logistical considerations of sampling during a simulated ASFV outbreak in the U.S., calculating the total and daily number of blood samples, and an alternative scenario with oral fluids, the necessary personnel to collect these samples, the time from collecting to processing samples, and the laboratory capacity needed.

## Methodology

### Data

This study used data from 1,898 farms from eight commercial swine companies in one U.S. state. The data comprises farm identification, production type (e.g., wean-to-finisher), number of pigs per farm, latitude, and longitude. For each farm, we also collected the lines of separations (LOS), which is part of enhanced on-farm Secure Pork Supply (SPS) biosecurity plans, to identify the number of barns within each farm (Center for Food Security and Public Health, 2017). These data were collected from the companies through the Rapid Access Biosecurity application (RABapp^TM^) (https://machado-lab.github.io/rabapp/), a web application that serves as a platform for standardizing the approval of SPS biosecurity plans while storing and analyzing animal movement data (Machado et al., 2023). In this study, 9.5% of farms were not in RABapp^TM^, missing essential LOS data to identify the number of barns. We estimated the number of barns for these farms based on the average number of barns from the remaining 90.5% farms with LOS information. In addition, we collected addresses, latitudes, and longitudes from 27 sample supply offices of the participant companies and one certified laboratory from the National Animal Health Laboratory Network (NAHLN) to process ASFV samples from all swine commercial companies. This laboratory reported the capacity to process 1,000 ASFV samples daily through polymerase chain reaction (PCR) tests, following the ASFV laboratory guidelines (USDA, 2023).

The commercial swine companies also provided pig and vehicle movements from January 1st, 2020, to December 31st, 2020. Animal movement data included the identification number of farms sending and receiving pigs, the date, and the number of pigs moved. Vehicle movement was shared by two companies that represented 97% of the farms within the region, this data comprehended geographic coordinates every five seconds, speed (km/h), date, and time for each vehicle. The companies provided a list of 599 vehicles; these vehicles include (i) 224 trucks used to deliver feed to farms; (ii) 168 vehicles utilized in the transportation of live pigs between farms; (iii) 125 vehicles used in the transportation of pigs to markets (a.k.a. slaughterhouse, packing plants); and (iv) 82 vehicles without a defined role, which are used for multiple tasks such as delivering feed and pigs. Finally, we also gathered information on 14 company-owned cleaning stations where vehicles are regularly cleaned; more details about the vehicle data and contact network are presented elsewhere (Galvis and Machado, 2024a).

### African swine fever outbreak simulation

We simulated the ASFV epidemic through a stochastic, farm-level, compartmental transmission model with four health states: Susceptible (S), Exposed (E), Infected (I), and Detected (D), available in PigSpread-ASF (Sykes et al., 2023). This model included six transmission routes: i) movement of pigs between farms; ii) local transmission, reflecting transmission related to spatial proximity; iii) vehicles moving pigs between farms (pig trucks); iv) vehicles moving pigs from farms to slaughterhouses (market trucks); v) vehicles delivering feed to farms (feed trucks); and vi) vehicle movements between farms without a defined role (undefined trucks) (Galvis and Machado, 2024a). The pig movement network was reconstructed using the origin and destination farms of the pigs moved daily, and the vehicle contact network was reconstructed by a methodology that used the proximity between vehicles and farms to identify a vehicle contacting a farm while considering the effect of disinfection when vehicles are at a cleaning station (Galvis and Machado, 2024a). Ultimately, ASFV dissemination via pig and vehicle is driven by directed temporal networks. The local transmission was based on a kernel density where the probability of infection decreases with increased distance. Initial simulation conditions included ASFV infection seeded in a random farm for each simulation, and transmission was allowed from day two. 94,900 simulations were run, which included seeding infection in each of the farms 50 times. The ASFV model simulated all control actions listed in the U.S. response plan (USDA, 2023), including i) depopulation of ASF-positive farms; ii) a 72-hour standstill of live pig movements; iii) contact tracing of farms connected to ASF positive cases by animal and vehicle movements; and iv) the implementation of control areas (a 3 km infected zone and a 2 km buffer zone) and surveillance zones (5 km). The model calculated the number of diagnostic tests required for both surveillance purposes and movements to/from farms in the control areas, hereafter referred to as pre-permit testing (Sykes et al., 2023). Given the lack of specific USDA guidelines regarding the required number of individual blood swab samples for ASFV-suspect farms, as well as details on sampling frequency and duration (USDA, 2023), we adopted a strategy to collect 31 blood samples per barn based on the number required for pre-movement permits according to current guidelines (USDA, 2023). This approach also utilizes the sampling frequency and duration parameters from the USDA’s 2020 guidelines (USDA, 2020), as detailed in Supplementary Material Table S1. Additionally, we evaluated the use of oral fluid sampling. Even though oral fluids are a sample type not approved for ASFV testing by USDA policies, oral fluids could eventually be approved to serve as a complementary sample type alongside blood samples in future protocols (Goonewardene et al., 2021a). For this scenario, we assumed that oral fluid samples would be collected from five different pens per barn. Of note, we calculated logistics, sample collection, and testing for oral fluid based on discussions with the swine industry and state animal health officials (personal communication). Therefore, the results from the oral fluid should be considered as an initial exploration of its potential future adoption in some use cases. Our ASFV model calculated the samples required for all simulated outbreaks for up to 150 days.

### Sample collectors and laboratory capacity estimation

#### Sampler collector route

Sample collectors from each company are permanently assigned to pick up supplies at specific office locations, where they initiate their journey, which includes collecting the necessary sampling material (i.e., tube, needles) and driving to an assigned farm to perform the required sampling (Figure 1.) Once the sampler arrives at the farm, sampling is initiated immediately; the average time for each blood swab sample to be drawn was assumed to be five minutes. For our oral fluid alternative sample scenario, the collector used ropes hung at pens, and the simulated time for the five samples collection was 25 minutes (Goonewardene et al., 2021b; Mur et al., 2013). After the sampling, sample collectors transported the collected samples to the accredited laboratory. The sample collector’s journey path between supply office*→* farm*→* laboratory is calculated in our model; after calculating all possible pathways, the shortest driving route between these locations is used in the final simulation

**Figure 1.**
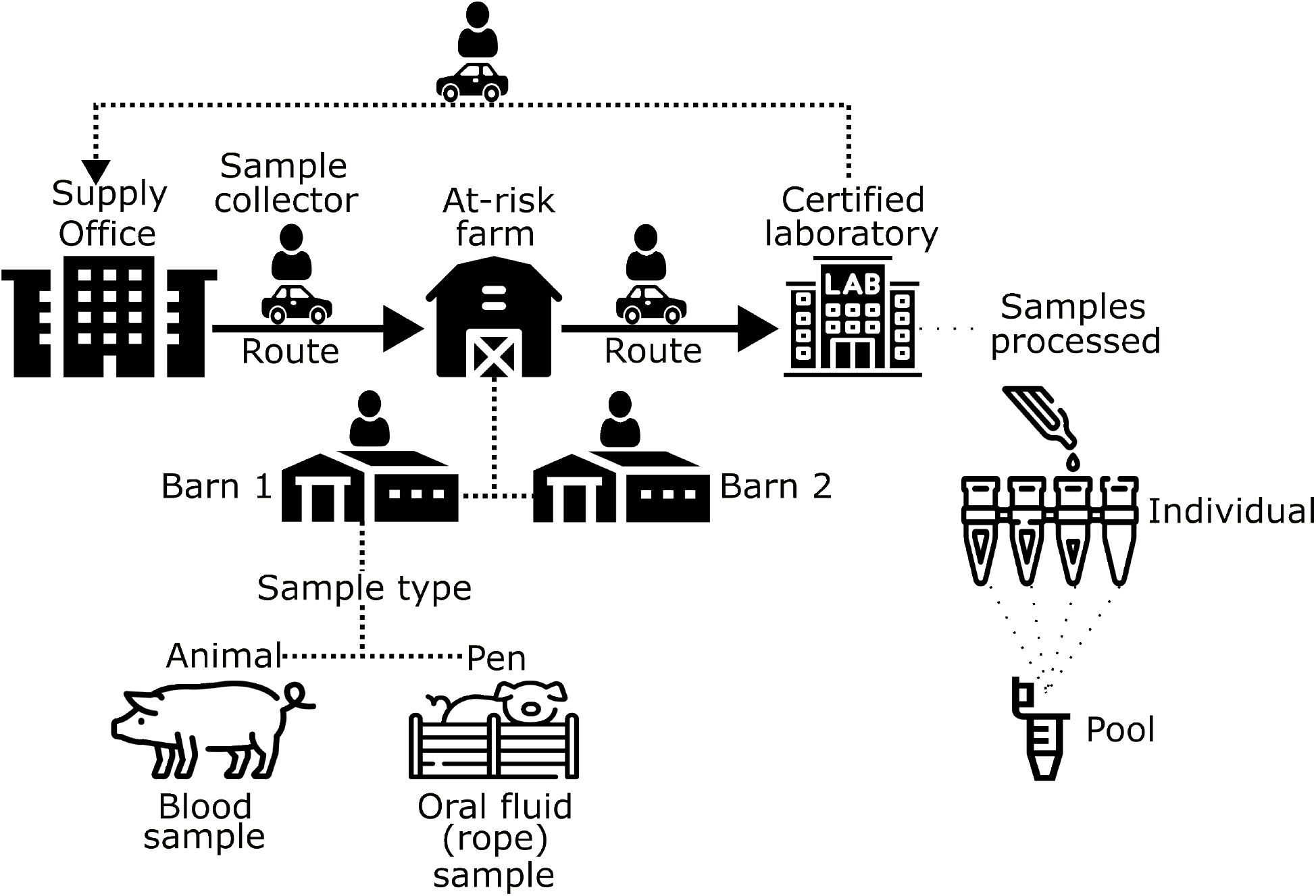
ASFV sampling flow. The diagram shows the order of events a sample collector follows to collect samples from farms and deliver them to the laboratory

(Figure 1). We used Python’s package OSRM for route calculation (Luxen and Vetter, 2011).

We simulated the dispatch of sample collectors to farms that required sampling based on the ASFV national response plan sample requirements (USDA, 2023). We refer to those farms as “at-risk” farms. We also defined the order of sampling as follows: 1) farms that recorded animal visits from infected premises within the last 30 days; 2) vehicle visits from infected premises within the last 15 days; 3) Pre-movement permit testing from farms within infected and buffer zones three and one day before movement; 4) farms in infected zone(s); 5) farms in buffer zone(s); 6) farms in surveillance zone(s). In addition, farms with breeding age animals, a.k.a. sow farms, farrow-to-finisher, and boar studs, were a priority for sampling, followed by nursery, wean-to-finisher, and finisher farms (Figure 2.)

**Figure 2.**
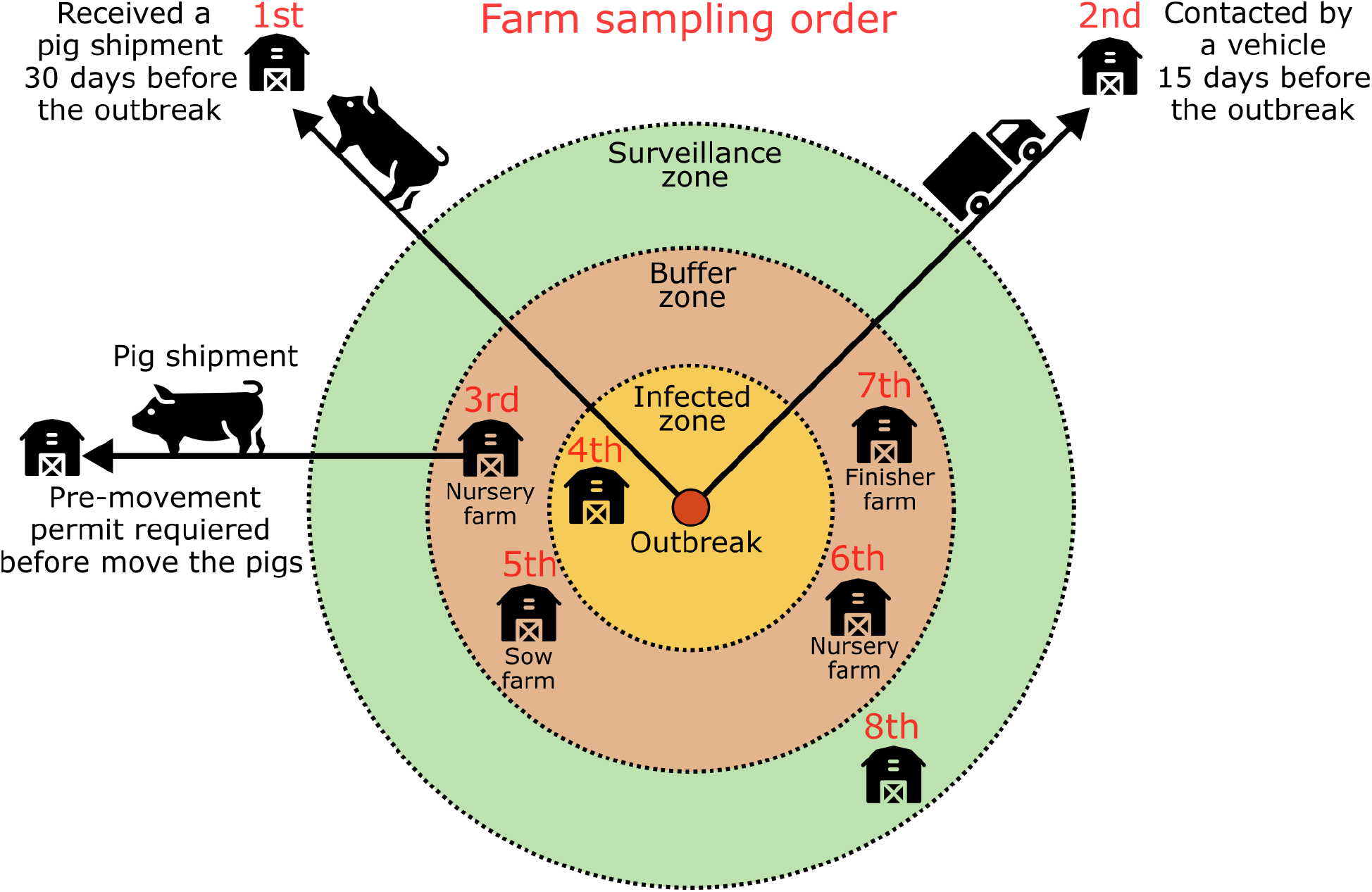
Farm sampling order for a group of eight farms. The first priority was given to the farm that received pigs from the ASFV-positive farm. The second priority was the farm that has been in contact with vehicles exposed to an ASFV-positive farm. The third priority was the farm requiring a pre-movement permit. Priorities four through eight involved farms within the control zones, where farm types are prioritized hierarchically. Specifically, the fourth priority was a farm within the infected zone, the fifth was the farm with the highest strategic value within the buffer zone, notably a sow farm, followed by a nursery farm, ranked sixth, and a finisher farm at seventh. The eighth priority was assigned to a farm within the surveillance zone.

#### Sample collectors allocation

Our methodology utilized a Greedy Algorithm to generate solutions that satisfy the constraints of the situation and estimate the total number of sample collectors needed to finish the sampling process while keeping the sampling time reasonably low (Jungnickel, 1999; Hernando et al., 2018). Figure 3 displays the overall flow of the sample collector allocation algorithm and a pseudo-code is available in Supplementary Material Appendices A.

**Figure 3.**
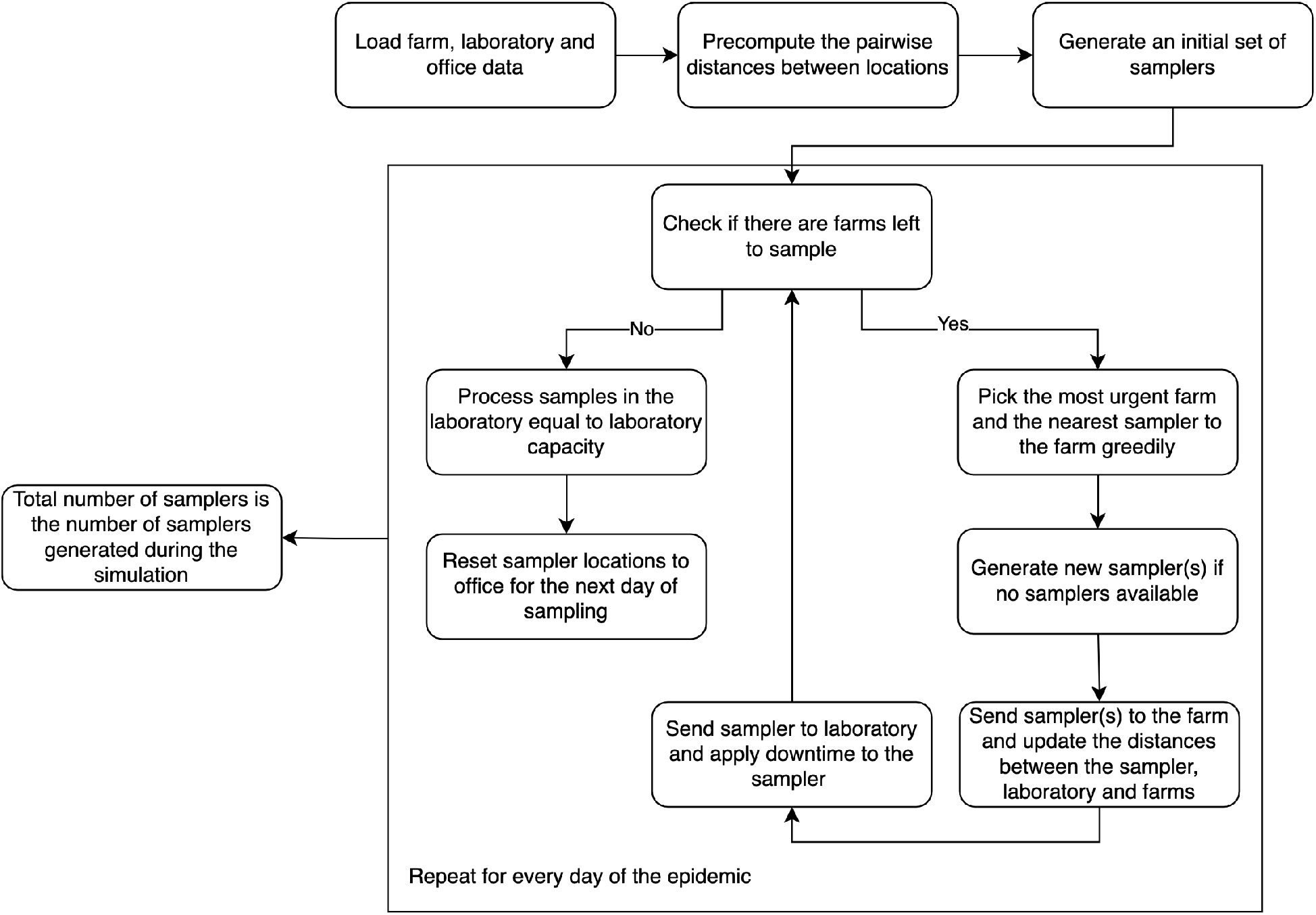
Flow chart of the sample collector allocation. The diagram describes the steps from the sample(s) required until the sample(s) is tested at the laboratory

Our algorithm incorporates 17 conditions to estimate the number of sample collectors (Table 1.) It operates under the assumption that farm personnel will aid in the sampling process by restraining pigs. Additionally, several sample collectors were trained and certified and are available to sample any farm within the study region. However, once a sample collector is assigned to an office from a specific company, that sample collector is permanently assigned to that location. We have implemented an allocation cap to prevent any single company from monopolizing all available sample collectors. This cap is proportional to the company’s farms within the region. For example, if company A owns 90% of farms, it can only claim up to 90% of the available number of sampler collectors. Furthermore, the algorithm dictates that the number of samples required at a farm directly correlates with its number of barns (Figure 1), meaning larger farms with more barns will need more samples (USDA, 2023). If a single sample collector cannot complete the sampling required of the farm they have been dispatched to within one day due to their working schedule, the farm receives additional sample collectors. To prevent large farms from overwhelming the system, a cap on the maximum number of collectors per farm has been set (Table 1). In instances where no extra sample collectors can be allocated (e.g., all sample collectors are dispatched to farms, downtime restraints are placed on sample collectors), the algorithm permits existing collectors to extend their working hours (Table 1.)

**Table 1.** Conditions to estimate the number of sample collectors and laboratory capacity.

| Condition | Description | Study parameters |
| --- | --- | --- |
| Trained sample collectors (TSC) | Sum of trained personnel to collect ASFV samples from all companies in the region | 500, 1,000, 1,250, 1,500, 1,750, 2,000, and no limit |
| Sample collectors per farm (SCPF) | Upper limit on the number of collectors that can be assigned to a single farm | 3 and 5 |
| Regular working schedule (RWS) | Regular number of hours a sample collector can work per day | 8 hours and 12 hours |
| Exceptional working schedule | Maximum number of hours a sample collector can work per day | 23 hours |
| Collected sample type | Sample methods at the farm | Blood and oral fluids |
| Collected sample size | Number of samples collected from a barn | 31 blood samples and five oral fluids samples |
| Farm's barns | Number of barns within a farm. | Estimated per farm through on-farm biosecurity plans |
| Total farm samples | Number of samples that need to be collected from an at-risk farm | Collected sample size multiplied by Farm's barns |
| Time per sampling | Number of minutes a sample collector needs to collect one sample at the farm | Five minutes per blood sample and 25 minutes per barn with oral fluid samples |
| Time to farm | Number of minutes sample collector takes to go from the office to the farm | Estimated by driving time |
| Time to sample | Number of minutes sample collector takes to collect the samples from the farm | Time per sampling multiplied by Total farm samples |
| Time to laboratory | Number of minutes sample collector takes to go from the farm to the laboratory | Estimated by driving time |
| Total sampling time | Number of minutes required for a sample collector to collect and deliver the samplers to the laboratory | Sum of Time to farm, Time to sample and Time to laboratory |
| Downtime | Number of hours a sample collector is unavailable after finishing the sampling on a farm | 0 hours, 24 hours, and 72 hours |
| Sample pool size | Number of samples combined from a single farm for processing at the laboratory | 1, 5, and 10 |
| Laboratory capacity | Daily number of samples the laboratory can process | 1,000, 2,500, and 5,000 |
| Laboratory storage | Total number of samples the laboratory can store | Not limited |

At the beginning of the simulated ASFV epidemic, the algorithm assigned one sample collector to each office (Figure 3), prioritizing the assignment of collectors to farms according to the relationship with the infected farms (e.g., zones) and farm type (Figure 2.) However, if the number of collectors proves insufficient to cover all farms, the algorithm deploys an additional collector to the office that can achieve the new sampling collection in the shortest time. In scenarios where downtime is minimal (e.g., no downtime), a collector might be capable of visiting several farms within a single day, depending on their working schedule (e.g., eight hours/day). However, if all trained sample collectors are either actively sampling or in their downtime period, the algorithm postpones further sampling activities. For instance, if 500 sample collectors exist, all of whom have already finished their working schedule for the current day, then the sampling of the remaining farms will resume the next day. The algorithm then seeks the next available collector who can most efficiently fulfill the pending sampling requests. Indeed, our approach ensures flexibility by allowing the assigned group of sample collectors to initiate their tasks at different times at a designated farm, thereby minimizing delays in the sampling process. The algorithm continues to reallocate sample collectors until sampling is completed across all farms. After the sampling collection at a farm is finished, each collector is responsible for delivering the samples to the laboratory before returning to their respective offices for a required downtime period (Figure 1).

#### Sample processing

Upon arrival at the laboratory, samples are stored and labeled based on the testing priority order of the originating farms (Figure 2.) This sample order urgency dictates their processing, ensuring that samples from farms with higher urgency are prioritized over those arriving earlier from less critical locations. The laboratory’s daily processing capacity is limited, and once this limit is reached, we evaluated two possible scenarios: 1) any remaining samples are deferred to the next day or 2) redistributed to other certified laboratories from NAHLN (USDA, 2024). In the first scenario, the backlog of unprocessed samples accumulates, adding to the new samples received each day up to a pre-established storage capacity limit. In the second scenario, we assume that the surplus samples are redistributed on the same day they are received and that the destination laboratories can accommodate and process all redirected samples. Moreover, to enhance efficiency, our algorithm permits the laboratory to process samples individually or in pools, with a predetermined number of samples from the same farm combined into a single test (Figure 1). The maximum size of these sample pools was set at 10, based on discussions with animal health officials involved in ASFV surveillance in the U.S. (personal communication.)

#### Outputs

In this study, we estimated the sample collectors and laboratory processing capacities required for a simulated ASFV epidemic by employing a factorial combination of various conditions detailed in Table 1. Our model outputs are derived from the number of ASFV simulated outbreaks, which provided the number of at-risk farms and barns that needed to be sampled. Therefore, our results include 1) the total and daily number of sample collectors, 2) the number of days sampling was delayed, 3) the number of samples waiting to undergo processing at the laboratory, and 4) the number of samples that needed to be redistributed to other laboratories to prevent processing delays.

## Results

### African swine fever simulated epidemics

Our model simulated the spread of ASFV for 150 days. The summary of model simulations shows a median of 0 (range = 0-28) secondary infection detected daily, culminating in 27 outbreaks (range = 1-68) throughout the simulation period. In 26% of simulations, infection continued until the last day of the simulation, indicating that the ASFV epidemic was not completely eliminated. The median sampling duration in simulations, defined as the time between the first sampling event and the last sampling event, was 54 (range = 1-135) days. This duration represents the sampling under optimal conditions, where farms were sampled without any delays, and it is used as a baseline to compare our results. The daily number of farms sampled peaked at 99 (range = 1-291) on day 39 (range = 37-78), while over the 150-day timeframe, a median of 23 (range = 1-516) farms were sampled per day (Figure 4A). By day 54 (median sampling duration) of the sampling, the cumulative number of sampling events at the farm level was 420 (range = 1-635) which increased to 616 (range = 1-15,011) by day 150. At the barn level (Figure 4B), a median of 107 (range = 1-2,312) barns were sampled per day, resulting in 2,173 (range = 7-2,975) cumulative sampling events at day 54, and 3,068 (range = 7-69,118) at day 150.

**Figure 4.**
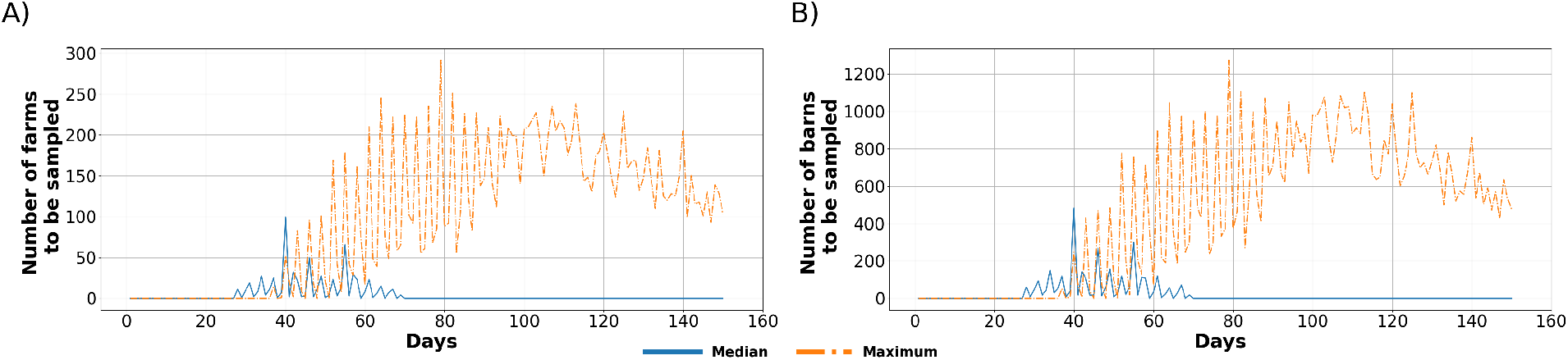
ASFV outbreak simulations. A) Number of farms sampled per day; and B) number of barns sampled per day. The blue solid line represents the median value and the orange dashed line indicates the maximum values.

### Estimated number of sample collectors

The number of sample collectors and days required for sampling throughout the simulated ASFV epidemics were significantly impacted by the number of sampler collectors available, downtime, and the sample type (blood swabs vs oral fluids) (Tables 2 and 3.) Scenarios collecting blood samples involving 72 hours of downtime, and unlimited trained sample collectors proved to be the best, completing all sampling within a median of 54 days and 135 days in the worst epidemic scenario. However, this efficiency required a significantly higher median number of sample collectors, ranging between 247 and 367, and in the worst scenario between 2,095 and 3,115 (Table 2.) Scenarios with the same downtime but limited to sample collectors available also completed sampling within a median of 54 days; however, the worst epidemic scenarios required from 139 to 292 days. Thus, in comparison to the maximum expected sampling duration of 135 days, the worst-case scenarios required an additional 4 to 157 days to sample all at-risk farms. Increasing the regular working hours allowed per collector from eight to 12 hours decreased the median number of sample collectors needed by 33%. Similarly, reducing the number of sampler collectors sent per farm from five to three led to a 4% reduction in the median number of collectors required.

**Table 2.** Number of sample collectors–median and min-max range–needed to collect 31 blood samples from each barn of at-risk farms in a simulated ASFV epidemic.

| Conditions |  |  | 72 hours downtime |  | 24 hours downtime |  | 0 hours downtime |  |
| --- | --- | --- | --- | --- | --- | --- | --- | --- |
| SCPF | TSC | RWS | Sample collectors | Last sample day | Sample collectors | Last sample day | Sample collectors | Last sample day |
| 5 | ∞ | 12 | 247 (1 - 2,095) | 54 (1 - 135) | 181 (1 - 1,206) | 54 (1 - 135) | 136 (1 - 833) | 54 (1 - 135) |
| 3 | ∞ | 12 | 238 (1 - 1,992) | 54 (1 - 135) | 176 (1 - 1,167) | 54 (1 - 135) | 137 (1 - 841) | 54 (1 - 135) |
| 5 | ∞ | 8 | 367 (1 - 3,115) | 54 (1 - 135) | 264 (1 - 1,818) | 54 (1 - 135) | 228 (1 - 1,433) | 54 (1 - 135) |
| 5 | 2,000 | 12 | 247 (1 - 1,959) | 54 (1 - 139) | 181 (1 - 1,206) | 54 (1 - 135) | 136 (1 - 833) | 54 (1 - 135) |
| 5 | 1,750 | 12 | 247 (1 - 1,750) | 54 (1 - 146) | 181 (1 - 1,206) | 54 (1 - 135) | 136 (1 - 833) | 54 (1 - 135) |
| 5 | 1,500 | 12 | 247 (1 - 1,500) | 54 (1 - 154) | 181 (1 - 1,192) | 54 (1 - 135) | 136 (1 - 833) | 54 (1 - 135) |
| 5 | 1,250 | 12 | 247 (1 - 1,250) | 54 (1 - 163) | 181 (1 - 1,138) | 54 (1 - 135) | 136 (1 - 832) | 54 (1 - 135) |
| 5 | 1,000 | 12 | 247 (1 - 1,000) | 54 (1 - 180) | 181 (1 - 1,000) | 54 (1 - 135) | 136 (1 - 830) | 54 (1 - 135) |
| 5 | 500 | 12 | 245 (1 - 500) | 54 (1 - 292) | 181 (1 - 500) | 54 (1 - 158) | 136 (1 - 500) | 54 (1 - 136) |

**Table 3.** Number of sample collectors–median and min-max range–needed to collect five oral fluid samples from each barn in at-risk farms in a simulated ASFV epidemic.

| Conditions |  |  | 72 hours downtime |  | 24 hours downtime |  | 0 hours downtime |  |
| --- | --- | --- | --- | --- | --- | --- | --- | --- |
| SCPF | TSC | RWS | Sample collectors | Last sample day | Sample collectors | Last sample day | Sample collectors | Last sample day |
| 5 | ∞ | 12 | 116 (1 - 882) | 54 (1 - 135) | 99 (1 - 573) | 54 (1 - 135) | 28 (1 - 172) | 54 (1 - 135) |
| 3 | ∞ | 12 | 116 (1 - 882) | 54 (1 - 135) | 99 (1 - 573) | 54 (1 - 135) | 35 (1 - 211) | 54 (1 - 135) |
| 5 | ∞ | 8 | 127 (1 - 1,047) | 54 (1 - 135) | 103 (1 - 619) | 54 (1 - 135) | 52 (1 - 339) | 54 (1 - 135) |
| 5 | 2,000 | 12 | 116 (1 - 882) | 54 (1 - 135) | 99 (1 - 573) | 54 (1 - 135) | 28 (1 - 172) | 54 (1 - 135) |
| 5 | 1,750 | 12 | 116 (1 - 882) | 54 (1 - 135) | 99 (1 - 573) | 54 (1 - 135) | 28 (1 - 172) | 54 (1 - 135) |
| 5 | 1,500 | 12 | 116 (1 - 882) | 54 (1 - 135) | 99 (1 - 573) | 54 (1 - 135) | 28 (1 - 172) | 54 (1 - 135) |
| 5 | 1,250 | 12 | 116 (1 - 882) | 54 (1 - 135) | 99 (1 - 573) | 54 (1 - 135) | 28 (1 - 172) | 54 (1 - 135) |
| 5 | 1,000 | 12 | 116 (1 - 866) | 54 (1 - 135) | 99 (1 - 573) | 54 (1 - 135) | 28 (1 - 172) | 54 (1 - 135) |
| 5 | 500 | 12 | 116 (1 - 500) | 54 (1 - 155) | 99 (1 - 496) | 54 (1 - 135) | 28 (1 - 172) | 54 (1 - 135) |

In scenarios with 24 hours or no downtime and at least 1,000 trained sample collectors, sampling was completed within a median of 54 days and a maximum of 135 days in the worst epidemic scenarios (Table 2). However, with only 500 sample collectors, the sampling time exceeded 135 days in the worst epidemic scenarios, requiring an additional 23 days for the 24-hour downtime scenario and one extra day for the no-downtime scenario. For scenarios with 24 hours of downtime, the median number of required sample collectors ranged from 181 to 264, with the maximum ranging from 500 to 1,818. Scenarios with no downtime required a median of 136 to 228 sample collectors and a maximum of 500 to 1,433 in the worst epidemic scenarios (Table 2).

In our alternative scenarios using oral fluids, sampling was completed within a median of 54 days and a maximum of 135 days in all tested scenarios (Table 3.) Additionally, the number of sample collectors needed for oral fluids compared with blood samples was notably lower (Tables 2 and 3.) Specifically, in scenarios with 72 hours of downtime, oral fluid sampling required a median between 51% and 65% fewer sample collectors, while in the worst epidemic scenario required between 0% and 66% fewer sample collectors (Table 3.) In scenarios with 24 hours of downtime, the reduction in the median number of sample collectors ranged between 43% and 61%, and between 1% and 66% in the worst epidemic scenario. Finally, in scenarios without any downtime, the median reduction was between 74% and 79%, and 66% and 79% in the worst epidemic scenario (Table 3.)

The median and maximum demand for sample collectors gradually increased over the first week after the sampling started (Figure 5). The median daily growth rate of sample collectors for blood samples ranged between 5.1% and 34.4%, but in the worst epidemic scenario, it varied between 2.5% and 5.9%. For oral fluid samples, the median daily growth rate was between 8.1% and 182%, while in the worst epidemic scenario, it fluctuated between 2.5% and 6.0%.

**Figure 5.**
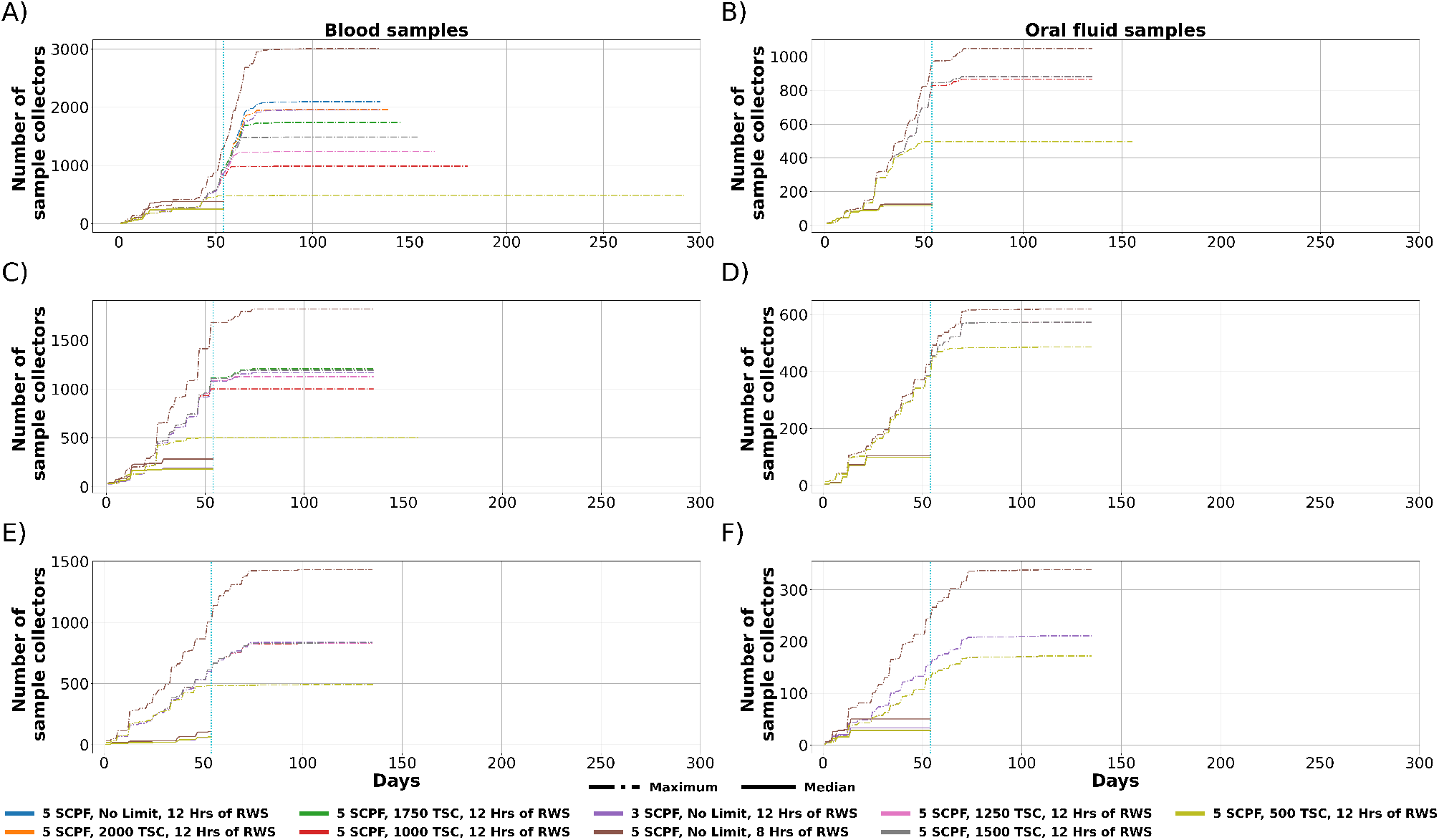
Cumulative number of sample collectors needed during the simulated ASFV epidemic considering 31 blood or five oral fluid samples from each barn in at-risk farms. The x-axis terminates at the median sampling duration (54 days) where the total number of samplers plateaus. A) blood samples with 72 hours downtime, B) oral fluid samples with 72 hours downtime, C) blood samples with 24 hours downtime, D) oral fluid samples with 24 hours downtime, E) blood samples with 0 hours downtime, F) oral fluid samples with 0 hours downtime. SCPF = Sample collectors per farm, TSC = Trained sample collectors, RWS = Regular working schedule.

An optimal ASFV response requires that at-risk farms be sampled on the same day they are identified to avoid delays. Among the scenarios evaluated, collecting blood samples with a 72-hour downtime resulted in significant sampling delays (Figure 6.) Specifically, the scenario exhibited an average delay of 23 days and a maximum delay of 247 days with 500 trained sample collectors, marking it as the least effective. Conversely, scenarios with an unlimited number of trained sample collectors, a downtime of 24 or zero hours, or the collection of oral fluid samples demonstrated considerably lower sampling delays, with a median delay that ranged between zero and six days (Figure 6.)

**Figure 6.**
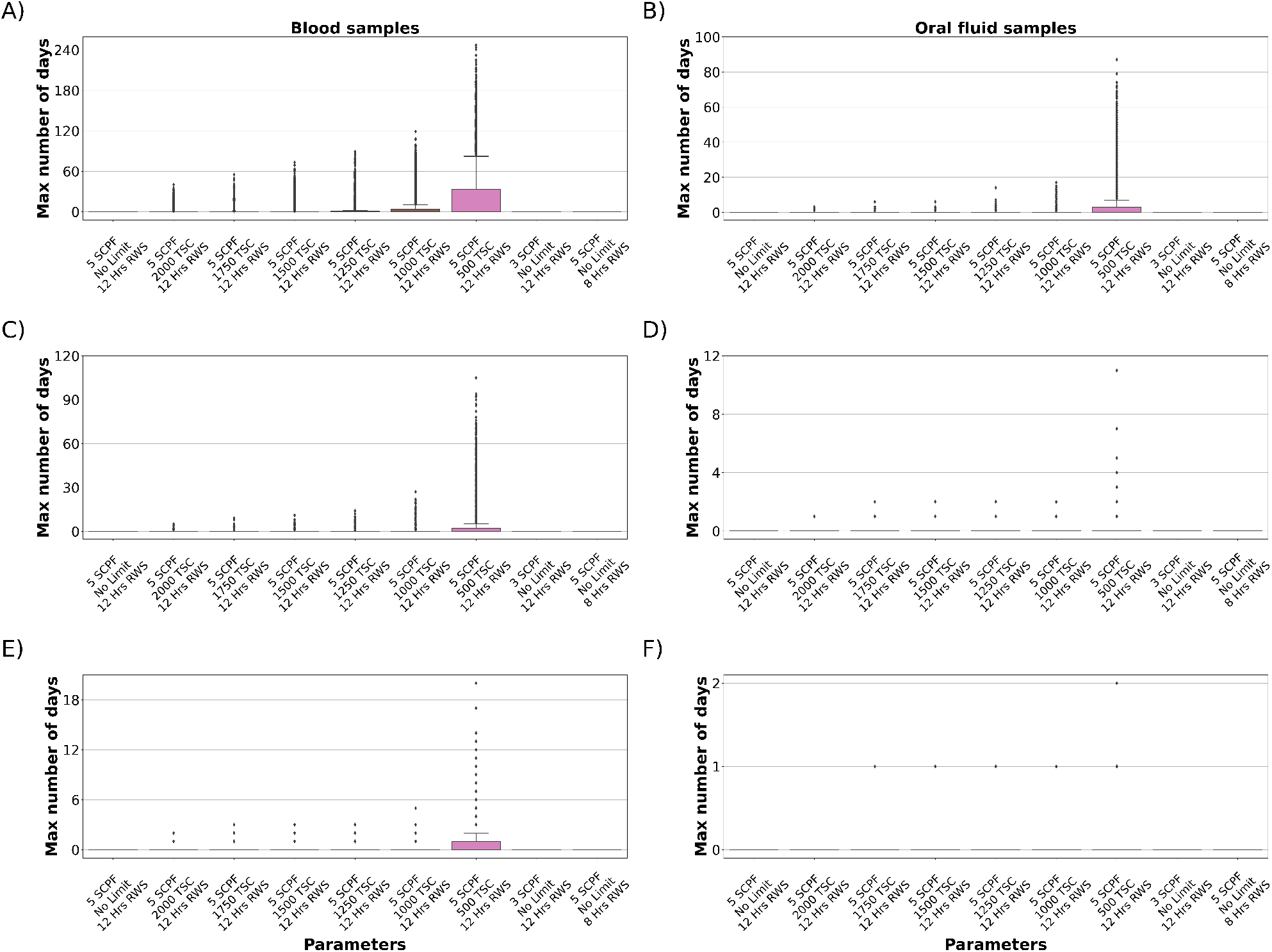
Maximum number of days sampling is delayed during epidemic; SCPF = Sample collectors per farm, TSC = Trained sample collectors, RWS = Regular working schedule. A) blood samples with 72 hours downtime, B) oral fluid samples with 72 hours downtime, C) blood samples with 24 hours downtime, D) oral fluid samples with 24 hours downtime, E) blood samples with 0 hours downtime, F) oral fluid samples with 0 hours downtime.

### Estimating laboratory capacity requirements

Among the different scenarios evaluated, the median number of blood samples received at the laboratory was 84,830 (range = 52 -2,066,831) (Table 4), and a median of 1,530 (range = 0-68,939) per day (Supplementary Material Figure S1.) Regardless of downtime, unprocessed individual blood samples began accumulating rapidly after the sampling was initiated, reaching a peak of 40,618 (range = 52 -1,955,126) unprocessed samples (Supplementary Material Figure S2-S37.) Over time, the unprocessed samples gradually reduced, yet processing all samples required a median of 92 days and, in the worst-case scenario, more than 5.7 years. Implementing sample pooling by five and 10 significantly reduced the processing time. For pools of five samples, the processing period was shortened to a median of 56 days and 1.2 years in the worst scenario (Table 4), a reduction of 39% and 79% in time compared to individual sample processing, respectively. For pools of 10 samples, the median time required was 55 days, reducing the processing time by 40% compared to scenarios without pooling. In the worst scenario, the median time required for pools of 10 samples ranged from 234 days to 293 days (Table 4), reducing the processing time between 85% and 88% compared to scenarios without pooling.

**Table 4.**
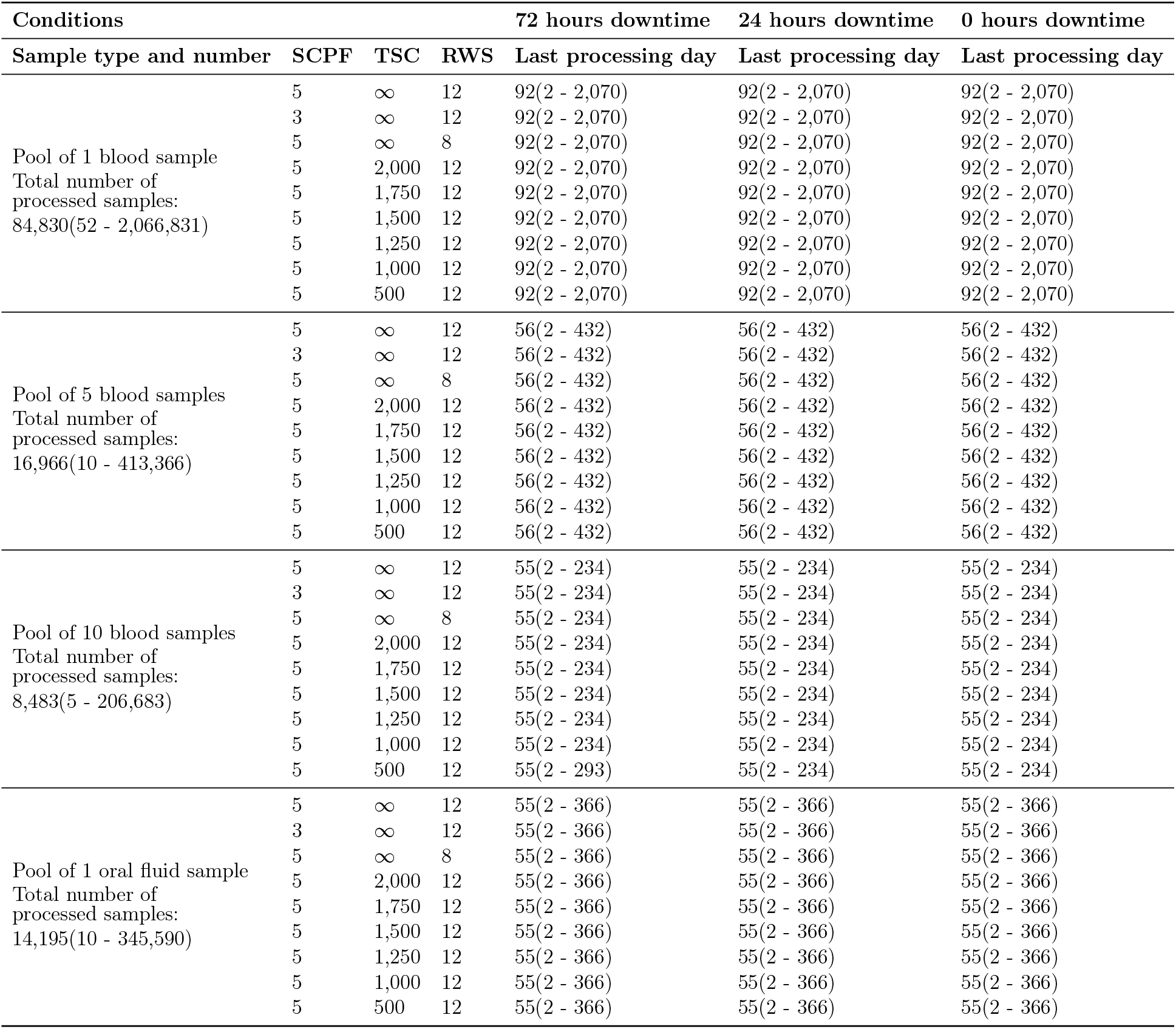
Number of samples–median and min-max range–arrived at the laboratory and needed to be processed in a simulated ASFV epidemic.

The alternative scenario of collecting oral fluid samples showed that the laboratory received a median of 14,195 (range = 10 -345,590) samples (Table 4), with a median of 255 samples per day (Supplementary Material Figure S1.) Independent of the conditions evaluated, the time required to process these samples was 55 days and one year in the worst scenario, reducing the processing time between 40% and 82% compared to individual blood samples, respectively.

The median number of unprocessed single blood samples that the laboratory needed to redistribute was 10,062, given a daily capacity of 1,000 samples 7. This number decreases to 8,562 and 6,062 for increased capacities of 2,500 and 5,000 samples per day, respectively.

On the days requiring the most redistribution—top 25%—the range of samples varied from 19,434 to 67,940 with a capacity of 1,000 samples per day, from 17,934 to 66,440 with a capacity of 2,500 samples per day, and from 15,434 to 63,490 with a capacity of 5,000 samples per day. Pooling samples remarkably reduces the need for redistribution. With the three different laboratory capacities considered, pooling five samples decreased the median daily redistributed samples by 88% to 100% and the maximum by 83% to 91%. Similarly, a pool of 10 samples reduced the median daily redistributed samples by 99% to 100% and the maximum by 91% to 97%. With a laboratory capacity of 1,000 samples per day, the median number of total samples that needed redistribution ranged from 118 to 57,826, while the maximum ranged from 115,783 to 1,958,888 (Supplementary Material Figure S38). Increasing the laboratory capacity to 2,500 decreased the median values by 43% and 100%,and further increasing the capacity to 5,000 reduced the median values by 75% and 100%.

## Discussion

Our results reveal that the number of sample collectors required in an ASFV outbreak was mostly impacted by the epidemic size, downtime requirements, and sample type blood versus oral fluids. For the study area, having 238 (range = 1-1,992) sample collectors was estimated to be sufficient to complete sampling without delays. Particularly, scenarios with no downtime or 24-hour downtime between sampling exhibited the lowest testing delays, highlighting the benefit of continuous operational capacity in surveillance systems but also introducing the risk of further dissemination as multiple farms were visited by the same collector. Additionally, oral fluid sampling required fewer collectors than blood samples. Oral fluid samples not only required fewer resources in terms of sample collectors but also contributed to shorter laboratory processing time (Goonewardene et al., 2021b; Mur et al., 2013). This suggests that integrating less conventional sampling techniques could enhance surveillance efficiency, especially during large-scale outbreaks where traditional resources are stretched thin. While it is unrealistic to expect that oral fluids could be used alone to address testing needs, they could become important tools in some use cases or could be used as complementary surveillance tools to conventional sampling techniques. The significant backlog of unprocessed samples in the laboratory, even with increased pooling and processing capacity, further illustrates the need for improvements in laboratory processing capabilities and or complement with additional strategies, such as oral fluid samples or sample redistribution to other NAHLN laboratories (Goonewardene et al., 2021b; USDA, 2024). Therefore, these findings advocate for an integrated approach to disease surveillance that enhances both field sample collection and laboratory processing capabilities, ensuring rapid response to outbreaks and effective disease management for future ASFV outbreaks or similar animal health emergencies in the U.S.

A large number of farms were identified as at-risk during the simulated ASFV epidemic, primarily due to the high density of farms in the studied regions—a median of 48 (range = 0-128) farms within a 10 km radius of one another (Supplementary Material Figure S39). As a result, many farms were within the control zones (infected, buffer, and surveillance areas) (USDA, 2023). Furthermore, the frequent movement of vehicles, especially those transporting feed and previously in contact with infected farms, further increased the number of at-risk farms (Galvis and Machado, 2024a). As a result, a large number of samples need to be collected, which impacts the demand for sample collection, necessitating a larger workforce of trained sample collectors and extending laboratory processing times. Alternatives to reduce the number of samples could include, for example, the use of oral fluids, which would reduce the number of farm visits, which is a concern for breeding farms with high biosecurity levels (Sykes et al., 2022; Harlow et al., 2024; Campler et al., 2024; Alarcón et al., 2021). Reducing the number of vehicle visits would reduce the number of samples; thus, implementing stringent policies on vehicle movements should ensure vehicles are decontaminated between farm visits and thus represent a low risk of disease transmission (Galvis and Machado, 2024a). However, this strategy hinges on the effectiveness of cleaning and disinfection protocols, which are not always reliable (Boniotti et al., 2018; Mannion et al., 2008). Another proactive measure could involve rerouting vehicle movements during the epidemic based on the ASFV status of farms, to minimize contact with infected sites (Galvis and Machado, 2024b). These measures could significantly reduce the number of at-risk farms and lessen the burden of sample collection.

The ASFV response scenario, which involves collecting blood samples from 31 animals per barn as the preferred sampling schema and implementing a 72-hour downtime between farm visits presents numerous logistical challenges. We demonstrated that in the worst-case scenarios, these requirements could delay farm sampling by up to 247 days (Figure 6). As a reflection, our findings suggested the need for the study region of 1,992 trained sample collectors (Table 2), which is indeed a large number and is expected to be costly for both the swine industry and the animal health officials who are in charge of certifying sampler collectors (Secure Pork Supply Plan, 2024). To reduce the number of sample collectors while maintaining efficient sampling times, our model shows that shorter downtime would have a notable benefit in completing sampling without delays. For instance, a downtime of 24 hours would necessitate between 8% and 49% fewer collectors, and eliminating downtime would require between 28% and 75% fewer collectors (Table 2). However, reducing or eliminating downtime requirements may increase the probability of between-farm disease transmission due to human-mediated transmission (Bellini et al., 2021). Another alternative solution could be to switch to oral fluid sampling, which requires fewer samples per farm and can be conducted more rapidly than blood sampling. For oral fluid sampling, the number of necessary collectors was reduced from 47% to 75% with a 72-hour downtime, from 69% to 75% with a 24-hour downtime, and from 83% to 91% with no downtime (Table 3). In addition to downtime and sample type, working schedules including a maximum of three sample collectors per farm with extended working hours—from eight to 12 hours—proved more effective, reducing collectors between 3% and 33% (Table 2). These alternatives not only streamline operations but could also ensure a faster, more cost-effective approach to managing ASFV outbreaks, ultimately enhancing our ability to control the spread of the virus promptly.

Experimental studies have demonstrated the effectiveness of collecting oral fluid samples via ropes in pens for identifying ASFV cases within 3-5 days of virus introduction (Goonewardene et al., 2021b). However, this method has noted limitations, such as variable interaction of pigs with the ropes limiting the ability to demonstrate if all pigs in a pen contributed to the sample (Grau et al., 2015; Guinat et al., 2014; Goonewardene et al., 2021b), environmental contamination that can skew results, false negatives before or within the first 3–5 days after exposure and limited field validation studies that affect the reliability of this approach (Goonewardene et al., 2021b). Therefore, while oral fluid sampling is a promising sampling method, it should be used cautiously and ideally complemented by other sampling strategies until further clinical trials provide support for its use alone.

Our findings reveal that the laboratory’s sample processing capacity cap leads to potential processing delays of up to 5.7 years for blood samples and up to one year for oral fluid samples without pooling. These results highlight the necessity of sample pooling to prevent delays, which can be conducted either in the laboratory or field as outlined in the certified swine sample collector program (Secure Pork Supply Plan, 2024). For instance, in our scenario with a pool of 10 samples, we observed a median processing delay of zero days. However, this strategy did not consistently ensure timely sample processing in the worst epidemic scenarios, leading to delays in ASFV detection and control measures. Thus, a pooling size of 10 samples may not be universally sufficient for all ASFV epidemic scenarios. Our results also suggest that the size of the sample pool should be adjusted based on the scale and timing of the epidemic and the number of at-risk farms throughout its course, starting with pools of one to five samples and increasing as necessary. However, it should be considered that larger pools are more likely to dilute the virus genetic material dramatically, impacting the performance of the test and thus the ability to accurately detect disease (Aira et al., 2019; Pikalo et al., 2021; López et al., 2021; Vilalta et al., 2019; Gallardo et al., 2019). An alternative approach could involve enhancing the laboratory’s daily processing capacity and redistributing unprocessed samples to other NAHLN laboratories (USDA, 2024). We demonstrated that in scenarios without pooling, the laboratory could reach its processing capacity from day one (Supplementary Material Figures S40 and S41), requiring a daily distribution of 10,062 (range = 2-67,940) samples (Figure 7). Such a large volume may potentially overwhelm the processing capacities of these other laboratories, which may also be receiving and processing samples from other states during an ASFV epidemic in the U.S. Additionally, assuming all other NAHLN laboratories have a similar capacity of 1,000 samples per day, 50% of cases would require collaboration with up to 10 NAHLN laboratories (Supplementary Material Figures S40), while in the worst epidemic scenario would require up to 50 laboratories (Supplementary Material Figures S41). Ideally, raising the daily laboratory capacity to 5,000 samples reduces the number of daily samples needing redistribution to 0 (range = 0-8,788) for pools of five samples and 0 (range = 0-1,894) for pools of 10 samples (7). Similarly, pooling samples also reduces the number of NAHLN laboratories needed during an ASFV epidemic (Supplementary Material Figures S42 -S45). Finally, portable devices capable of detecting ASFV directly on farms, despite potentially increasing processing costs, could significantly reduce sample processing delays (Chen et al., 2021; Ngan et al., 2023). Therefore, subsequent studies should include cost-benefit analyses to evaluate the effectiveness of these devices as part of an ASFV detection strategy while accounting for potential proficiency testing challenges with on-farm sample collectors and variations in diagnostic test performance.

**Figure 7.**
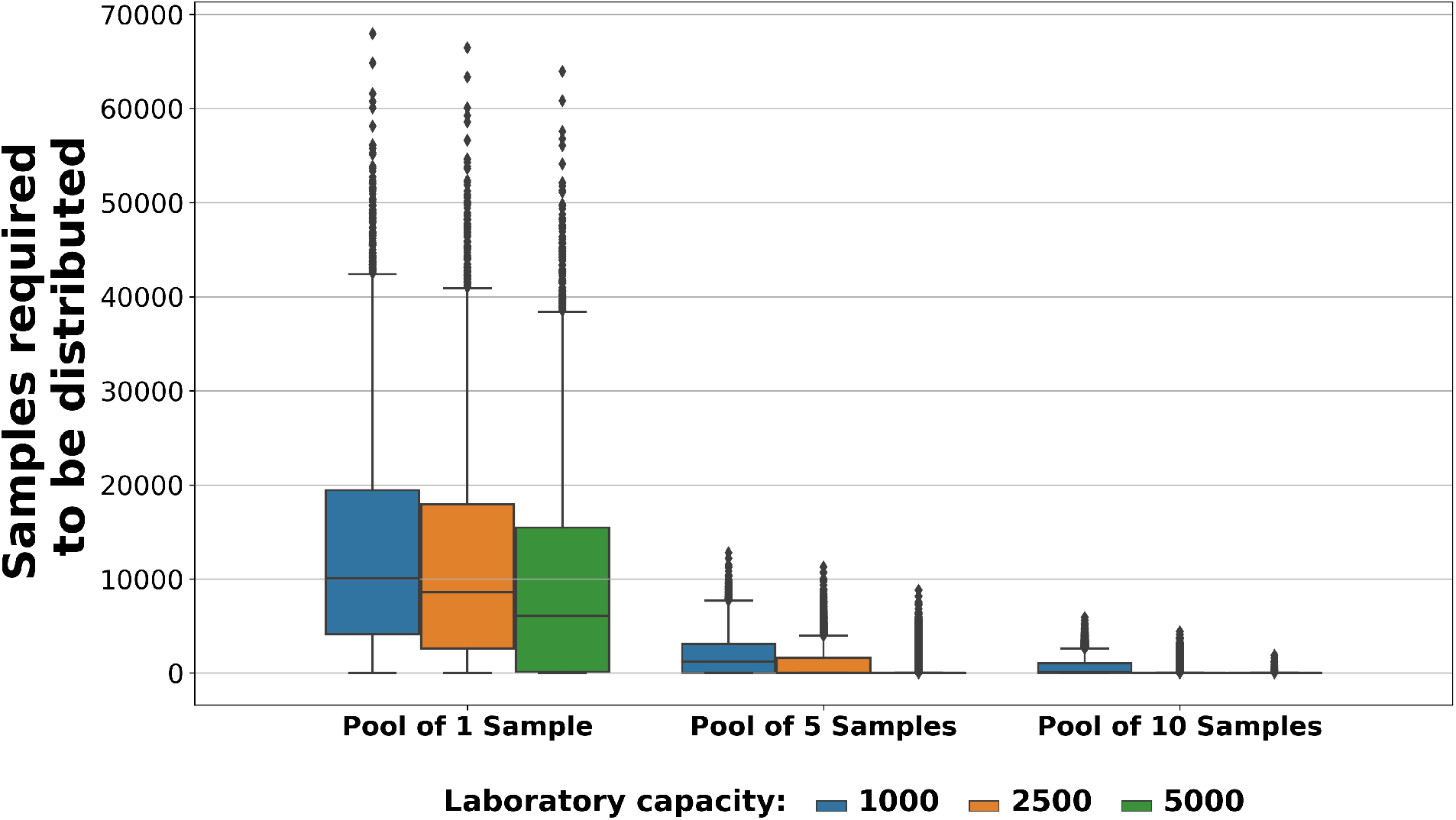
Daily number of samples required to be distributed to other laboratories in a sampling scenario using 72 hours downtime with 3 SCPF, 12 RWS, and no limit on TSC. SCPF = Sample collectors per farm, TSC = Trained sample collectors, RWS = Regular working schedule.

### Limitations and further remarks

This study has several limitations that require careful consideration when interpreting results. The ASFV model employed was calibrated using data from a previous epidemic of porcine epidemic diarrhea virus in the U.S., aimed at robustly predicting the spread of a new disease in a susceptible population. Despite this, the model may not accurately reflect the actual number of ASFV outbreaks; it could either overestimate or underestimate the number of at-risk farms and, consequently, the number of samples and sample collectors required (Sykes et al., 2023). We developed an algorithm to emulate real-world conditions and closely estimate the needs for sample collectors and laboratory capacity. However, we recognize that some conditions assumed in our methodology may differ during an actual ASFV outbreak, potentially altering the number of sample collectors needed and sample processing time. Additionally, we excluded several key variables that could impact the results. These include the use of auxiliary drop-off locations to reduce travel times for sample collectors, modeling samples redistributed to certified laboratories from the NAHLN including their processing capacities and locations, and a validation period during which farms, once tested, are not required to be retested for a specified number of days. Furthermore, we omitted sampling strategies that involve on-site trained farm workers, such as caretakers, despite their training being a key objective of the certified swine sample collector program to meet the high demand for manpower during an epidemic (Secure Pork Supply Plan, 2024). By enabling farm workers to collect samples on-site, the need for additional travel and external sample collectors is reduced, thereby streamlining the sampling process and saving time. Incorporating these scenarios into future studies will yield more accurate results regarding the sampling process.

Another significant limitation is that the ASFV model was not run simultaneously with the sampling estimation algorithm. This absence of integration directly affects the transmission dynamics of ASFV, as delays in sampling can decrease the number of infected farms detected by active surveillance during an outbreak. Consequently, the effectiveness of control measures is compromised, allowing infected farms to spread the virus over longer periods and thereby extending the time required to eradicate the disease. Thus, a future approach requires an integration of these two models to provide more valuable results for the ASFV surveillance, control, and eradication plans.

Although the oral fluid sampling strategy demonstrated promising results, it alone is not considered in the current ASFV response plan (USDA, 2023). Therefore, it is necessary to model more realistic scenarios that combine oral fluid and blood sampling strategies across various use cases throughout the simulated epidemic. It is important to note that our study does not account for the sampling of wild boars, which play a crucial role in the spread of ASFV (Sauter-Louis et al., 2021; Sánchez-Cordón et al., 2019). Including wild boar sampling would require estimating sample sizes and wild boar locations, as well as employing personnel specifically trained for wild boar sampling, which differs from farm sampling techniques (Engeman et al., 2013). Additional sampling strategies should also consider fomites and vehicles that have been exposed, as they can serve as long-term virus reservoirs (Gebhardt et al., 2022). Moreover, the barn-level sampling strategy presents limitations, particularly in barns with multiple rooms. The heterogeneous distribution of infected animals across different rooms could increase the likelihood of false negative results due to inadequate sampling (Lamberga et al., 2022). Therefore, it is crucial to compare barn and room-level sampling strategies to determine the most effective approach for identifying ASFV-positive farms. Our results highlight the challenges sampling strategies would face should ASFV be introduced into the country. Addressing these challenges necessitates the development of additional strategies to optimize resource utilization. Consequently, our future efforts will focus on enhancing the sample collector allocation process through the use of advanced optimization methods, including deep reinforcement learning algorithms to further extend the current sampling and diagnostic capacities, utility attempting to identify the best set of tactics capable of fulfilling the needed actions in stamping out any ASFV incursion.

Our study remarks on the importance of optimizing sample collector allocation and laboratory capacities for efficient and strategic resource management in the surveillance and control of ASFV. Similarly, we emphasize the certified swine sample collector program’s crucial role in training individuals for effective sample handling during large-scale epidemics and encourage regulatory agencies and the swine industry to combine efforts to ensure that sample collectors can continue to be trained, certified, and tracked for ready mobilization (Secure Pork Supply Plan, 2024). Efficient sample collection is pivotal not only for early detection but also for the effective containment and mitigation of outbreaks. As demonstrated by our results, improving the precision and responsiveness of sample collector deployment can significantly strengthen disease surveillance systems and expedite emergency responses. This becomes even more crucial in scenarios where rapid response and adaptability are necessary to manage dynamic and potentially widespread epidemics. Ultimately, investing in and refining these aspects of surveillance infrastructure are vital steps toward safeguarding animal health and, by extension, related agricultural and economic sectors.

## Conclusion

We developed a strategic sampling model tailored to real-world conditions to accurately estimate the required number of sample collectors and necessary laboratory capacities in response to an ASFV introduction in the U.S. Our findings indicate that the number of samplers varies significantly mainly due to the size a possible epidemic, the sampling strategies employed and sample collector downtime. Depending on the sampling conditions, 238 sample collectors are enough in 50% of the ASFV epidemic scenarios, but in the worst-case scenarios, 1,992 sample collectors are needed to complete sampling the same day the sample is required. However, it can be reduced between 28% and 75% with more flexible conditions that eliminate downtime or between 47% and 75% when compared to oral fluid sample collection. Our study also highlighted significant challenges in processing samples when laboratory capacity was capped at 1,000 samples per day, with processing times extending over several years when not pooling. This underscores the urgent need for sample pooling, increasing laboratory capacities, or redistributing unprocessed samples to USDA-certified laboratories from the NAHLN. Therefore, it is crucial to revisit and refine ASFV sampling strategies to ensure efficient resource allocation and enhance disease preparedness and management capabilities for potential ASFV outbreaks.

## Acknowledgments

The authors would like to acknowledge the valuable advice provided by participating companies and members of the Animal and Plant Health Inspection Service (APHIS): Barb Porter Spalding, Columb Rigney, Holden Hutchinson, Lindsey Holstrom, Lisa Rochette, Lydia Carpenter, and Sasidhar Malladi.

## Authors’ contributions

JAG, DR, and GM conceived the study. JAG, MYS, ALS, KCO, LR, DR, and GM participated in the study design. JAG and ALS prepared population and movement data and developed the initial modeling computer code adapted here for the ASF. ALS designed the ASFV model and simulated scenarios. MYS designed a sample collector model and computational analysis. JAG, MYS, and GM wrote and edited the manuscript. All authors discussed the results and critically reviewed the manuscript. GM secured the funding.

## Data Availability Statement

The data supporting this study’s findings are not publicly available and are protected by confidential agreements; therefore, they are not available.

