## Supplementary material for "Estimating sampling and laboratory capacity for a simulated African swine fever outbreak in the United States": ss

### Appendices

#### A. Pseudocode for sample collector simulator

*These authors contributed equally to this work.
**Corresponding Author.

**Vars:** Sampler Limit: TNS; Samplers Per Farm: SPF, Time to Work: TTW
$urgencyLevel1 \leftarrow ['CONTACT','INFECTED','SURVEILLANCE','BUFFER']$
$urgencyLevel2 \leftarrow [['SOW','GILT','BOARSTUD','FARROWTOFINISH'], ['NURSERY','WEANTOFINISH'],'FINISHER',['ISOLATION','OTHER']]$
farms, offices, labs ← Load the farm, office and lab data
distances, times ← Calculate the travel distances and times using OSRM
$samplers \leftarrow \{\}$
**if** TNS ≠ 'No Limit' **then**
  **for** $o_i \in$ offices **do**
    $samplerLimits[o_i] \leftarrow \frac{\sum_{o_i} farms[o_i]}{\sum_{o_j \in offices} farms[o_j]} * TNS$
  **end for**
**end if**
$samplers \leftarrow generateNewSamplers(office, samplerLimits)$
**while** there are unprocessed farms **do**
  $currentFarms \leftarrow$ farms.getFarmsForCurrentDay()
  $urgentFarms \leftarrow$ currentFarms.getMostUrgent(urgencyLevel1, urgencyLevel2)
  **while** $urgentFarms \neq \emptyset$ **and** there are available samplers **do**
    $farmID, samplerID \leftarrow argmin(times[urgentFarms])$
    $numSampler \leftarrow 1$
    $timeToSample \leftarrow animalNum * samplingTime/numSampler$
    $sampler \leftarrow samplers[samplerID]$
    $timeLeft \leftarrow TTW - sampler.getTime() - times[farm, sampler] - times[lab, farm] - timeToSample$
    **while** $timeLeft \leq 0$ **and** $numSampler < SPF$ **do**
      Generate a new sampler if no available samplers
      Assign one more sampler to the farm
      $numSampler \leftarrow numSampler + 1$
      Recalculate timeLeft and timeToSample
    **end while**
    Send sampler to the farm
    Increment samples sampler has by amount of samples taken
    **if** $TTW <$ sampler.getTime() + times[lab, sampler] **then**
      Send sampler to lab
      Increment the samples in the lab by the amount of samples taken
    **end if**
    **if** there are no available samplers **then**
      $samplers \leftarrow generateNewSamplers(office, samplerLimits)$
    **end if**
  **end while**
  **if** the end of the day **then**
    Process samples in lab equal to lab capacity
    Reset sampler location to office
    Reset sampler times
    Move farms left in urgentFarms to next day if there are unprocessed farms
  **end if**
**end while**

**Algorithm 1:** Sampler simulation algorithm

#### B. Additional information

Table 1: Recommended sampling scheme for an ASFV outbreak response according to USDA guidelines from 2020

| Sampling reason | Frequency of sampling | Sampling duration |
|---|---|---|
| Infected zone | Every 3 days for 2 samplings, then every 6 days | Duration of quarantine |
| Buffer zone | Every 6 days | Duration of quarantine |
| Surveillance zone | Within 15 days of first ASF detection, then every 15 days or as new zones are designated | Duration of quarantine |
| Contact tracing for animal and vehicle movements | Every 6 days | Duration of quarantine |
| Pre-movement permit | 1 and 3 days before movement | Duration of quarantine |

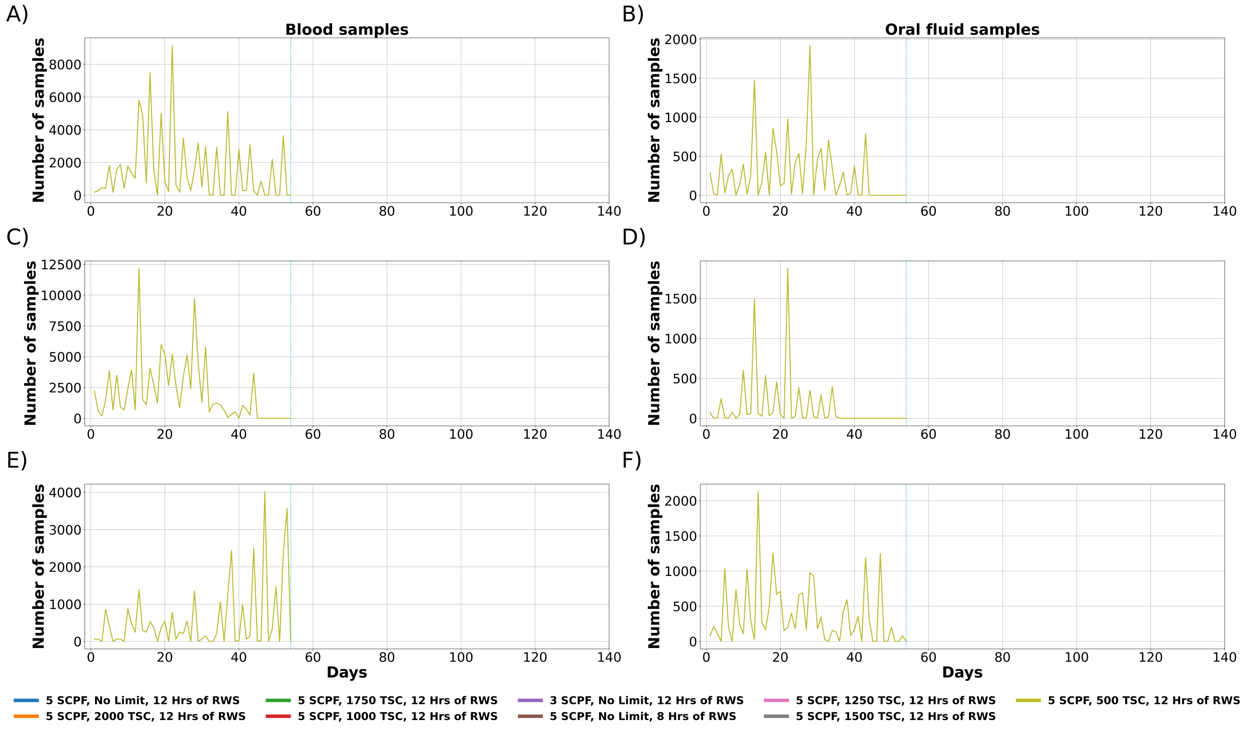


Figure 1: Median daily number of samples sent to the laboratory. All SCPF, TSC and RWS conditions result in very similar median values, causing the line graphs to significantly overlap. SCPF = Sampler collectors per farm, TSC = Trained sample collectors, RWS = Regular working schedule. A) 72 Hours Downtime, B) Pooling with 72 Hours Downtime, C) 24 Hours Downtime, D) Pooling with 24 Hours Downtime, E) 0 Hours Downtime, F) Pooling with 0 Hours Downtime.

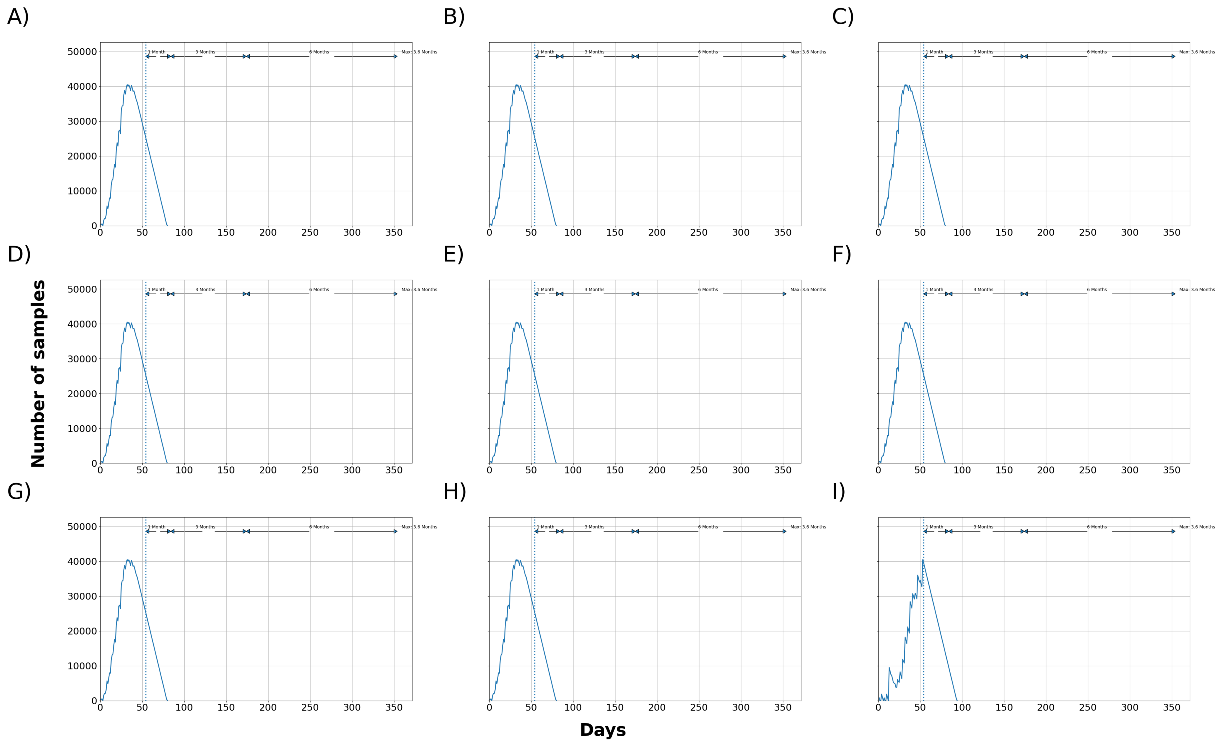


Figure 2: Median total number of samples waiting to be processed at the laboratory. 72 hours downtime and pool of 1 blood sample. SCPF = Sampler collectors per farm, TSC = Trained sample collectors. A) 5 SCPF, No Limit, 12 Hrs of RWS, B) 5 SCPF, 2000 TSC, 12 Hrs of RWS, C) 5 SCPF, 1750 TSC, 12 Hrs of RWS, D) 5 SCPF, 1000 TSC, 12 Hrs of RWS, E) 3 SCPF, No Limit, 12 Hrs of RWS, F) 5 SCPF, No Limit, 8 Hrs of RWS, G) 5 SCPF, 1250 TSC, 12 Hrs of RWS, H) 5 SCPF, 1500 TSC, 12 Hrs of RWS, I) 5 SCPF, 500 TSC, 12 Hrs of RWS.

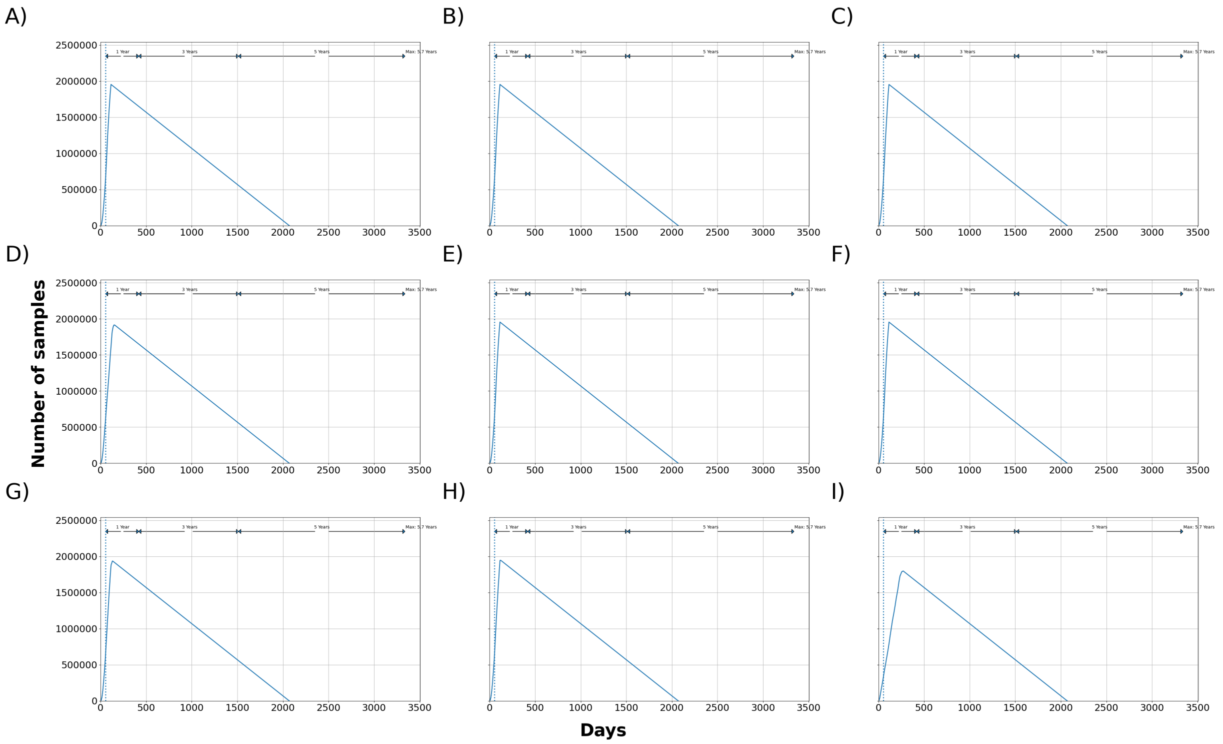


Figure 3: Maximum total number of samples waiting to be processed at the laboratory. 72 hours downtime and pool of 1 blood sample. SCPF = Sampler collectors per farm, TSC = Trained sample collectors. A) 5 SCPF, No Limit, 12 Hrs of RWS, B) 5 SCPF, 2000 TSC, 12 Hrs of RWS, C) 5 SCPF, 1750 TSC, 12 Hrs of RWS, D) 5 SCPF, 1000 TSC, 12 Hrs of RWS, E) 3 SCPF, No Limit, 12 Hrs of RWS, F) 5 SCPF, No Limit, 8 Hrs of RWS, G) 5 SCPF, 1250 TSC, 12 Hrs of RWS, H) 5 SCPF, 1500 TSC, 12 Hrs of RWS, I) 5 SCPF, 500 TSC, 12 Hrs of RWS.

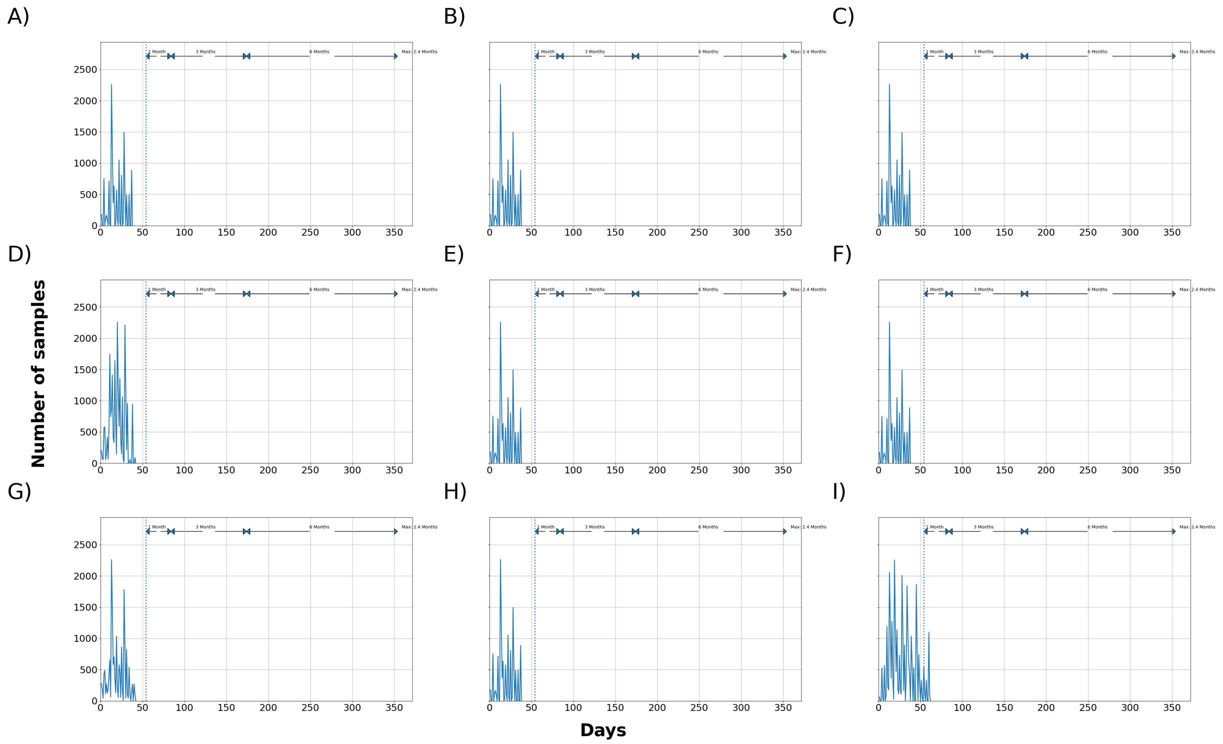


Figure 4: Median total number of samples waiting to be processed at the laboratory. 72 hours downtime and pool of 5 blood samples. SCPF = Sampler collectors per farm, TSC = Trained sample collectors. A) 5 SCPF, No Limit, 12 Hrs of RWS, B) 5 SCPF, 2000 TSC, 12 Hrs of RWS, C) 5 SCPF, 1750 TSC, 12 Hrs of RWS, D) 5 SCPF, 1000 TSC, 12 Hrs of RWS, E) 3 SCPF, No Limit, 12 Hrs of RWS, F) 5 SCPF, No Limit, 8 Hrs of RWS, G) 5 SCPF, 1250 TSC, 12 Hrs of RWS, H) 5 SCPF, 1500 TSC, 12 Hrs of RWS, I) 5 SCPF, 500 TSC, 12 Hrs of RWS.

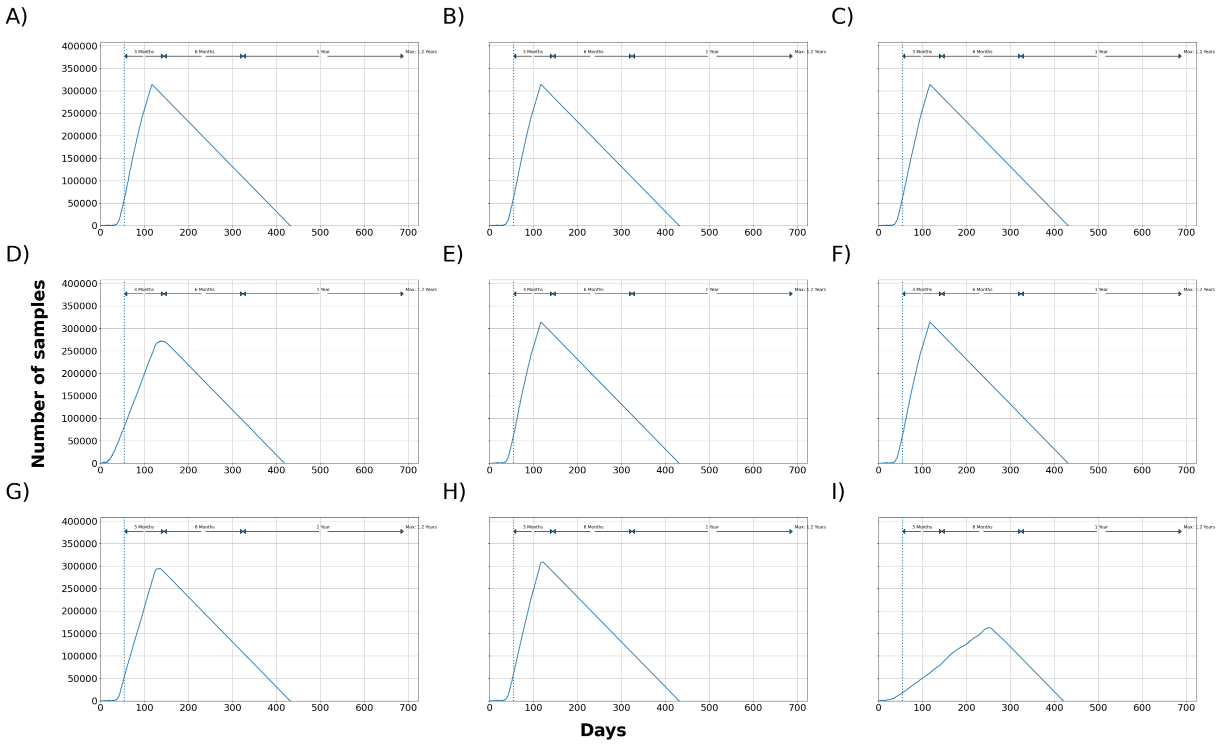


Figure 5: Maximum total number of samples waiting to be processed at the laboratory. 72 hours downtime and pool of 5 blood sample. SCPF = Sampler collectors per farm, TSC = Trained sample collectors. A) 5 SCPF, No Limit, 12 Hrs of RWS, B) 5 SCPF, 2000 TSC, 12 Hrs of RWS, C) 5 SCPF, 1750 TSC, 12 Hrs of RWS, D) 5 SCPF, 1000 TSC, 12 Hrs of RWS, E) 3 SCPF, No Limit, 12 Hrs of RWS, F) 5 SCPF, No Limit, 8 Hrs of RWS, G) 5 SCPF, 1250 TSC, 12 Hrs of RWS, H) 5 SCPF, 1500 TSC, 12 Hrs of RWS, I) 5 SCPF, 500 TSC, 12 Hrs of RWS.

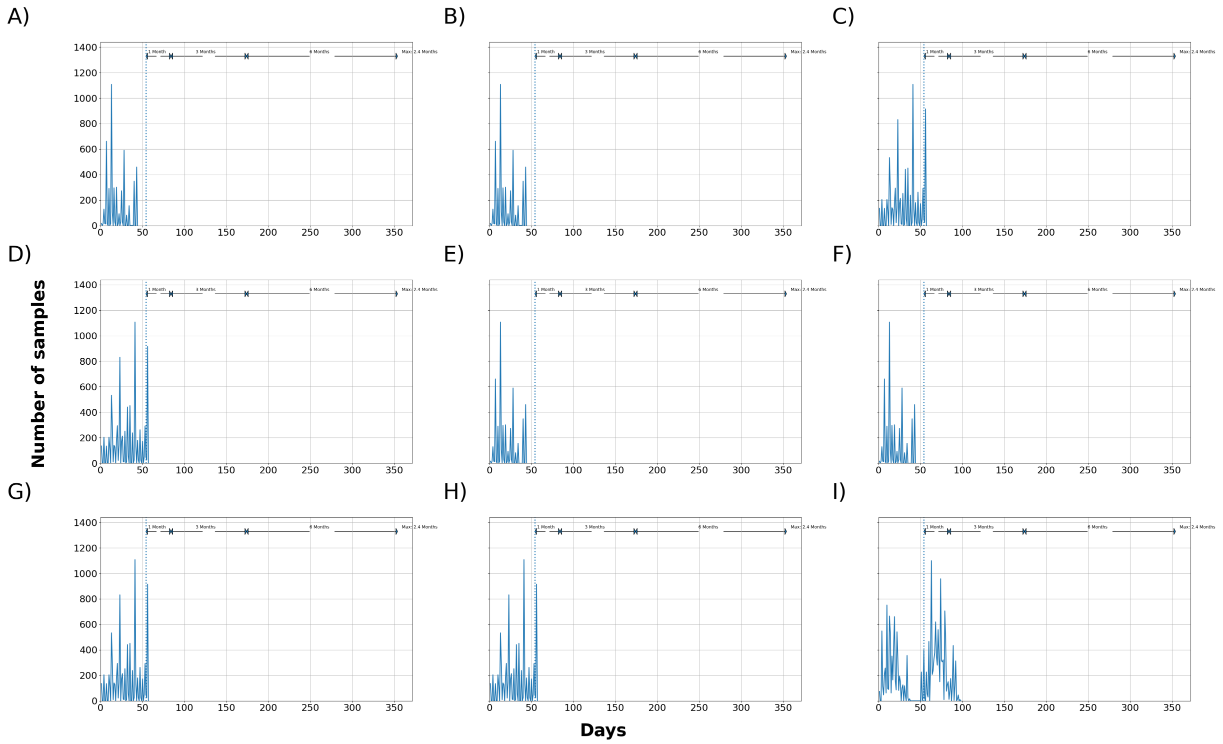


Figure 6: Median total number of samples waiting to be processed at the laboratory. 72 hours downtime and pool of 10 blood samples. SCPF = Sampler collectors per farm, TSC = Trained sample collectors. A) 5 SCPF, No Limit, 12 Hrs of RWS, B) 5 SCPF, 2000 TSC, 12 Hrs of RWS, C) 5 SCPF, 1750 TSC, 12 Hrs of RWS, D) 5 SCPF, 1000 TSC, 12 Hrs of RWS, E) 3 SCPF, No Limit, 12 Hrs of RWS, F) 5 SCPF, No Limit, 8 Hrs of RWS, G) 5 SCPF, 1250 TSC, 12 Hrs of RWS, H) 5 SCPF, 1500 TSC, 12 Hrs of RWS, I) 5 SCPF, 500 TSC, 12 Hrs of RWS.

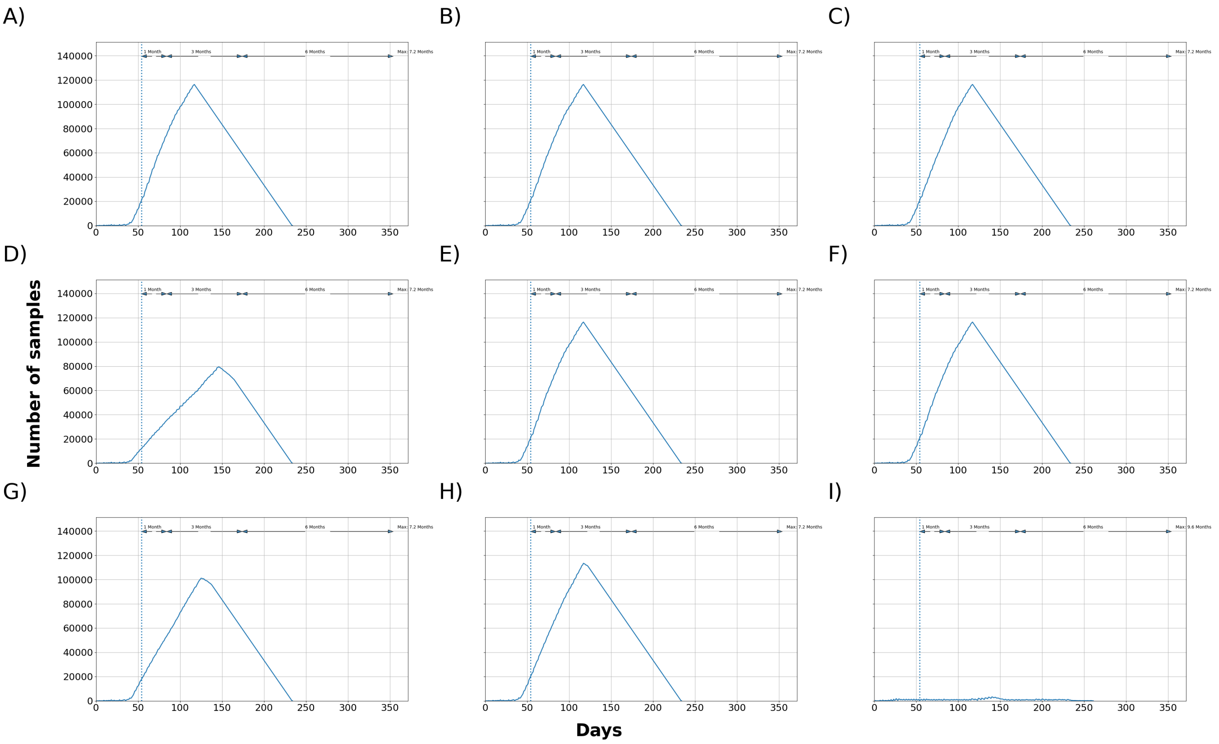


Figure 7: Maximum total number of samples waiting to be processed at the laboratory. 72 hours downtime and pool of 10 blood samples. SCPF = Sampler collectors per farm, TSC = Trained sample collectors. A) 5 SCPF, No Limit, 12 Hrs of RWS, B) 5 SCPF, 2000 TSC, 12 Hrs of RWS, C) 5 SCPF, 1750 TSC, 12 Hrs of RWS, D) 5 SCPF, 1000 TSC, 12 Hrs of RWS, E) 3 SCPF, No Limit, 12 Hrs of RWS, F) 5 SCPF, No Limit, 8 Hrs of RWS, G) 5 SCPF, 1250 TSC, 12 Hrs of RWS, H) 5 SCPF, 1500 TSC, 12 Hrs of RWS, I) 5 SCPF, 500 TSC, 12 Hrs of RWS.

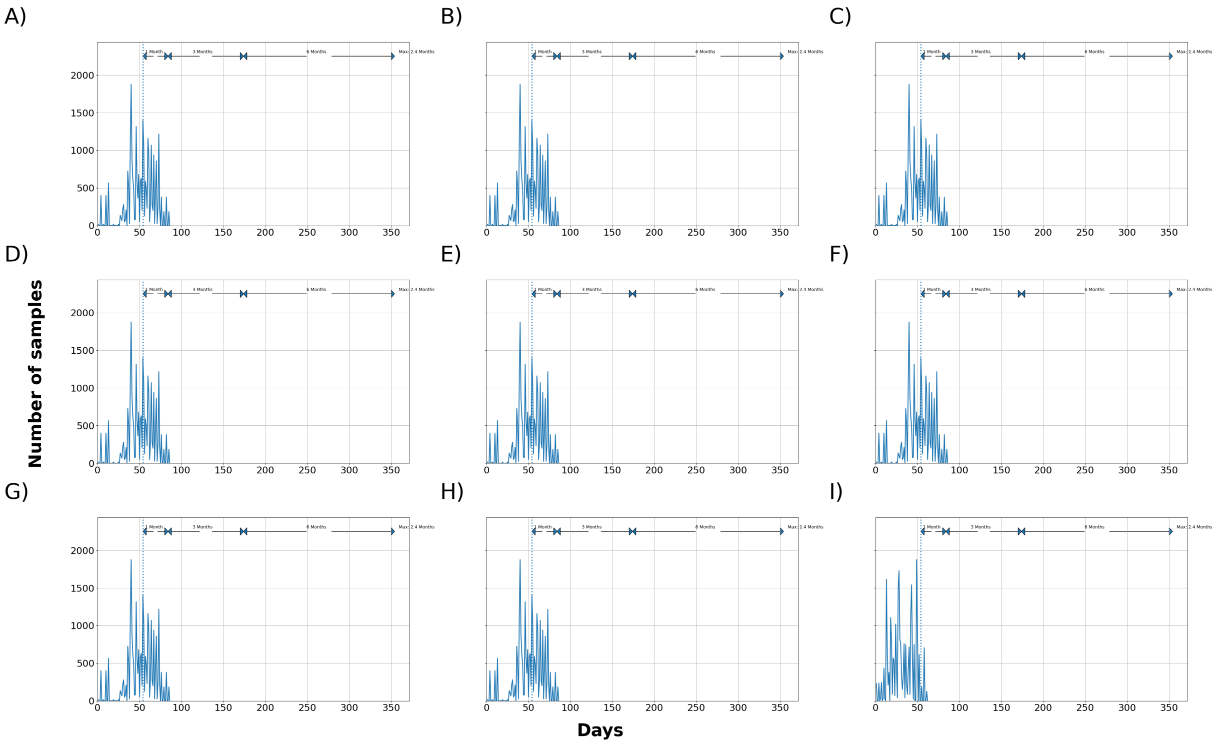


Figure 8: Median total number of samples waiting to be processed at the laboratory. 72 hours downtime and pool of 1 oral fluid sample. SCPF = Sampler collectors per farm, TSC = Trained sample collectors. A) 5 SCPF, No Limit, 12 Hrs of RWS, B) 5 SCPF, 2000 TSC, 12 Hrs of RWS, C) 5 SCPF, 1750 TSC, 12 Hrs of RWS, D) 5 SCPF, 1000 TSC, 12 Hrs of RWS, E) 3 SCPF, No Limit, 12 Hrs of RWS, F) 5 SCPF, No Limit, 8 Hrs of RWS, G) 5 SCPF, 1250 TSC, 12 Hrs of RWS, H) 5 SCPF, 1500 TSC, 12 Hrs of RWS, I) 5 SCPF, 500 TSC, 12 Hrs of RWS.

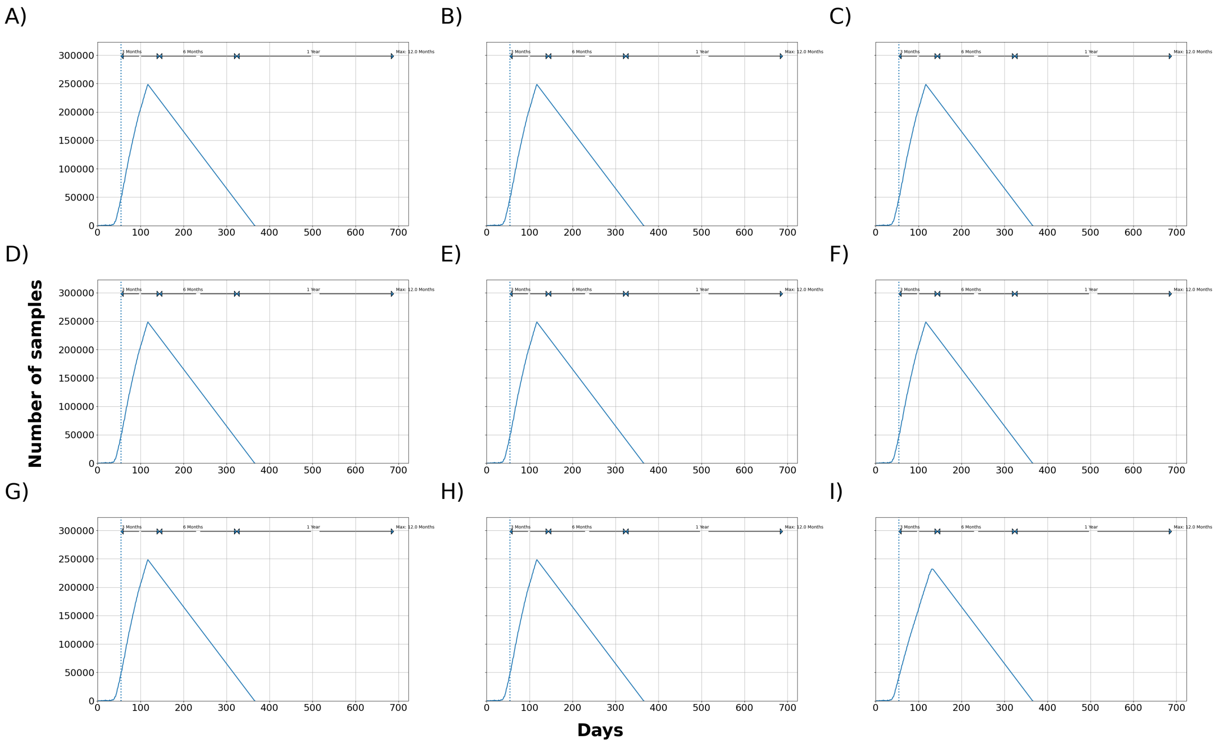


Figure 9: Maximum total number of samples waiting to be processed at the laboratory. 72 hours downtime and pool of 1 oral fluid sample. SCPF = Sampler collectors per farm, TSC = Trained sample collectors. A) 5 SCPF, No Limit, 12 Hrs of RWS, B) 5 SCPF, 2000 TSC, 12 Hrs of RWS, C) 5 SCPF, 1750 TSC, 12 Hrs of RWS, D) 5 SCPF, 1000 TSC, 12 Hrs of RWS, E) 3 SCPF, No Limit, 12 Hrs of RWS, F) 5 SCPF, No Limit, 8 Hrs of RWS, G) 5 SCPF, 1250 TSC, 12 Hrs of RWS, H) 5 SCPF, 1500 TSC, 12 Hrs of RWS, I) 5 SCPF, 500 TSC, 12 Hrs of RWS.

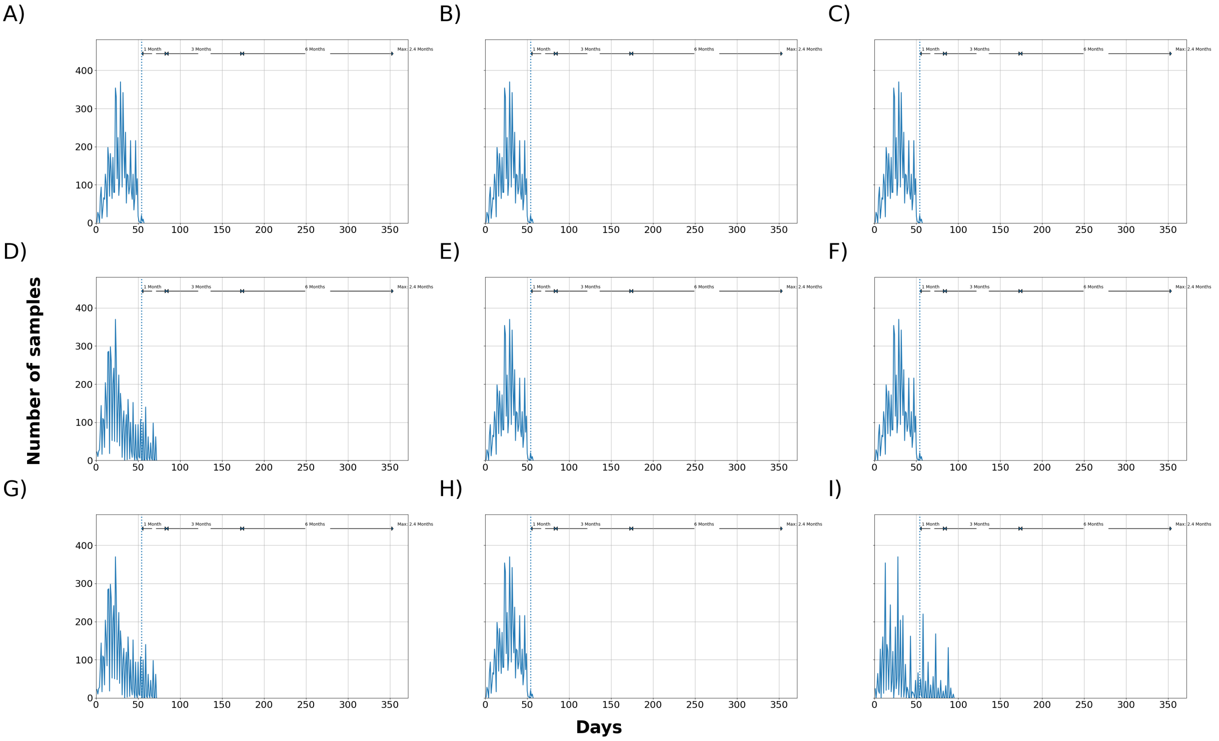


Figure 10: Median total number of samples waiting to be processed at the laboratory. 72 hours downtime and pool of 5 oral fluid samples. SCPF = Sampler collectors per farm, TSC = Trained sample collectors. A) 5 SCPF, No Limit, 12 Hrs of RWS, B) 5 SCPF, 2000 TSC, 12 Hrs of RWS, C) 5 SCPF, 1750 TSC, 12 Hrs of RWS, D) 5 SCPF, 1000 TSC, 12 Hrs of RWS, E) 3 SCPF, No Limit, 12 Hrs of RWS, F) 5 SCPF, No Limit, 8 Hrs of RWS, G) 5 SCPF, 1250 TSC, 12 Hrs of RWS, H) 5 SCPF, 1500 TSC, 12 Hrs of RWS, I) 5 SCPF, 500 TSC, 12 Hrs of RWS.

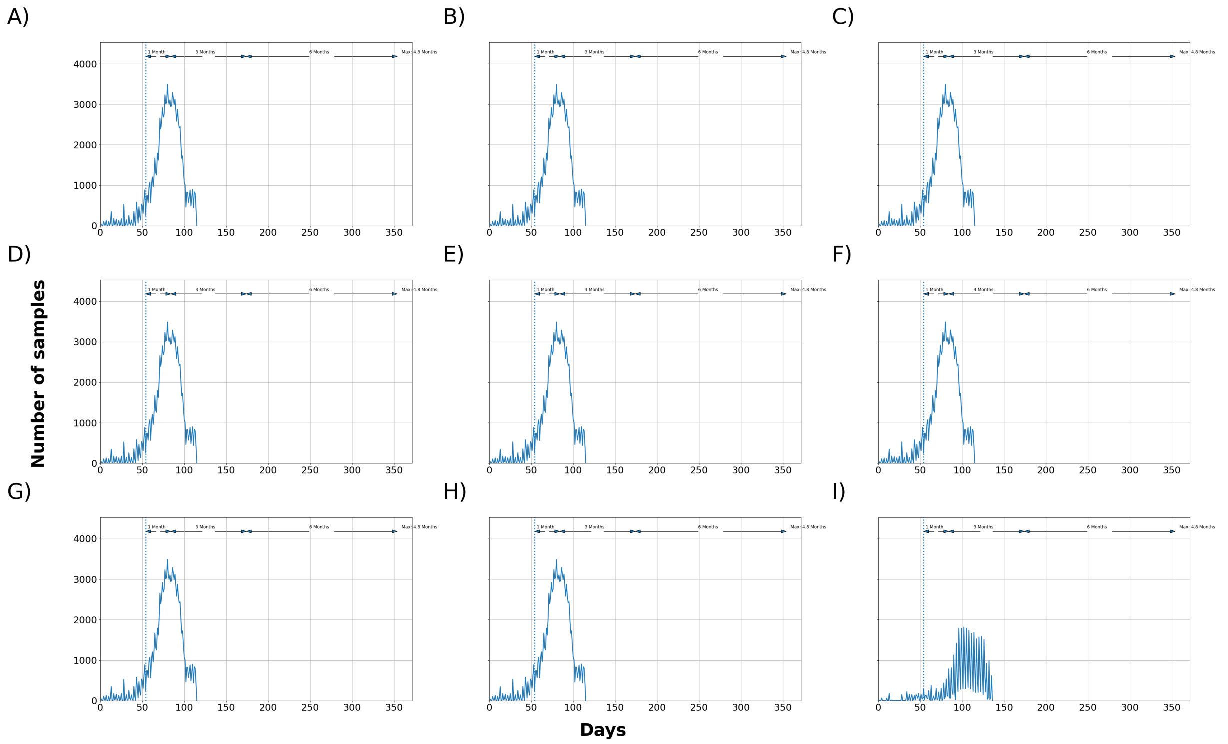


Figure 11: Maximum total number of samples waiting to be processed at the laboratory. 72 hours downtime and pool of 5 oral fluid samples. SCPF = Sampler collectors per farm, TSC = Trained sample collectors. A) 5 SCPF, No Limit, 12 Hrs of RWS, B) 5 SCPF, 2000 TSC, 12 Hrs of RWS, C) 5 SCPF, 1750 TSC, 12 Hrs of RWS, D) 5 SCPF, 1000 TSC, 12 Hrs of RWS, E) 3 SCPF, No Limit, 12 Hrs of RWS, F) 5 SCPF, No Limit, 8 Hrs of RWS, G) 5 SCPF, 1250 TSC, 12 Hrs of RWS, H) 5 SCPF, 1500 TSC, 12 Hrs of RWS, I) 5 SCPF, 500 TSC, 12 Hrs of RWS.

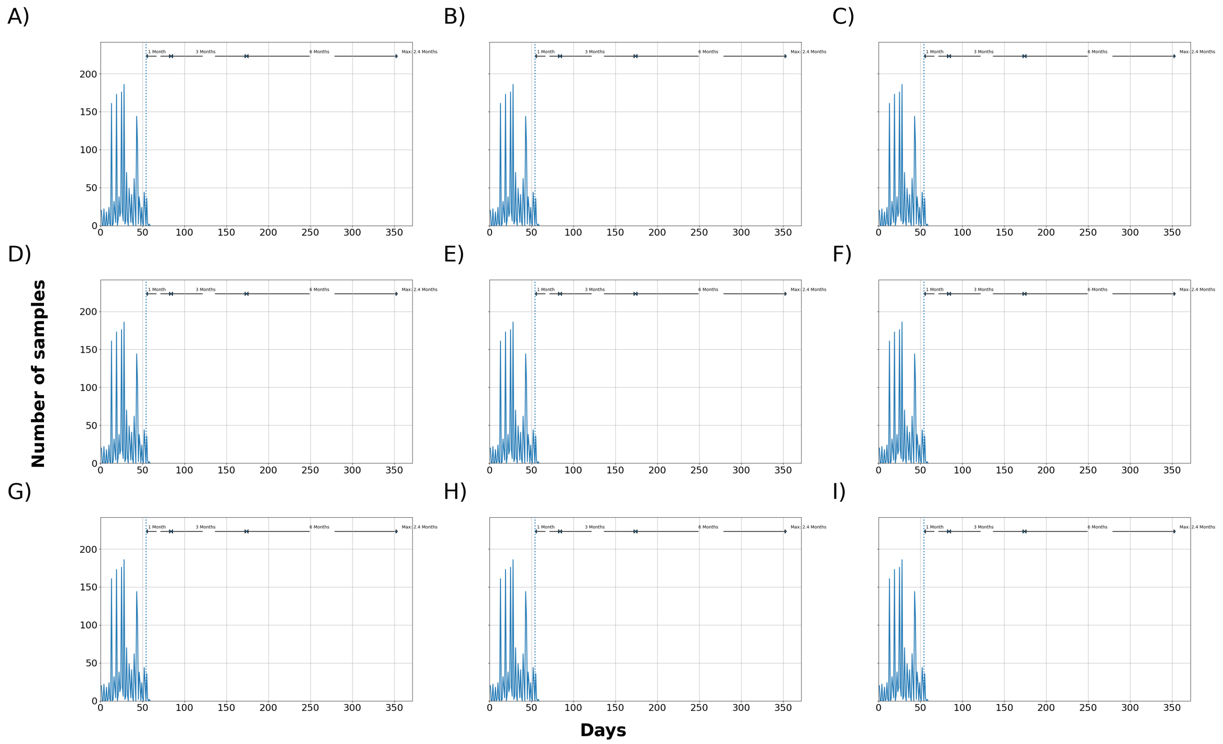


Figure 12: Median total number of samples waiting to be processed at the laboratory. 72 hours downtime and pool of 10 oral fluid samples. SCPF = Sampler collectors per farm, TSC = Trained sample collectors. A) 5 SCPF, No Limit, 12 Hrs of RWS, B) 5 SCPF, 2000 TSC, 12 Hrs of RWS, C) 5 SCPF, 1750 TSC, 12 Hrs of RWS, D) 5 SCPF, 1000 TSC, 12 Hrs of RWS, E) 3 SCPF, No Limit, 12 Hrs of RWS, F) 5 SCPF, No Limit, 8 Hrs of RWS, G) 5 SCPF, 1250 TSC, 12 Hrs of RWS, H) 5 SCPF, 1500 TSC, 12 Hrs of RWS, I) 5 SCPF, 500 TSC, 12 Hrs of RWS.

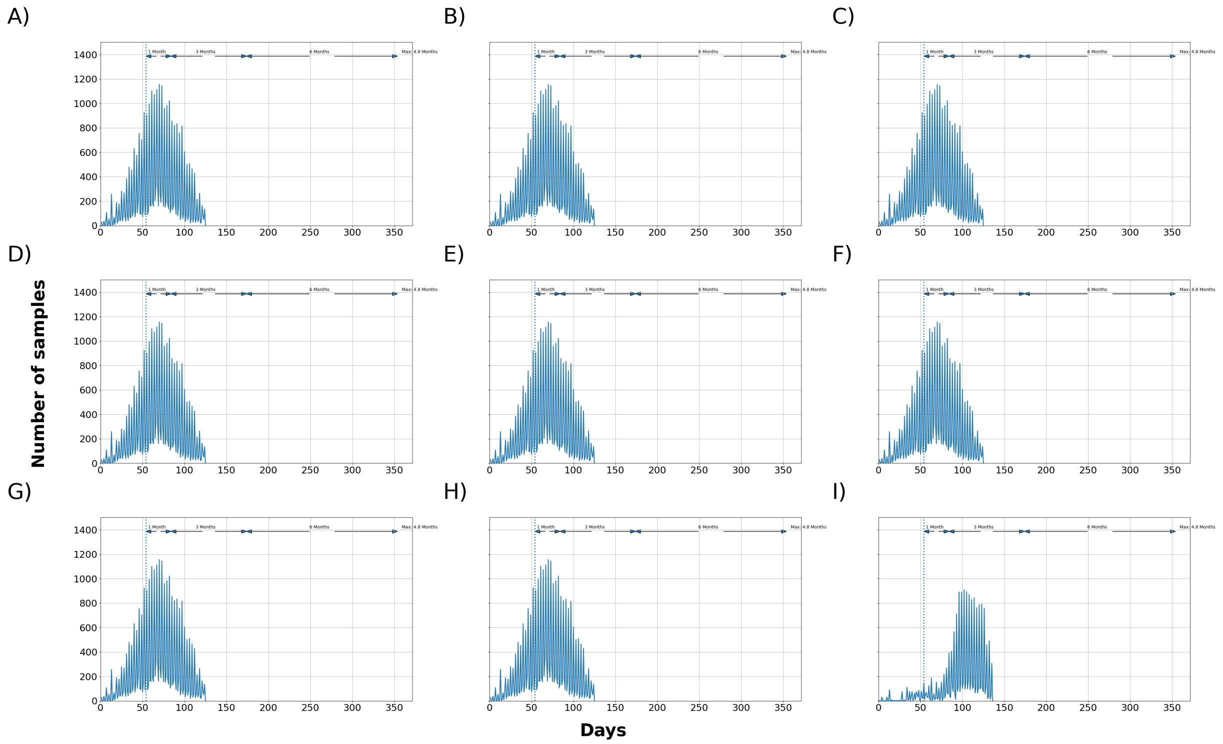


Figure 13: Maximum total number of samples waiting to be processed at the laboratory. 72 hours downtime and pool of 10 oral fluid samples. SCPF = Sampler collectors per farm, TSC = Trained sample collectors. A) 5 SCPF, No Limit, 12 Hrs of RWS, B) 5 SCPF, 2000 TSC, 12 Hrs of RWS, C) 5 SCPF, 1750 TSC, 12 Hrs of RWS, D) 5 SCPF, 1000 TSC, 12 Hrs of RWS, E) 3 SCPF, No Limit, 12 Hrs of RWS, F) 5 SCPF, No Limit, 8 Hrs of RWS, G) 5 SCPF, 1250 TSC, 12 Hrs of RWS, H) 5 SCPF, 1500 TSC, 12 Hrs of RWS, I) 5 SCPF, 500 TSC, 12 Hrs of RWS.

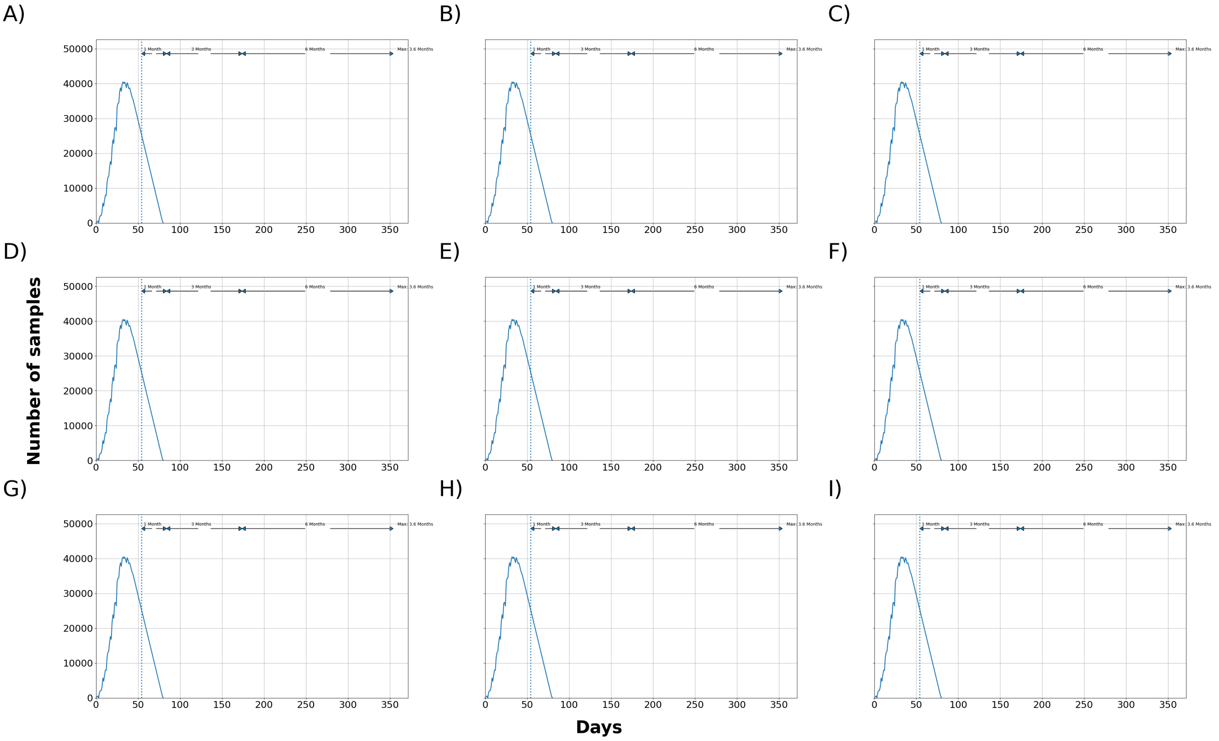


Figure 14: Median total number of samples waiting to be processed at the laboratory. 24 hours downtime and pool of 1 blood sample. SCPF = Sampler collectors per farm, TSC = Trained sample collectors. A) 5 SCPF, No Limit, 12 Hrs of RWS, B) 5 SCPF, 2000 TSC, 12 Hrs of RWS, C) 5 SCPF, 1750 TSC, 12 Hrs of RWS, D) 5 SCPF, 1000 TSC, 12 Hrs of RWS, E) 3 SCPF, No Limit, 12 Hrs of RWS, F) 5 SCPF, No Limit, 8 Hrs of RWS, G) 5 SCPF, 1250 TSC, 12 Hrs of RWS, H) 5 SCPF, 1500 TSC, 12 Hrs of RWS, I) 5 SCPF, 500 TSC, 12 Hrs of RWS.

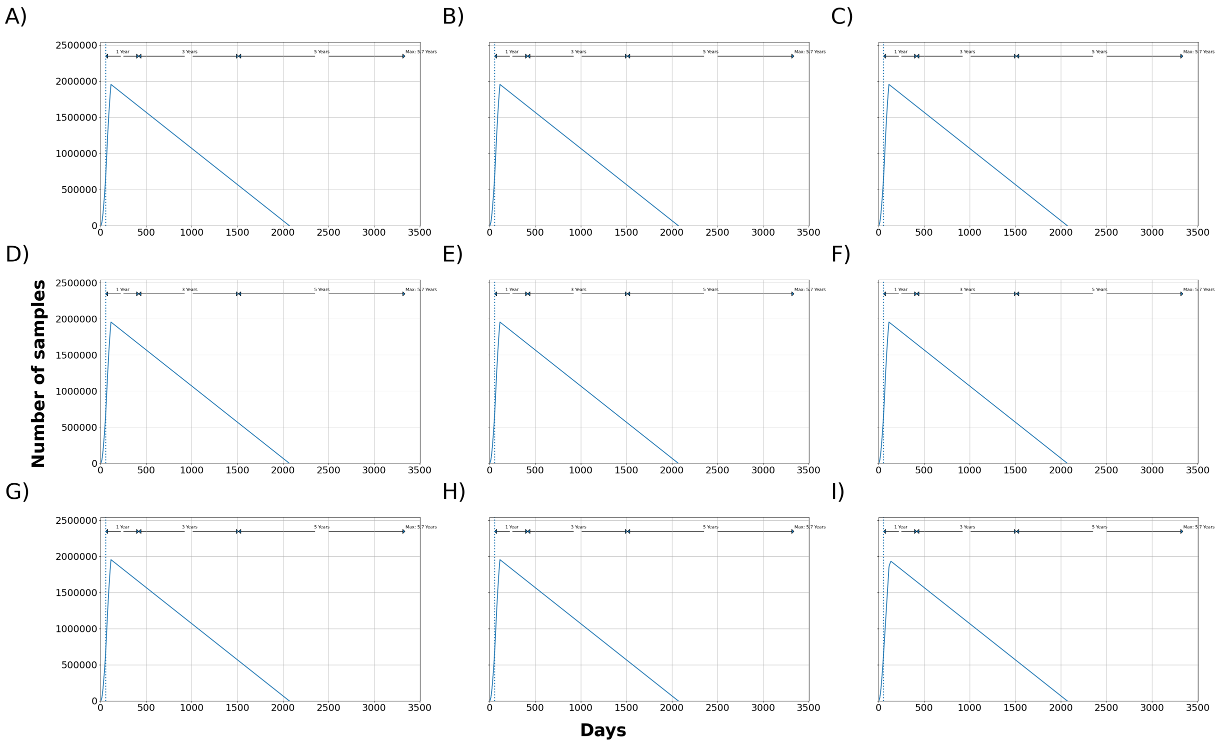


Figure 15: Maximum total number of samples waiting to be processed at the laboratory. 24 hours downtime and pool of 1 blood sample. SCPF = Sampler collectors per farm, TSC = Trained sample collectors. A) 5 SCPF, No Limit, 12 Hrs of RWS, B) 5 SCPF, 2000 TSC, 12 Hrs of RWS, C) 5 SCPF, 1750 TSC, 12 Hrs of RWS, D) 5 SCPF, 1000 TSC, 12 Hrs of RWS, E) 3 SCPF, No Limit, 12 Hrs of RWS, F) 5 SCPF, No Limit, 8 Hrs of RWS, G) 5 SCPF, 1250 TSC, 12 Hrs of RWS, H) 5 SCPF, 1500 TSC, 12 Hrs of RWS, I) 5 SCPF, 500 TSC, 12 Hrs of RWS.

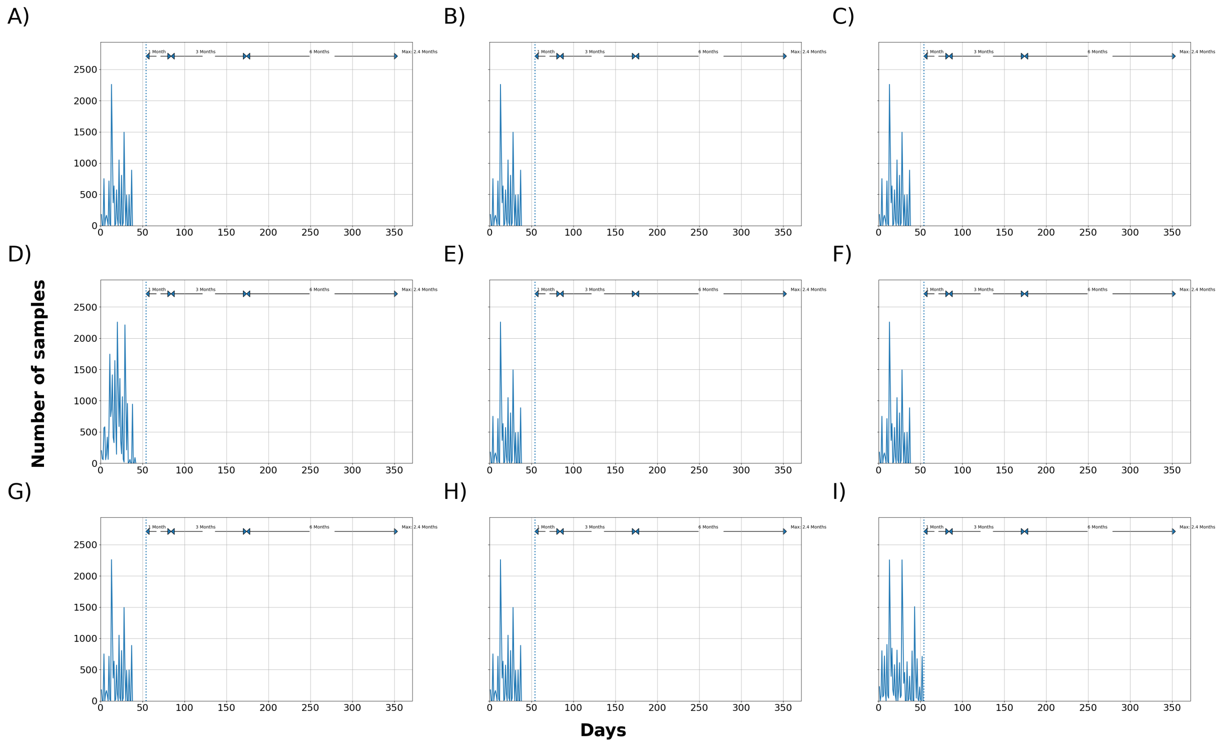


Figure 16: Median total number of samples waiting to be processed at the laboratory. 24 hours downtime and pool of 5 blood samples. SCPF = Sampler collectors per farm, TSC = Trained sample collectors. A) 5 SCPF, No Limit, 12 Hrs of RWS, B) 5 SCPF, 2000 TSC, 12 Hrs of RWS, C) 5 SCPF, 1750 TSC, 12 Hrs of RWS, D) 5 SCPF, 1000 TSC, 12 Hrs of RWS, E) 3 SCPF, No Limit, 12 Hrs of RWS, F) 5 SCPF, No Limit, 8 Hrs of RWS, G) 5 SCPF, 1250 TSC, 12 Hrs of RWS, H) 5 SCPF, 1500 TSC, 12 Hrs of RWS, I) 5 SCPF, 500 TSC, 12 Hrs of RWS.

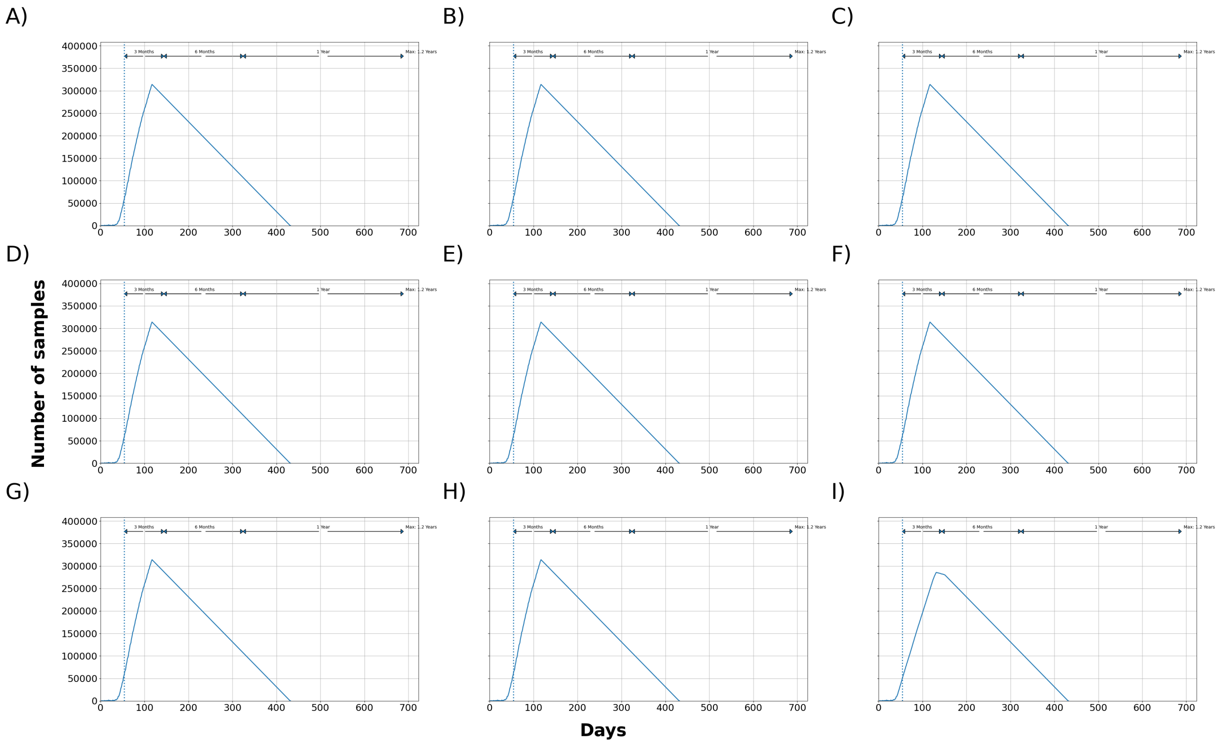


Figure 17: Maximum total number of samples waiting to be processed at the laboratory. 24 hours downtime and pool of 5 blood samples. SCPF = Sampler collectors per farm, TSC = Trained sample collectors. A) 5 SCPF, No Limit, 12 Hrs of RWS, B) 5 SCPF, 2000 TSC, 12 Hrs of RWS, C) 5 SCPF, 1750 TSC, 12 Hrs of RWS, D) 5 SCPF, 1000 TSC, 12 Hrs of RWS, E) 3 SCPF, No Limit, 12 Hrs of RWS, F) 5 SCPF, No Limit, 8 Hrs of RWS, G) 5 SCPF, 1250 TSC, 12 Hrs of RWS, H) 5 SCPF, 1500 TSC, 12 Hrs of RWS, I) 5 SCPF, 500 TSC, 12 Hrs of RWS.

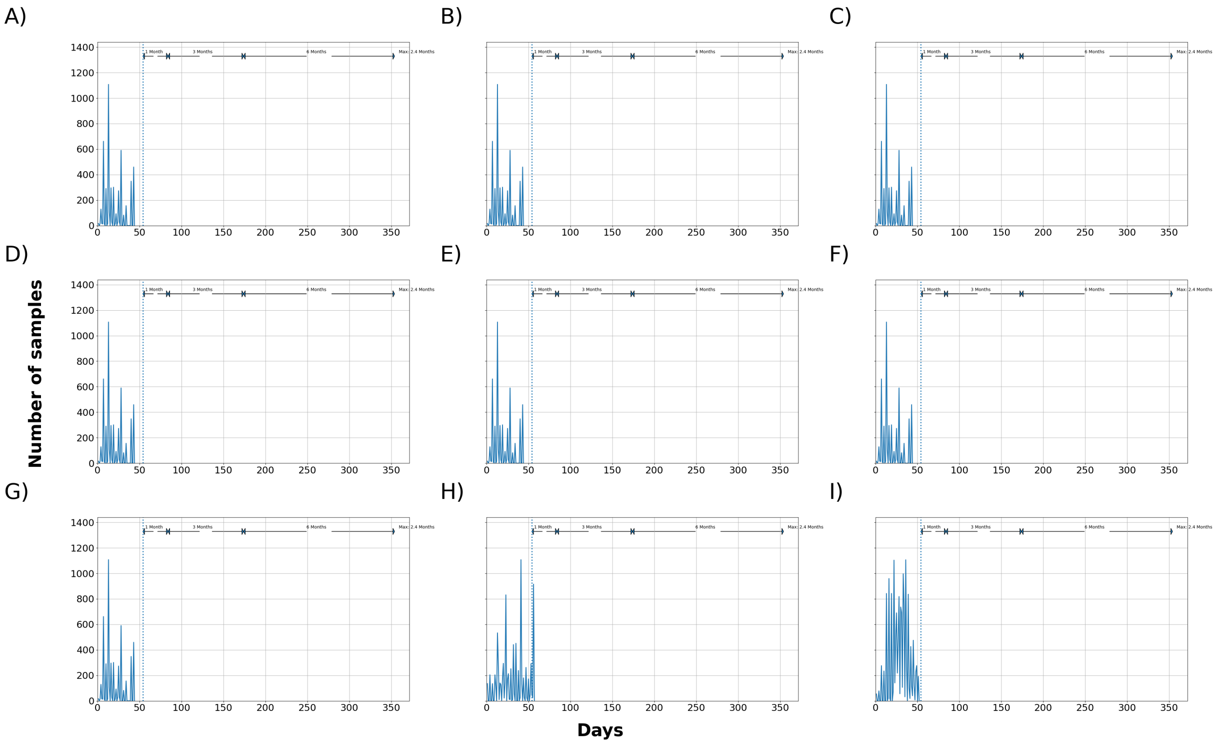


Figure 18: Median total number of samples waiting to be processed at the laboratory. 24 hours downtime and pool of 10 blood samples. SCPF = Sampler collectors per farm, TSC = Trained sample collectors. A) 5 SCPF, No Limit, 12 Hrs of RWS, B) 5 SCPF, 2000 TSC, 12 Hrs of RWS, C) 5 SCPF, 1750 TSC, 12 Hrs of RWS, D) 5 SCPF, 1000 TSC, 12 Hrs of RWS, E) 3 SCPF, No Limit, 12 Hrs of RWS, F) 5 SCPF, No Limit, 8 Hrs of RWS, G) 5 SCPF, 1250 TSC, 12 Hrs of RWS, H) 5 SCPF, 1500 TSC, 12 Hrs of RWS, I) 5 SCPF, 500 TSC, 12 Hrs of RWS.

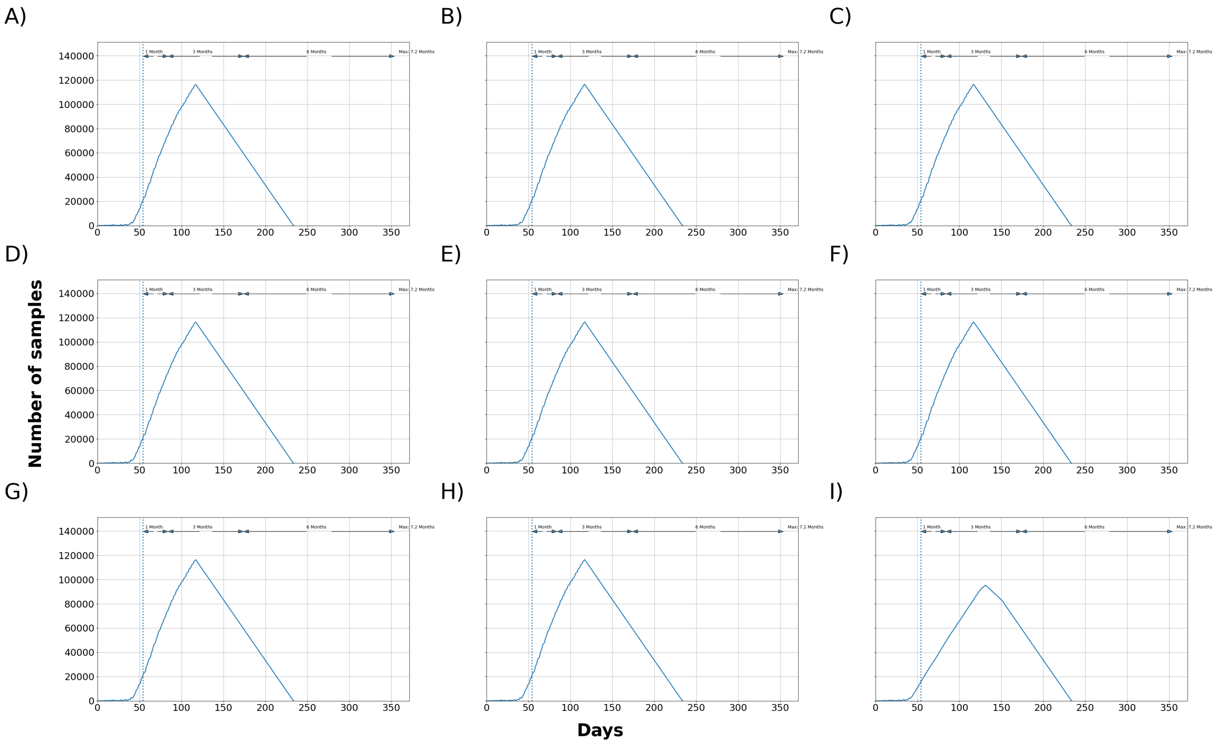


Figure 19: Maximum total number of samples waiting to be processed at the laboratory. 24 hours downtime and pool of 10 blood samples. SCPF = Sampler collectors per farm, TSC = Trained sample collectors. A) 5 SCPF, No Limit, 12 Hrs of RWS, B) 5 SCPF, 2000 TSC, 12 Hrs of RWS, C) 5 SCPF, 1750 TSC, 12 Hrs of RWS, D) 5 SCPF, 1000 TSC, 12 Hrs of RWS, E) 3 SCPF, No Limit, 12 Hrs of RWS, F) 5 SCPF, No Limit, 8 Hrs of RWS, G) 5 SCPF, 1250 TSC, 12 Hrs of RWS, H) 5 SCPF, 1500 TSC, 12 Hrs of RWS, I) 5 SCPF, 500 TSC, 12 Hrs of RWS.

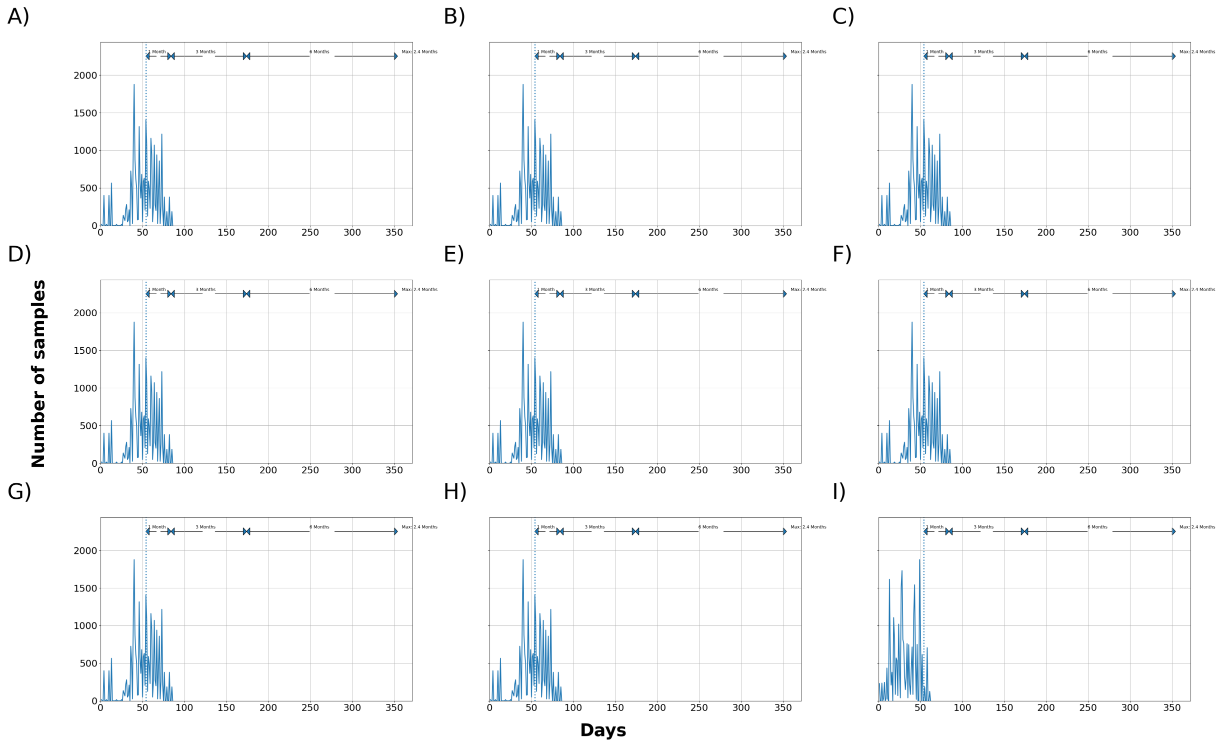


Figure 20: Median total number of samples waiting to be processed at the laboratory. 24 hours downtime and pool of 1 oral fluid sample. SCPF = Sampler collectors per farm, TSC = Trained sample collectors. A) 5 SCPF, No Limit, 12 Hrs of RWS, B) 5 SCPF, 2000 TSC, 12 Hrs of RWS, C) 5 SCPF, 1750 TSC, 12 Hrs of RWS, D) 5 SCPF, 1000 TSC, 12 Hrs of RWS, E) 3 SCPF, No Limit, 12 Hrs of RWS, F) 5 SCPF, No Limit, 8 Hrs of RWS, G) 5 SCPF, 1250 TSC, 12 Hrs of RWS, H) 5 SCPF, 1500 TSC, 12 Hrs of RWS, I) 5 SCPF, 500 TSC, 12 Hrs of RWS.

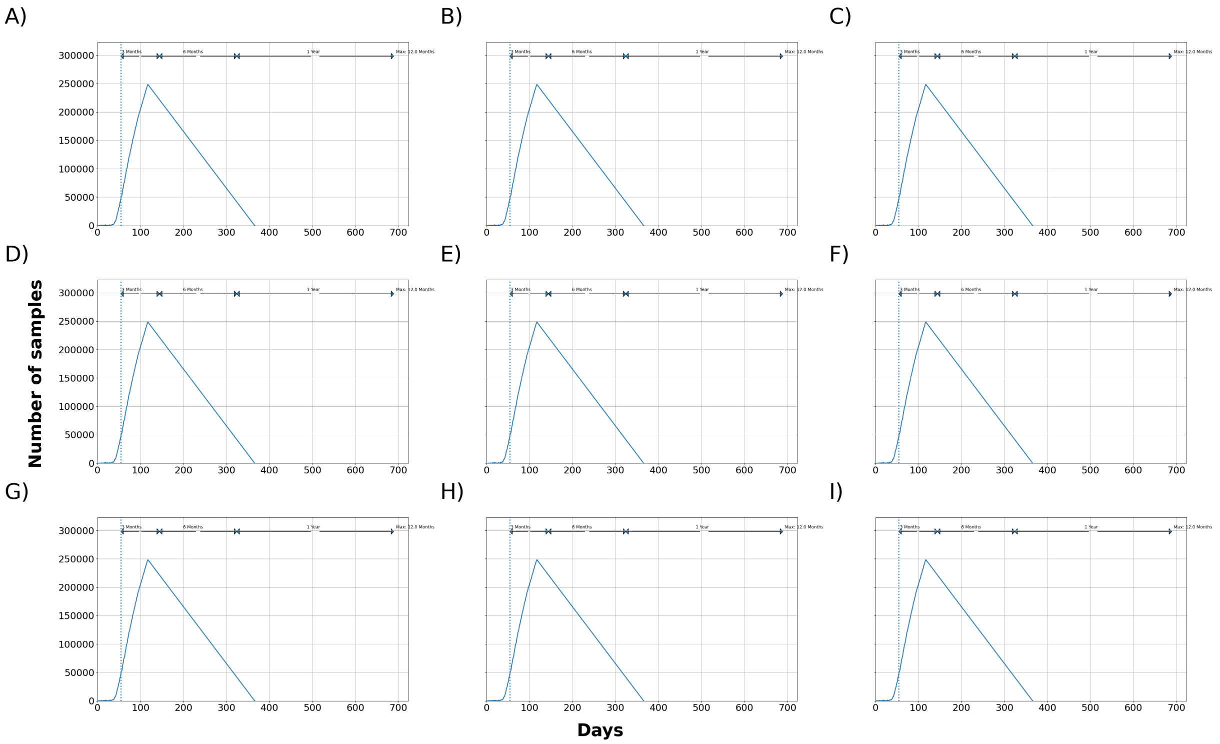


Figure 21: Maximum total number of samples waiting to be processed at the laboratory. 24 hours downtime and pool of 1 oral fluid sample. SCPF = Sampler collectors per farm, TSC = Trained sample collectors. A) 5 SCPF, No Limit, 12 Hrs of RWS, B) 5 SCPF, 2000 TSC, 12 Hrs of RWS, C) 5 SCPF, 1750 TSC, 12 Hrs of RWS, D) 5 SCPF, 1000 TSC, 12 Hrs of RWS, E) 3 SCPF, No Limit, 12 Hrs of RWS, F) 5 SCPF, No Limit, 8 Hrs of RWS, G) 5 SCPF, 1250 TSC, 12 Hrs of RWS, H) 5 SCPF, 1500 TSC, 12 Hrs of RWS, I) 5 SCPF, 500 TSC, 12 Hrs of RWS.

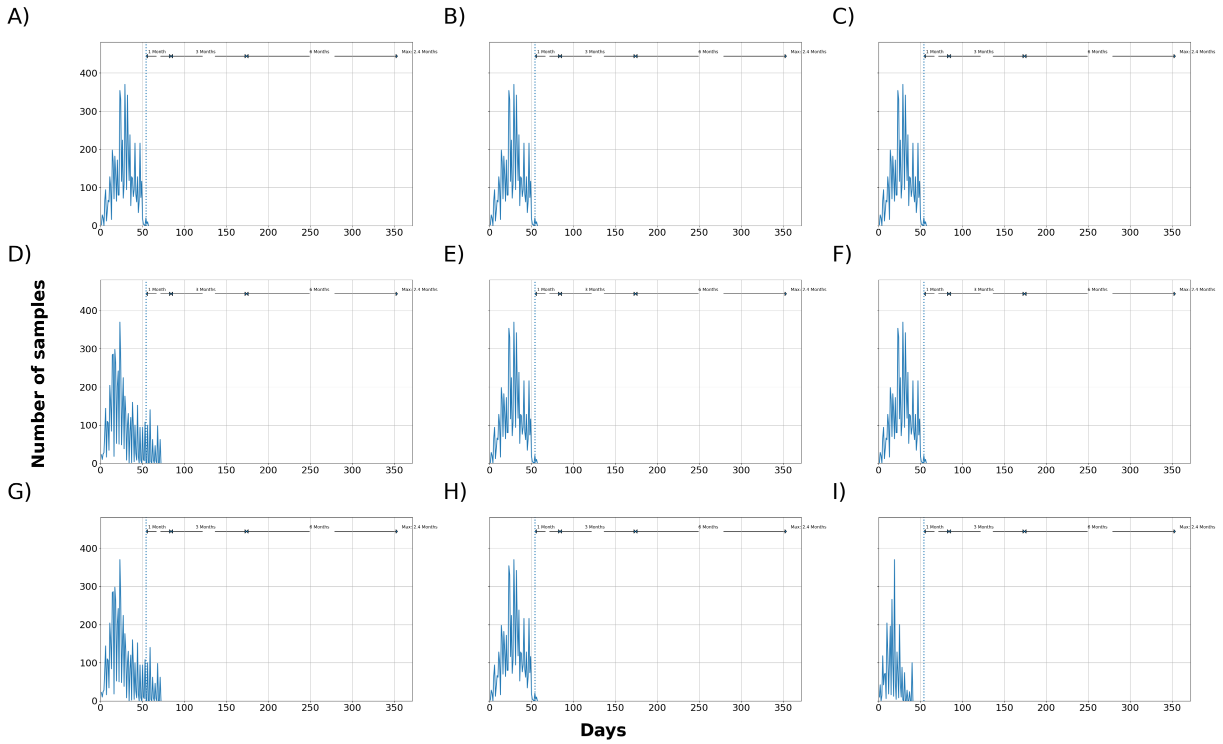


Figure 22: Median total number of samples waiting to be processed at the laboratory. 24 hours downtime and pool of 5 oral fluid samples. SCPF = Sampler collectors per farm, TSC = Trained sample collectors. A) 5 SCPF, No Limit, 12 Hrs of RWS, B) 5 SCPF, 2000 TSC, 12 Hrs of RWS, C) 5 SCPF, 1750 TSC, 12 Hrs of RWS, D) 5 SCPF, 1000 TSC, 12 Hrs of RWS, E) 3 SCPF, No Limit, 12 Hrs of RWS, F) 5 SCPF, No Limit, 8 Hrs of RWS, G) 5 SCPF, 1250 TSC, 12 Hrs of RWS, H) 5 SCPF, 1500 TSC, 12 Hrs of RWS, I) 5 SCPF, 500 TSC, 12 Hrs of RWS.

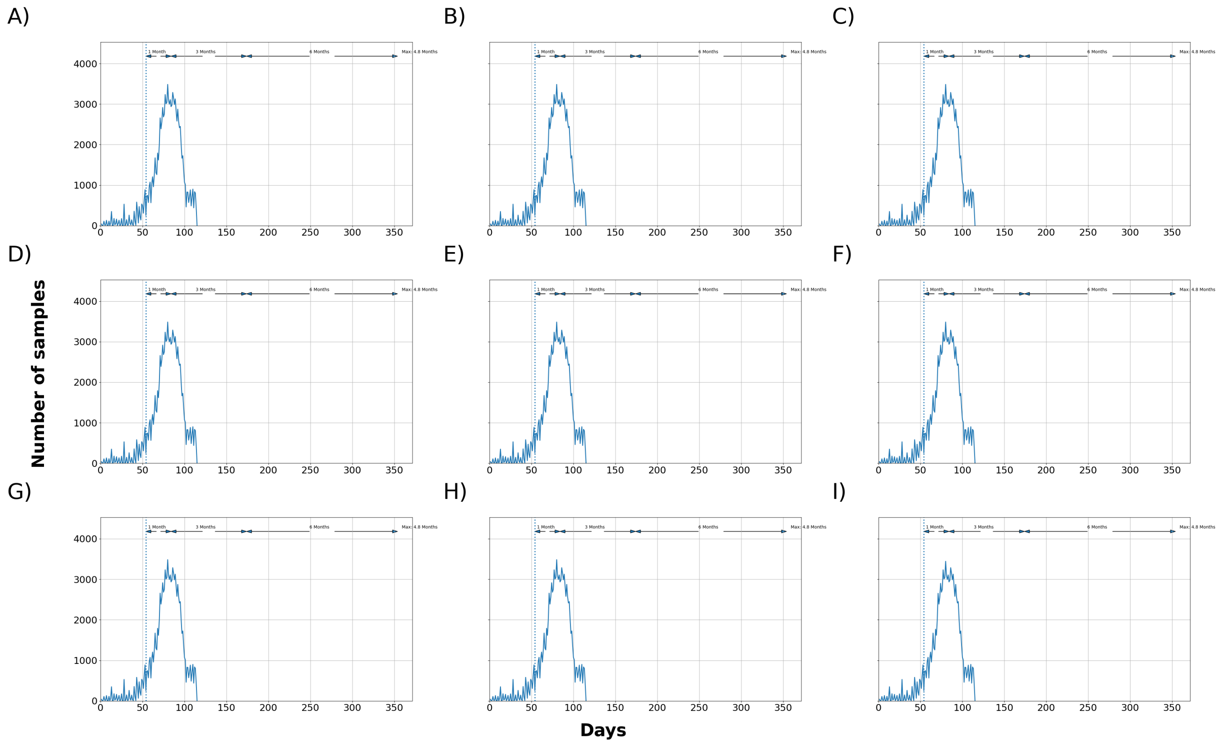


Figure 23: Maximum total number of samples waiting to be processed at the laboratory. 24 hours downtime and pool of 5 oral fluid samples. SCPF = Sampler collectors per farm, TSC = Trained sample collectors. A) 5 SCPF, No Limit, 12 Hrs of RWS, B) 5 SCPF, 2000 TSC, 12 Hrs of RWS, C) 5 SCPF, 1750 TSC, 12 Hrs of RWS, D) 5 SCPF, 1000 TSC, 12 Hrs of RWS, E) 3 SCPF, No Limit, 12 Hrs of RWS, F) 5 SCPF, No Limit, 8 Hrs of RWS, G) 5 SCPF, 1250 TSC, 12 Hrs of RWS, H) 5 SCPF, 1500 TSC, 12 Hrs of RWS, I) 5 SCPF, 500 TSC, 12 Hrs of RWS.

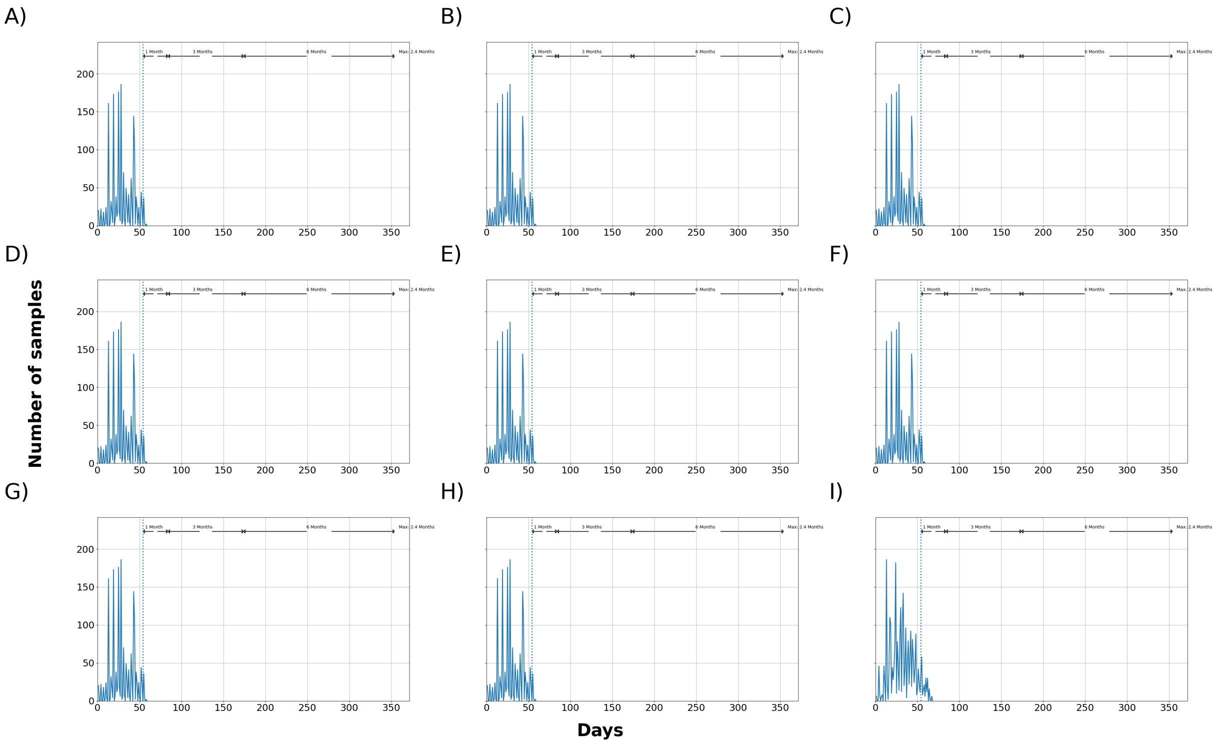


Figure 24: Median total number of samples waiting to be processed at the laboratory. 24 hours downtime and pool of 10 oral fluid samples. SCPF = Sampler collectors per farm, TSC = Trained sample collectors. A) 5 SCPF, No Limit, 12 Hrs of RWS, B) 5 SCPF, 2000 TSC, 12 Hrs of RWS, C) 5 SCPF, 1750 TSC, 12 Hrs of RWS, D) 5 SCPF, 1000 TSC, 12 Hrs of RWS, E) 3 SCPF, No Limit, 12 Hrs of RWS, F) 5 SCPF, No Limit, 8 Hrs of RWS, G) 5 SCPF, 1250 TSC, 12 Hrs of RWS, H) 5 SCPF, 1500 TSC, 12 Hrs of RWS, I) 5 SCPF, 500 TSC, 12 Hrs of RWS.

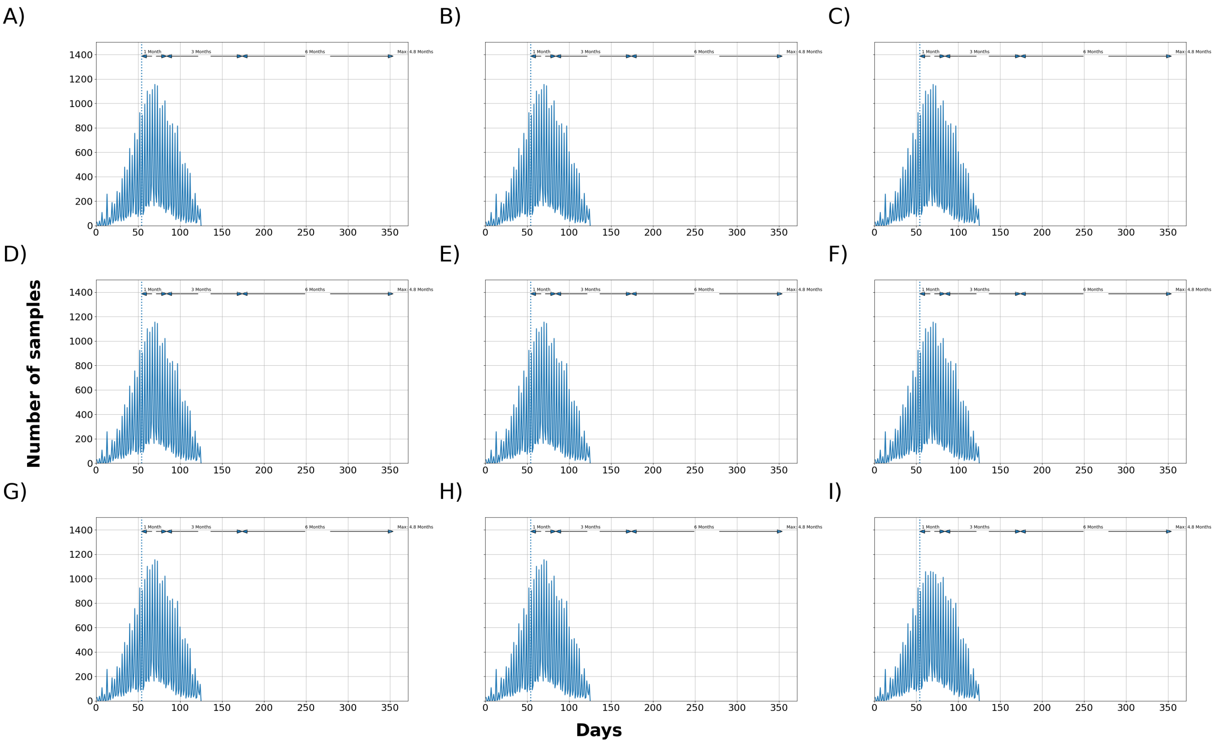


Figure 25: Maximum total number of samples waiting to be processed at the laboratory. 24 hours downtime and pool of 10 oral fluid samples. SCPF = Sampler collectors per farm, TSC = Trained sample collectors. A) 5 SCPF, No Limit, 12 Hrs of RWS, B) 5 SCPF, 2000 TSC, 12 Hrs of RWS, C) 5 SCPF, 1750 TSC, 12 Hrs of RWS, D) 5 SCPF, 1000 TSC, 12 Hrs of RWS, E) 3 SCPF, No Limit, 12 Hrs of RWS, F) 5 SCPF, No Limit, 8 Hrs of RWS, G) 5 SCPF, 1250 TSC, 12 Hrs of RWS, H) 5 SCPF, 1500 TSC, 12 Hrs of RWS, I) 5 SCPF, 500 TSC, 12 Hrs of RWS.

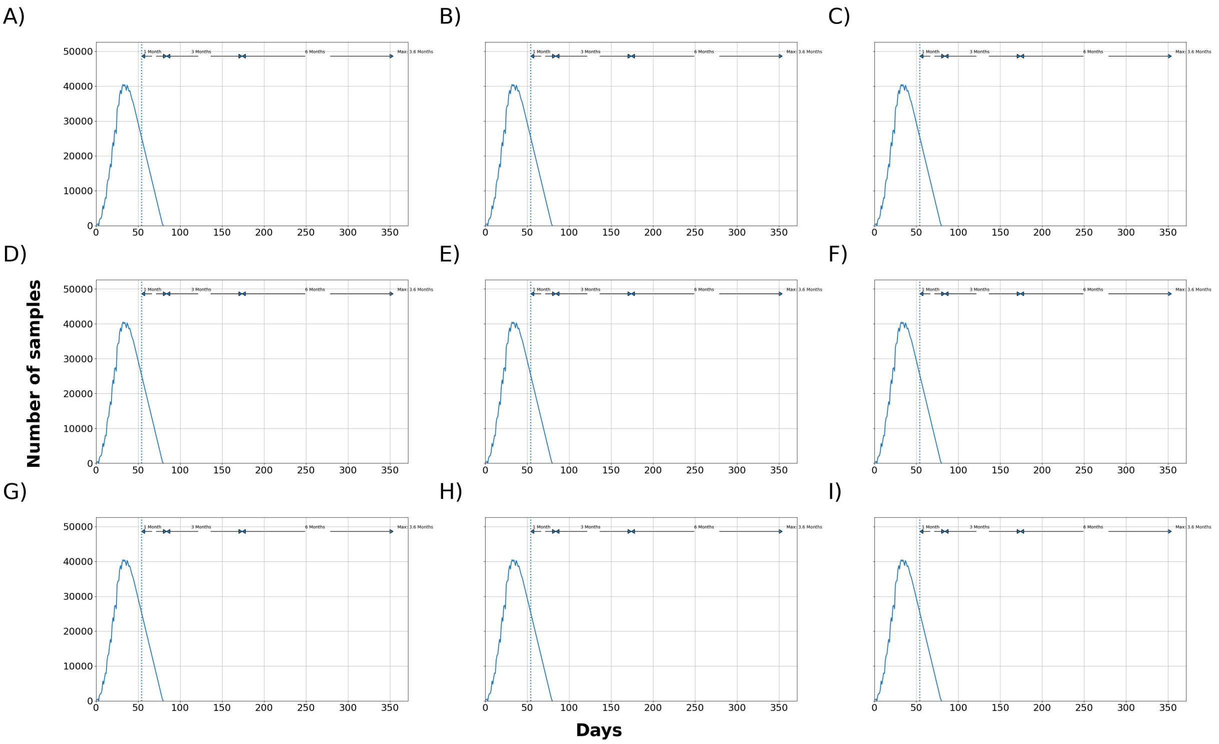


Figure 26: Median total number of samples waiting to be processed at the laboratory. 0 hours downtime and pool of 1 blood sample. SCPF = Sampler collectors per farm, TSC = Trained sample collectors. A) 5 SCPF, No Limit, 12 Hrs of RWS, B) 5 SCPF, 2000 TSC, 12 Hrs of RWS, C) 5 SCPF, 1750 TSC, 12 Hrs of RWS, D) 5 SCPF, 1000 TSC, 12 Hrs of RWS, E) 3 SCPF, No Limit, 12 Hrs of RWS, F) 5 SCPF, No Limit, 8 Hrs of RWS, G) 5 SCPF, 1250 TSC, 12 Hrs of RWS, H) 5 SCPF, 1500 TSC, 12 Hrs of RWS, I) 5 SCPF, 500 TSC, 12 Hrs of RWS.

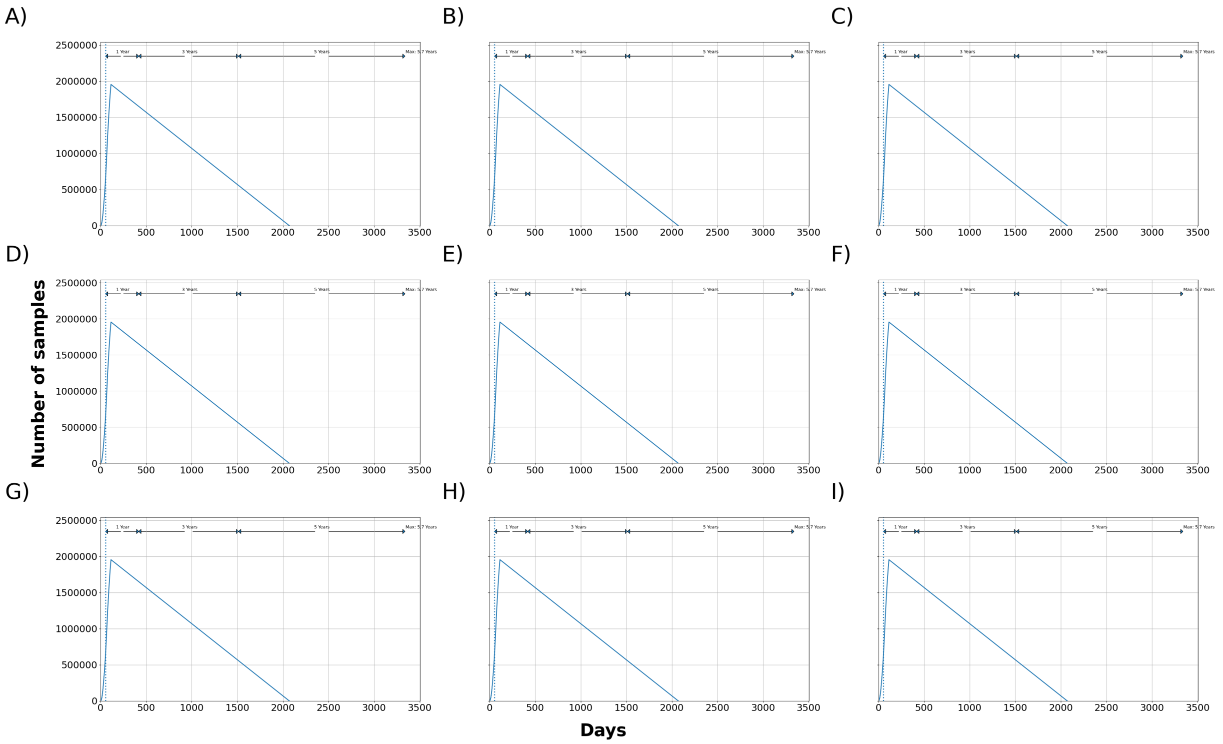


Figure 27: Maximum total number of samples waiting to be processed at the laboratory. 0 hours downtime and pool of 1 blood sample. SCPF = Sampler collectors per farm, TSC = Trained sample collectors. A) 5 SCPF, No Limit, 12 Hrs of RWS, B) 5 SCPF, 2000 TSC, 12 Hrs of RWS, C) 5 SCPF, 1750 TSC, 12 Hrs of RWS, D) 5 SCPF, 1000 TSC, 12 Hrs of RWS, E) 3 SCPF, No Limit, 12 Hrs of RWS, F) 5 SCPF, No Limit, 8 Hrs of RWS, G) 5 SCPF, 1250 TSC, 12 Hrs of RWS, H) 5 SCPF, 1500 TSC, 12 Hrs of RWS, I) 5 SCPF, 500 TSC, 12 Hrs of RWS.

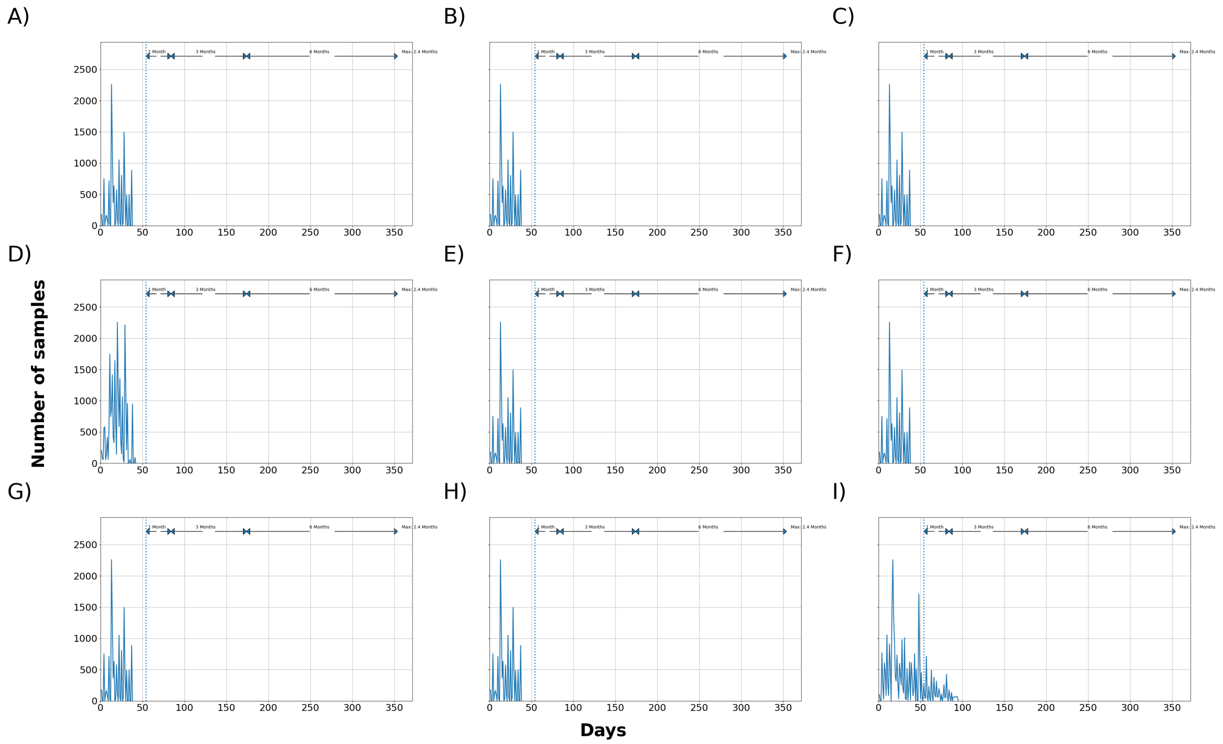


Figure 28: Median total number of samples waiting to be processed at the laboratory. 0 hours downtime and pool of 5 blood samples. SCPF = Sampler collectors per farm, TSC = Trained sample collectors. A) 5 SCPF, No Limit, 12 Hrs of RWS, B) 5 SCPF, 2000 TSC, 12 Hrs of RWS, C) 5 SCPF, 1750 TSC, 12 Hrs of RWS, D) 5 SCPF, 1000 TSC, 12 Hrs of RWS, E) 3 SCPF, No Limit, 12 Hrs of RWS, F) 5 SCPF, No Limit, 8 Hrs of RWS, G) 5 SCPF, 1250 TSC, 12 Hrs of RWS, H) 5 SCPF, 1500 TSC, 12 Hrs of RWS, I) 5 SCPF, 500 TSC, 12 Hrs of RWS.

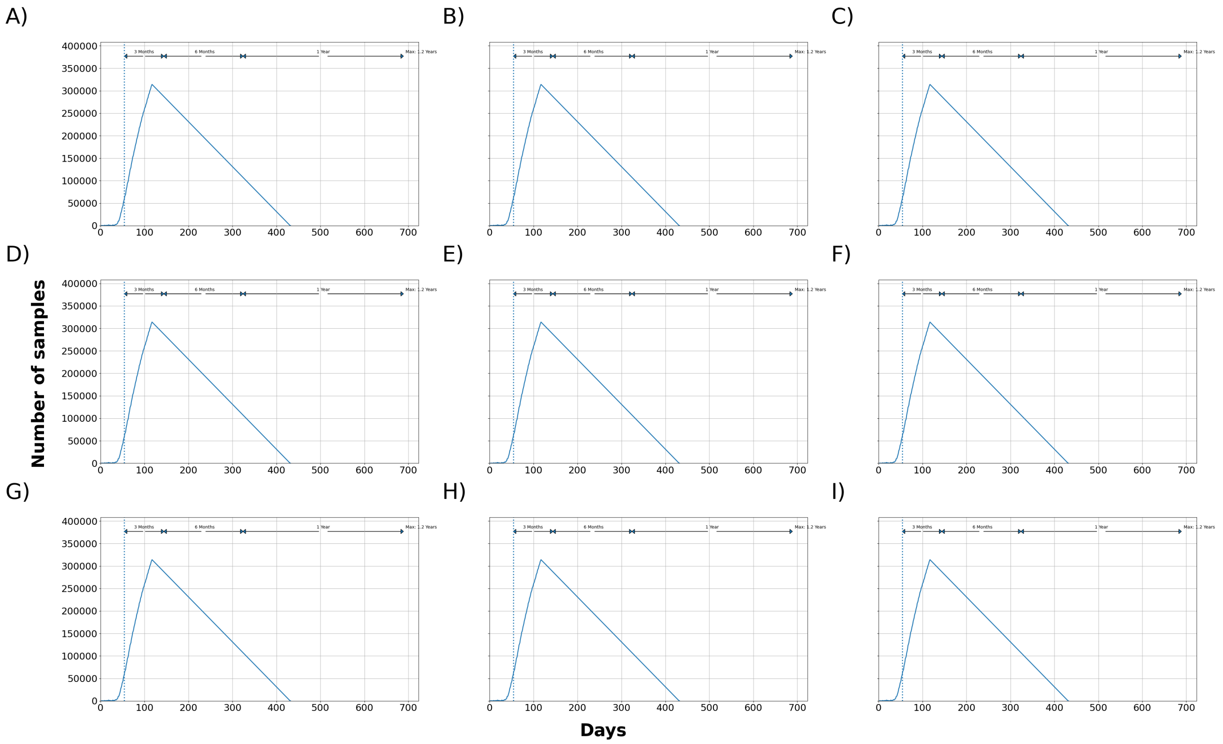


Figure 29: Maximum total number of samples waiting to be processed at the laboratory. 0 hours downtime and pool of 5 blood samples. SCPF = Sampler collectors per farm, TSC = Trained sample collectors. A) 5 SCPF, No Limit, 12 Hrs of RWS, B) 5 SCPF, 2000 TSC, 12 Hrs of RWS, C) 5 SCPF, 1750 TSC, 12 Hrs of RWS, D) 5 SCPF, 1000 TSC, 12 Hrs of RWS, E) 3 SCPF, No Limit, 12 Hrs of RWS, F) 5 SCPF, No Limit, 8 Hrs of RWS, G) 5 SCPF, 1250 TSC, 12 Hrs of RWS, H) 5 SCPF, 1500 TSC, 12 Hrs of RWS, I) 5 SCPF, 500 TSC, 12 Hrs of RWS.

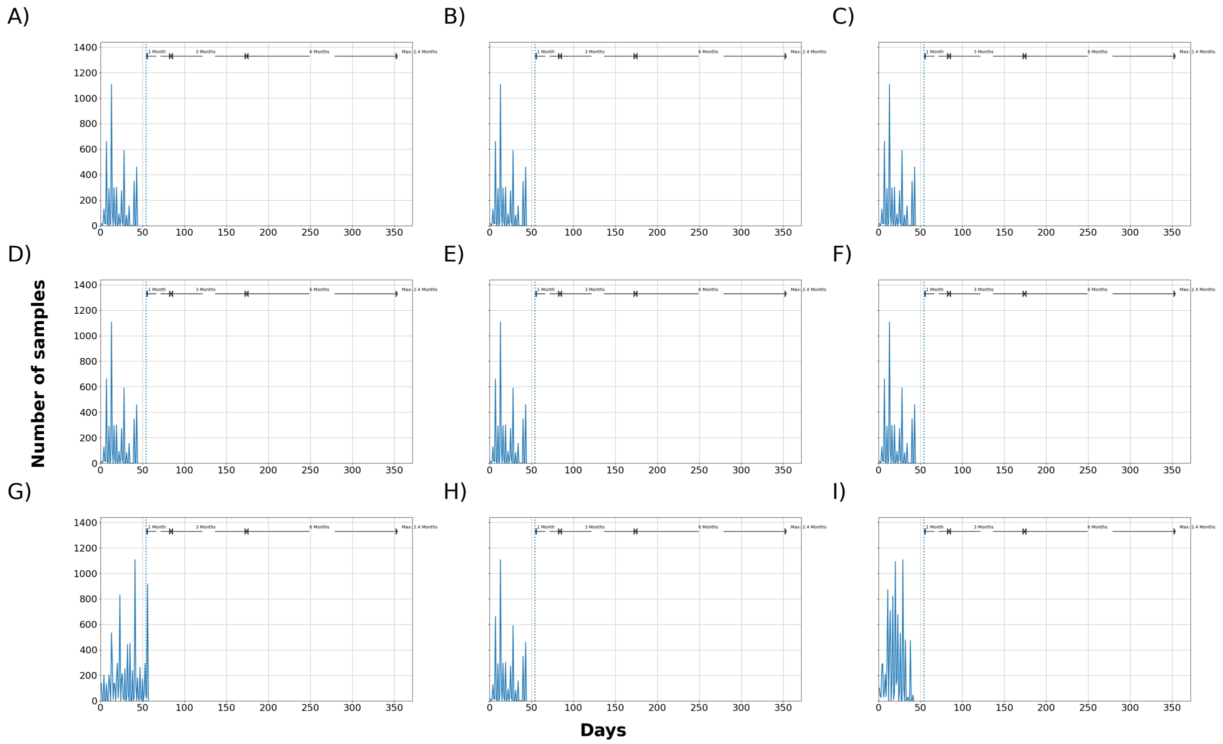


Figure 30: Median total number of samples waiting to be processed at the laboratory. 0 hours downtime and pool of 10 blood samples. SCPF = Sampler collectors per farm, TSC = Trained sample collectors. A) 5 SCPF, No Limit, 12 Hrs of RWS, B) 5 SCPF, 2000 TSC, 12 Hrs of RWS, C) 5 SCPF, 1750 TSC, 12 Hrs of RWS, D) 5 SCPF, 1000 TSC, 12 Hrs of RWS, E) 3 SCPF, No Limit, 12 Hrs of RWS, F) 5 SCPF, No Limit, 8 Hrs of RWS, G) 5 SCPF, 1250 TSC, 12 Hrs of RWS, H) 5 SCPF, 1500 TSC, 12 Hrs of RWS, I) 5 SCPF, 500 TSC, 12 Hrs of RWS.

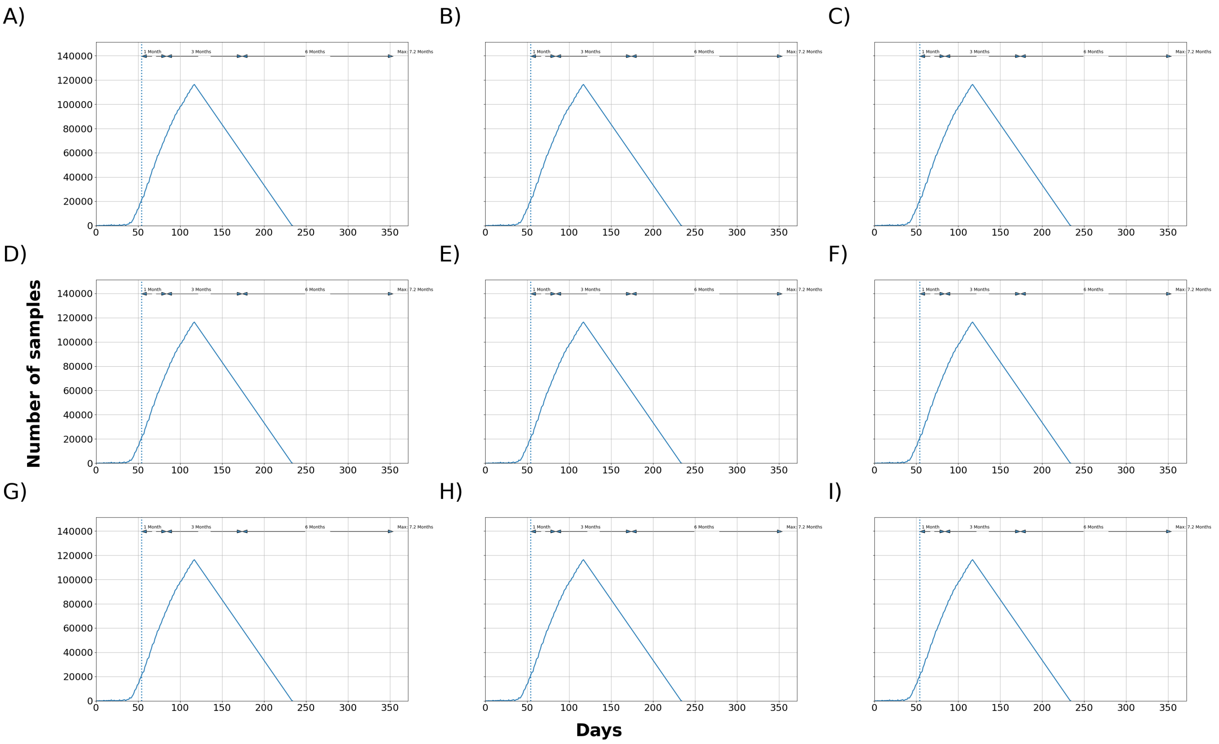


Figure 31: Maximum total number of samples waiting to be processed at the laboratory. 0 hours downtime and pool of 10 blood samples. SCPF = Sampler collectors per farm, TSC = Trained sample collectors. A) 5 SCPF, No Limit, 12 Hrs of RWS, B) 5 SCPF, 2000 TSC, 12 Hrs of RWS, C) 5 SCPF, 1750 TSC, 12 Hrs of RWS, D) 5 SCPF, 1000 TSC, 12 Hrs of RWS, E) 3 SCPF, No Limit, 12 Hrs of RWS, F) 5 SCPF, No Limit, 8 Hrs of RWS, G) 5 SCPF, 1250 TSC, 12 Hrs of RWS, H) 5 SCPF, 1500 TSC, 12 Hrs of RWS, I) 5 SCPF, 500 TSC, 12 Hrs of RWS.

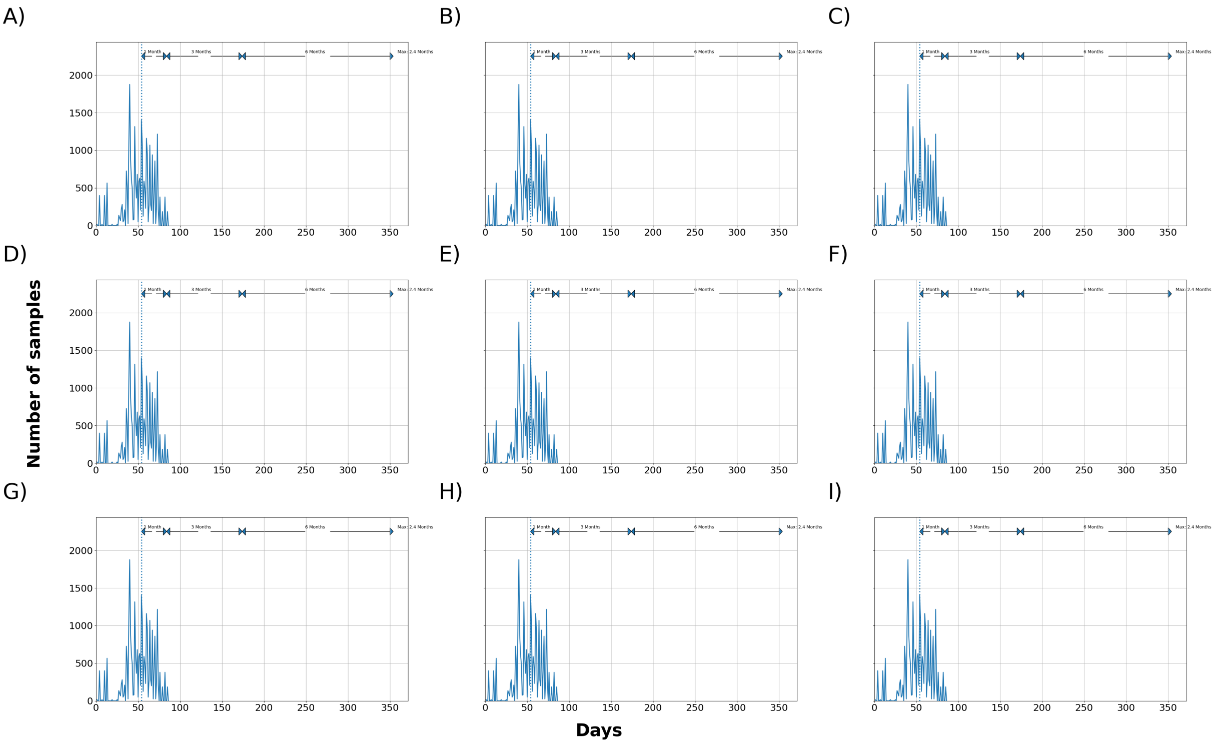


Figure 32: Median total number of samples waiting to be processed at the laboratory. 0 hours downtime and pool of 1 oral fluid sample. SCPF = Sampler collectors per farm, TSC = Trained sample collectors. A) 5 SCPF, No Limit, 12 Hrs of RWS, B) 5 SCPF, 2000 TSC, 12 Hrs of RWS, C) 5 SCPF, 1750 TSC, 12 Hrs of RWS, D) 5 SCPF, 1000 TSC, 12 Hrs of RWS, E) 3 SCPF, No Limit, 12 Hrs of RWS, F) 5 SCPF, No Limit, 8 Hrs of RWS, G) 5 SCPF, 1250 TSC, 12 Hrs of RWS, H) 5 SCPF, 1500 TSC, 12 Hrs of RWS, I) 5 SCPF, 500 TSC, 12 Hrs of RWS.

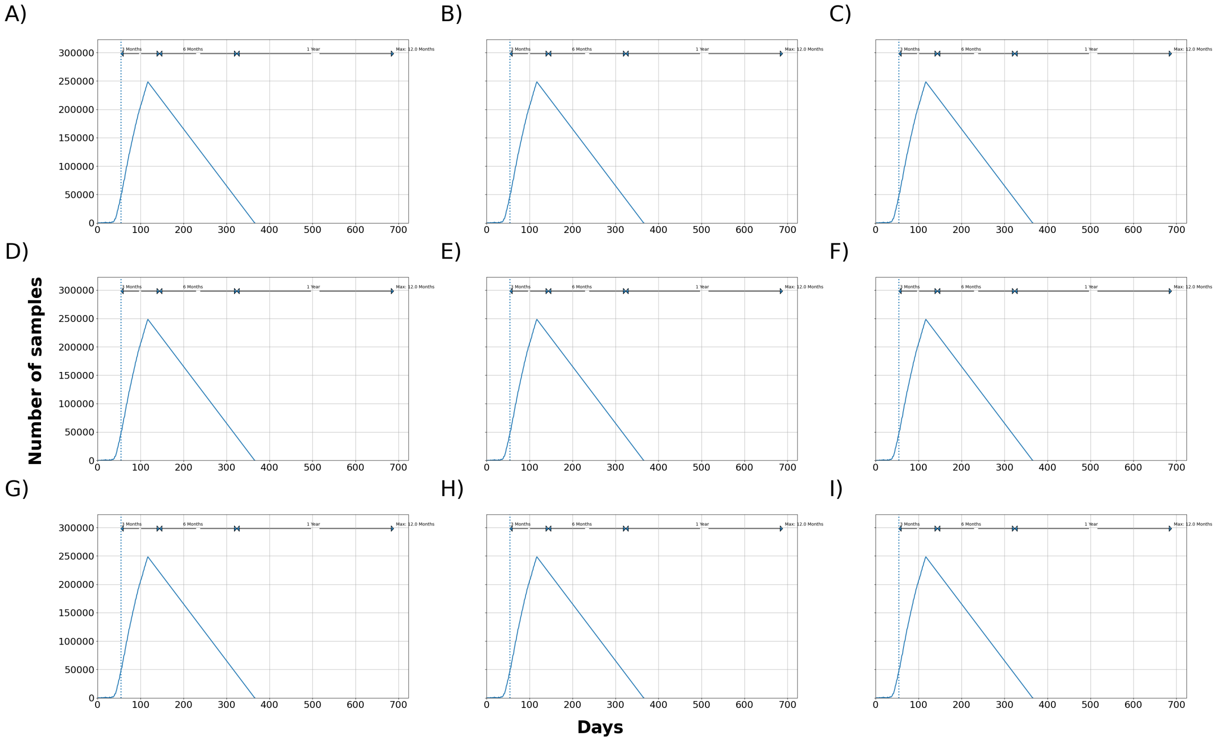


Figure 33: Maximum total number of samples waiting to be processed at the laboratory. 0 hours downtime and pool of 1 oral fluid sample. SCPF = Sampler collectors per farm, TSC = Trained sample collectors. A) 5 SCPF, No Limit, 12 Hrs of RWS, B) 5 SCPF, 2000 TSC, 12 Hrs of RWS, C) 5 SCPF, 1750 TSC, 12 Hrs of RWS, D) 5 SCPF, 1000 TSC, 12 Hrs of RWS, E) 3 SCPF, No Limit, 12 Hrs of RWS, F) 5 SCPF, No Limit, 8 Hrs of RWS, G) 5 SCPF, 1250 TSC, 12 Hrs of RWS, H) 5 SCPF, 1500 TSC, 12 Hrs of RWS, I) 5 SCPF, 500 TSC, 12 Hrs of RWS.

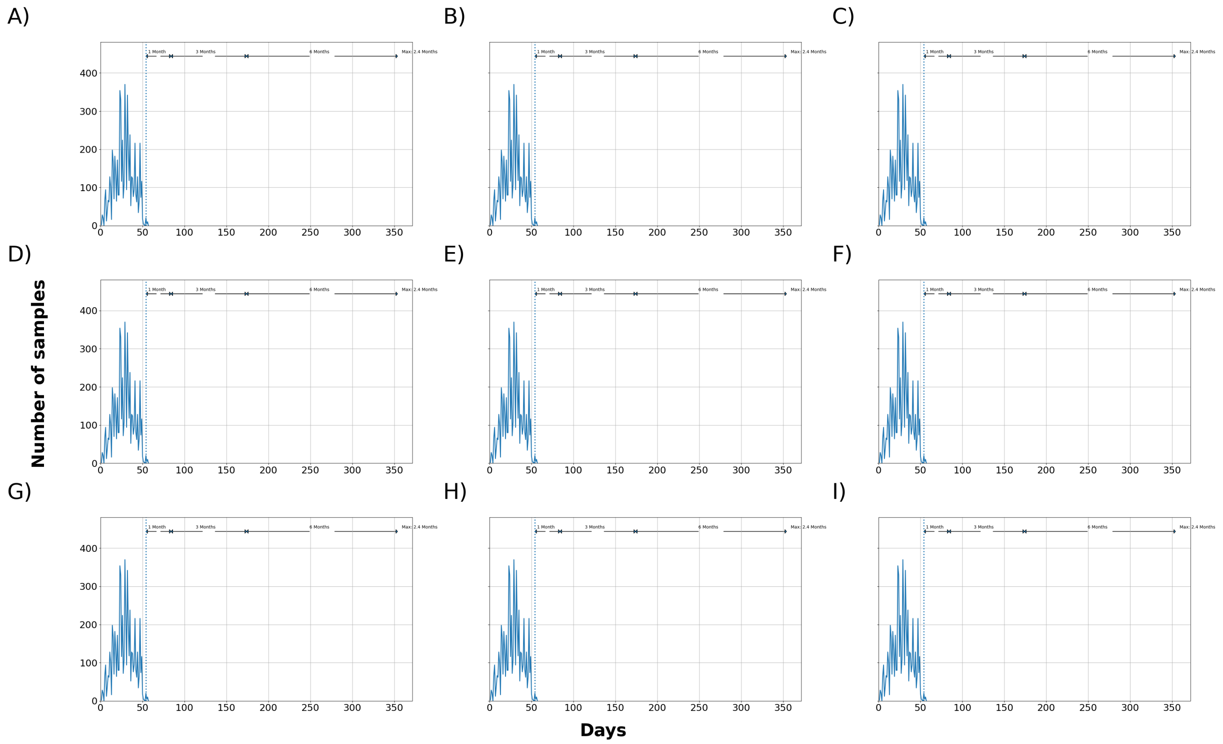


Figure 34: Median total number of samples waiting to be processed at the laboratory. 0 hours downtime and pool of 5 oral fluid samples. SCPF = Sampler collectors per farm, TSC = Trained sample collectors. A) 5 SCPF, No Limit, 12 Hrs of RWS, B) 5 SCPF, 2000 TSC, 12 Hrs of RWS, C) 5 SCPF, 1750 TSC, 12 Hrs of RWS, D) 5 SCPF, 1000 TSC, 12 Hrs of RWS, E) 3 SCPF, No Limit, 12 Hrs of RWS, F) 5 SCPF, No Limit, 8 Hrs of RWS, G) 5 SCPF, 1250 TSC, 12 Hrs of RWS, H) 5 SCPF, 1500 TSC, 12 Hrs of RWS, I) 5 SCPF, 500 TSC, 12 Hrs of RWS.

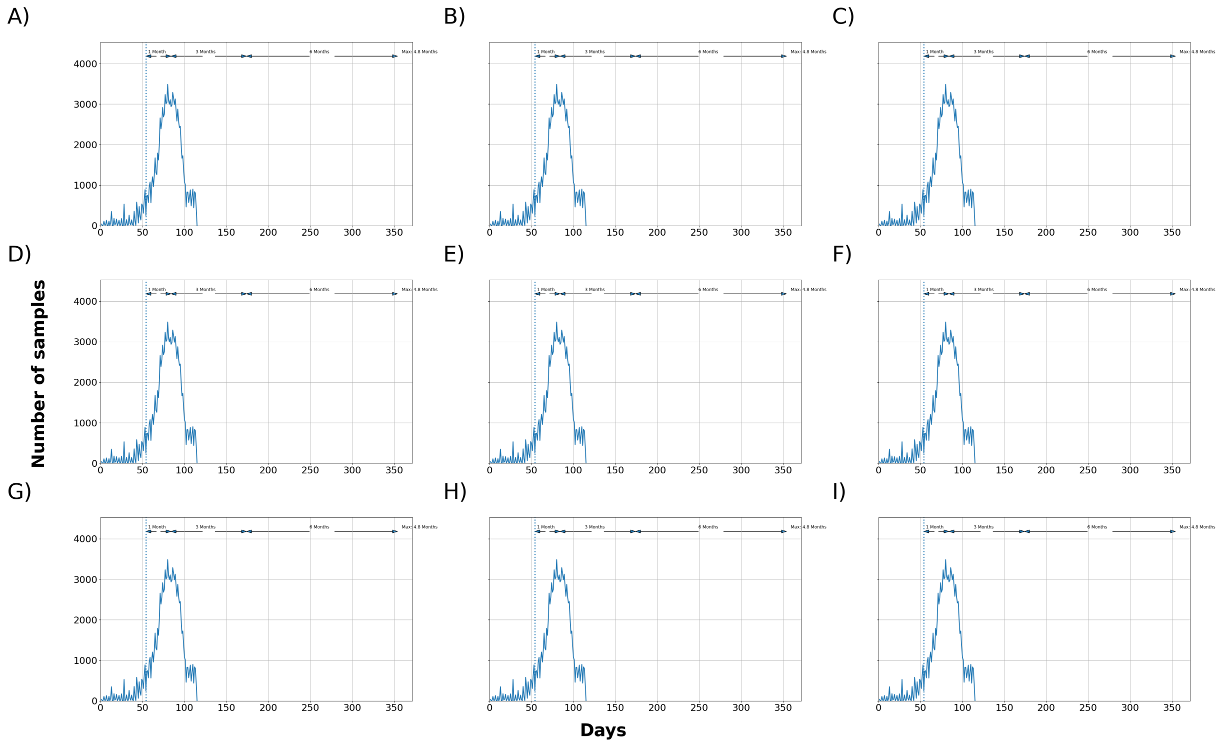


Figure 35: Maximum total number of samples waiting to be processed at the laboratory. 0 hours downtime and pool of 5 oral fluid samples. SCPF = Sampler collectors per farm, TSC = Trained sample collectors. A) 5 SCPF, No Limit, 12 Hrs of RWS, B) 5 SCPF, 2000 TSC, 12 Hrs of RWS, C) 5 SCPF, 1750 TSC, 12 Hrs of RWS, D) 5 SCPF, 1000 TSC, 12 Hrs of RWS, E) 3 SCPF, No Limit, 12 Hrs of RWS, F) 5 SCPF, No Limit, 8 Hrs of RWS, G) 5 SCPF, 1250 TSC, 12 Hrs of RWS, H) 5 SCPF, 1500 TSC, 12 Hrs of RWS, I) 5 SCPF, 500 TSC, 12 Hrs of RWS.

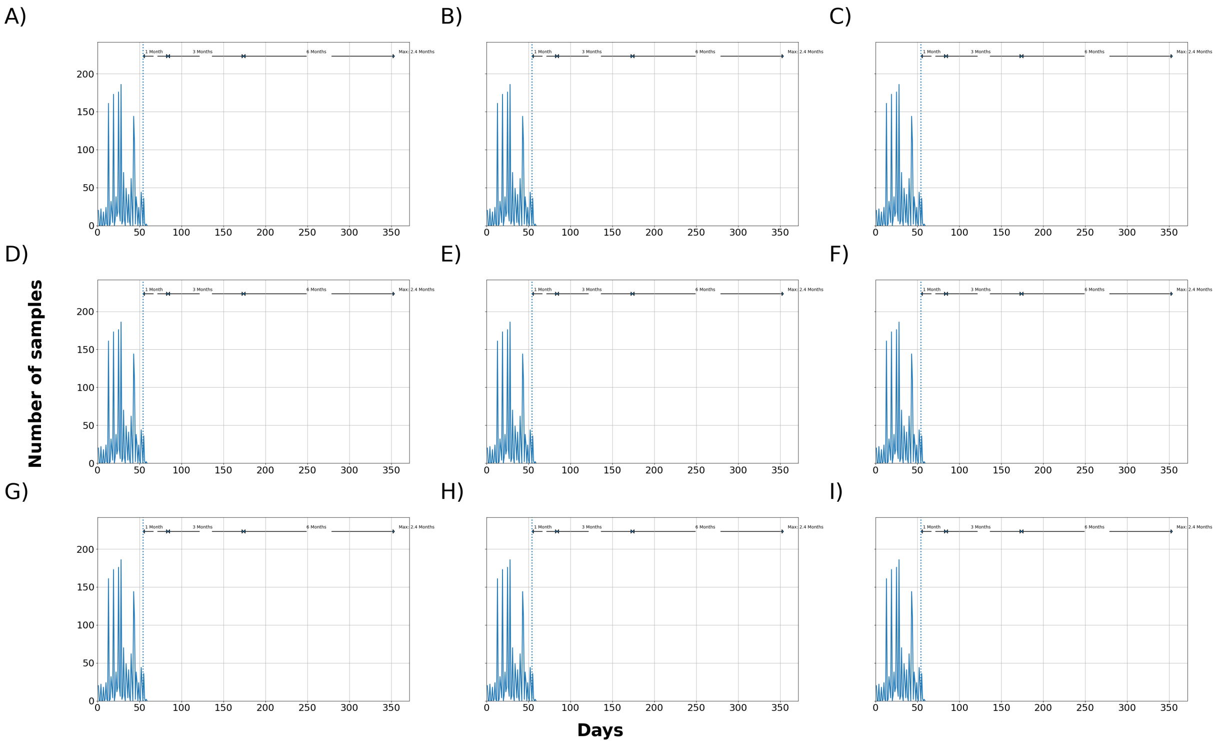


Figure 36: Median total number of samples waiting to be processed at the laboratory. 0 hours downtime and pool of 10 oral fluid samples. SCPF = Sampler collectors per farm, TSC = Trained sample collectors. A) 5 SCPF, No Limit, 12 Hrs of RWS, B) 5 SCPF, 2000 TSC, 12 Hrs of RWS, C) 5 SCPF, 1750 TSC, 12 Hrs of RWS, D) 5 SCPF, 1000 TSC, 12 Hrs of RWS, E) 3 SCPF, No Limit, 12 Hrs of RWS, F) 5 SCPF, No Limit, 8 Hrs of RWS, G) 5 SCPF, 1250 TSC, 12 Hrs of RWS, H) 5 SCPF, 1500 TSC, 12 Hrs of RWS, I) 5 SCPF, 500 TSC, 12 Hrs of RWS.

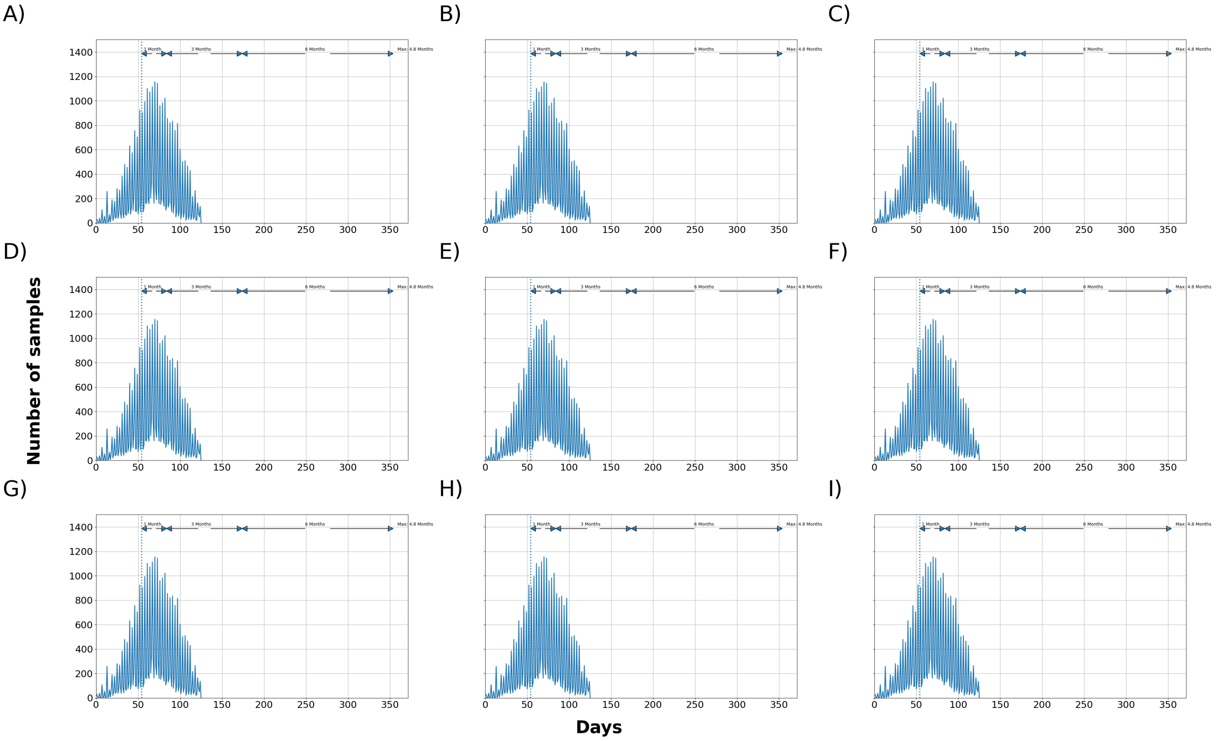


Figure 37: Maximum total number of samples waiting to be processed at the laboratory. 0 hours downtime and pool of 10 oral fluid samples. SCPF = Sampler collectors per farm, TSC = Trained sample collectors. A) 5 SCPF, No Limit, 12 Hrs of RWS, B) 5 SCPF, 2000 TSC, 12 Hrs of RWS, C) 5 SCPF, 1750 TSC, 12 Hrs of RWS, D) 5 SCPF, 1000 TSC, 12 Hrs of RWS, E) 3 SCPF, No Limit, 12 Hrs of RWS, F) 5 SCPF, No Limit, 8 Hrs of RWS, G) 5 SCPF, 1250 TSC, 12 Hrs of RWS, H) 5 SCPF, 1500 TSC, 12 Hrs of RWS, I) 5 SCPF, 500 TSC, 12 Hrs of RWS.

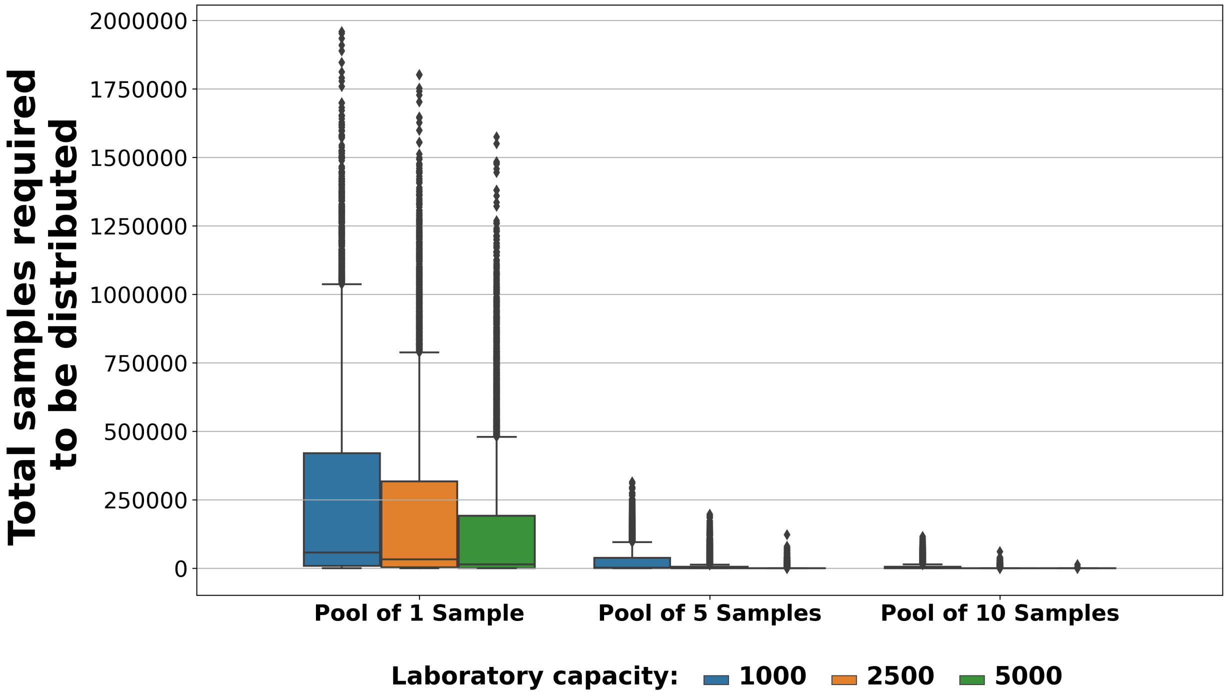


Figure 38: Total number of samples required to be distributed to other laboratories in a sampling scenario using 72 hours downtime with 3 SCPF, 12 RWS and no limit on TSC. SCPF = Sampler collectors per farm, TSC = Trained sample collectors, RWS = Regular working schedule.

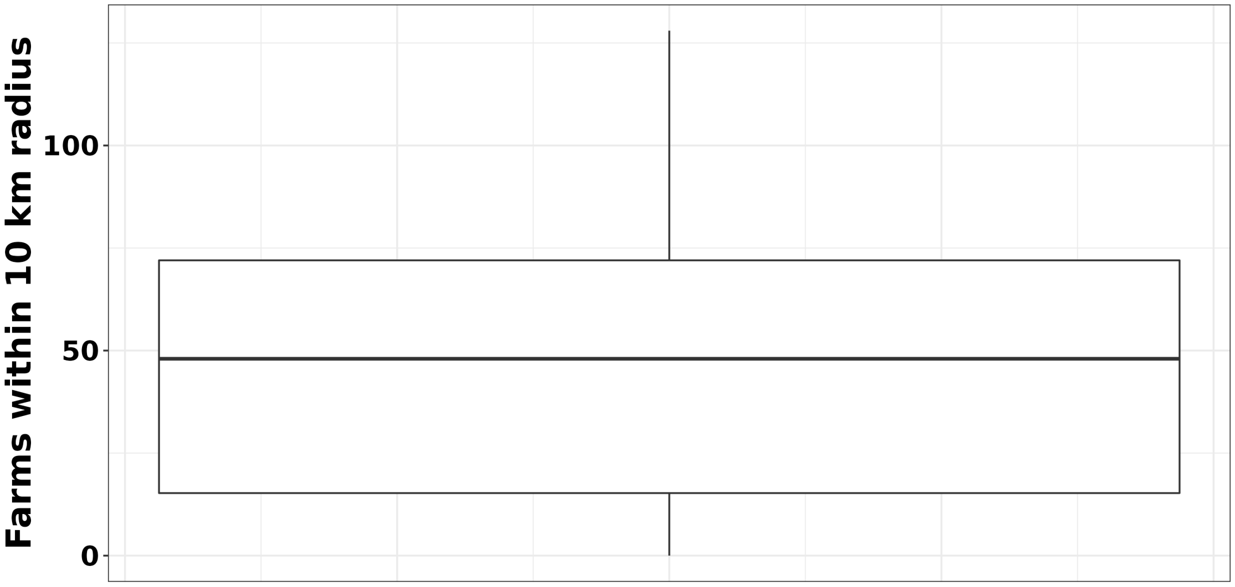


Figure 39: Farm neighbor distribution within 10 kilometers radius.

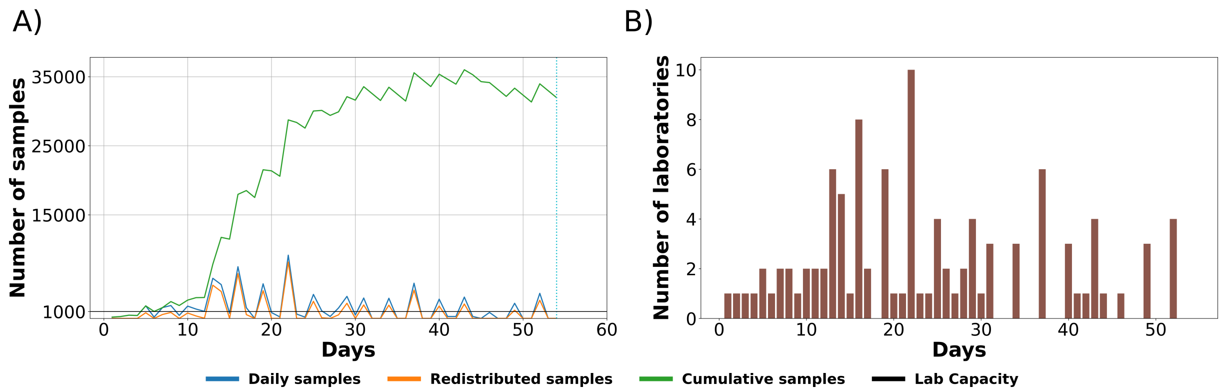


Figure 40: A) The daily median number of samples processed by the laboratory (blue line), distributed to other NAHLN laboratories upon reaching maximum processing capacity (orange line), or accumulated in a laboratory if not redistributed (green line), and B) daily number of NAHLN laboratories required to process the samples. This analysis involves a sampling scenario using 72 hours downtime with 3 SCPF, 12 RWS and no limit on TSC. SCPF = Sampler collectors per farm, TSC = Trained sample collectors, RWS = Regular working schedule.

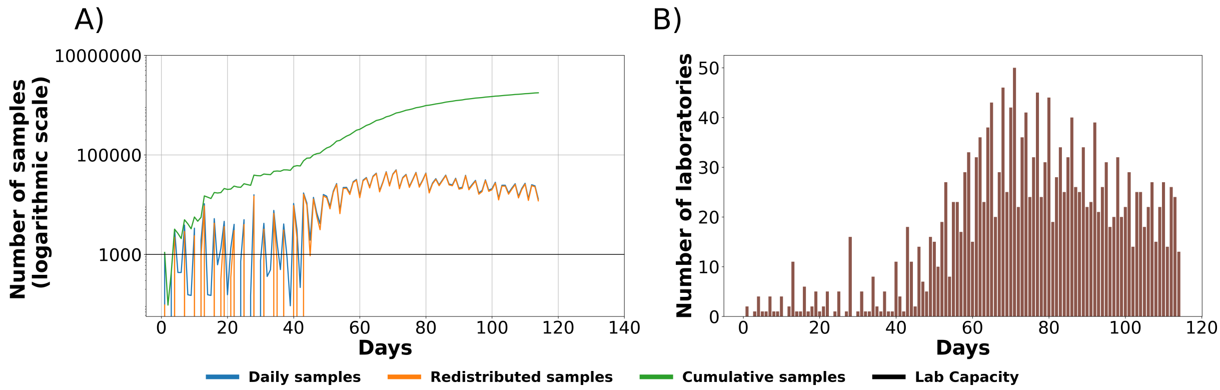


Figure 41: The daily maximum number of samples processed by the laboratory (blue line), distributed to other NAHLN laboratories upon reaching maximum processing capacity (orange line), or accumulated in a laboratory if not redistributed (green line), and B) daily number of NAHLN laboratories required to process the samples. This analysis involves a sampling scenario using 72 hours downtime with 3 SCPF, 12 RWS and no limit on TSC. SCPF = Sampler collectors per farm, TSC = Trained sample collectors, RWS = Regular working schedule.

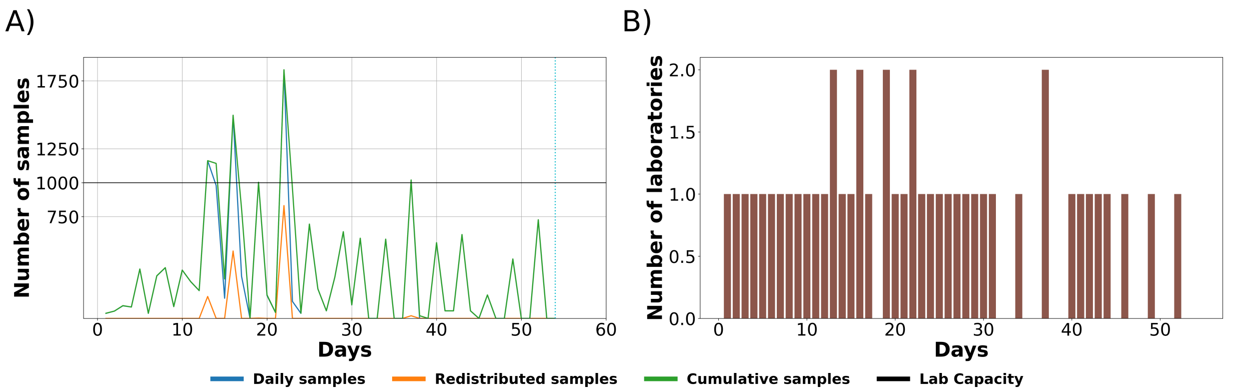


Figure 42: A) The daily median number of samples processed in pools of five (blue line), distributed to other NAHLN laboratories upon reaching maximum processing capacity (orange line), or accumulated in a laboratory if not redistributed (green line), and B) daily number of NAHLN laboratories required to process the samples. This analysis involves a sampling scenario using 72 hours downtime with 3 SCPF, 12 RWS and no limit on TSC. SCPF = Sampler collectors per farm, TSC = Trained sample collectors, RWS = Regular working schedule.

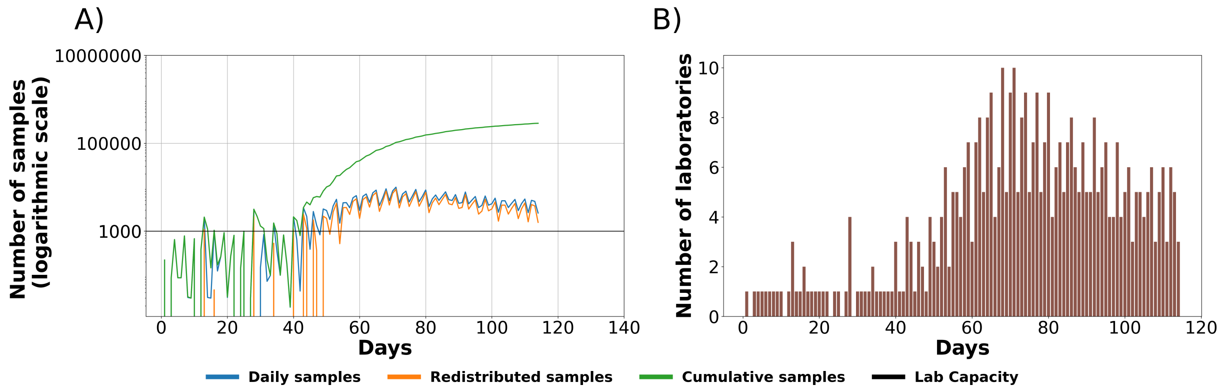


Figure 43: A) The daily maximum number of samples processed in pools of five (blue line), distributed to other NAHLN laboratories upon reaching maximum processing capacity (orange line), or accumulated in a laboratory if not redistributed (green line), and B) daily number of NAHLN laboratories required to process the samples. This analysis involves a sampling scenario using 72 hours downtime with 3 SCPF, 12 RWS and no limit on TSC. SCPF = Sampler collectors per farm, TSC = Trained sample collectors, RWS = Regular working schedule.

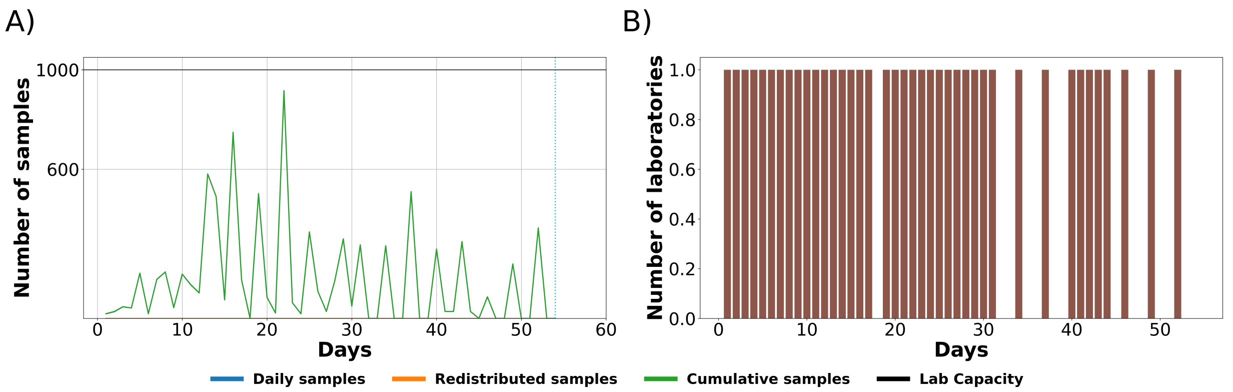


Figure 44: A) The daily median number of samples processed in pools of 10 (blue line), distributed to other NAHLN laboratories upon reaching maximum processing capacity (orange line), or accumulated in a laboratory if not redistributed (green line), and B) daily number of NAHLN laboratories required to process the samples. This analysis involves a sampling scenario using 72 hours downtime with 3 SCPF, 12 RWS and no limit on TSC. SCPF = Sampler collectors per farm, TSC = Trained sample collectors, RWS = Regular working schedule.

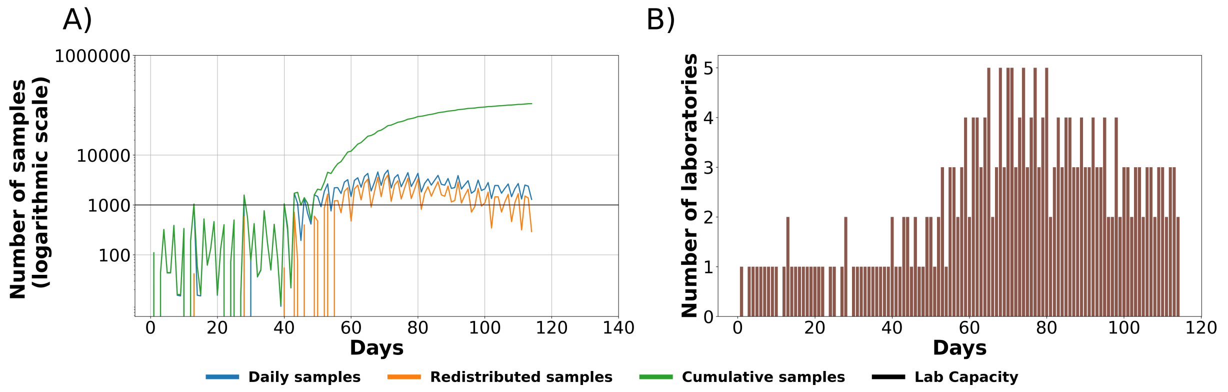


Figure 45: A) The daily maximum number of samples processed in pools of 10 (blue line), distributed to other NAHLN laboratories upon reaching maximum processing capacity (orange line), or accumulated in a laboratory if not redistributed (green line), and B) daily number of NAHLN laboratories required to process the samples. This analysis involves a sampling scenario using 72 hours downtime with 3 SCPF, 12 RWS and no limit on TSC. SCPF = Sampler collectors per farm, TSC = Trained sample collectors, RWS = Regular working schedule.
